# Unified classification of mouse retinal ganglion cells using function, morphology, and gene expression

**DOI:** 10.1101/2021.06.10.447922

**Authors:** Jillian Goetz, Zachary F. Jessen, Anne Jacobi, Adam Mani, Sam Cooler, Devon Greer, Sabah Kadri, Jeremy Segal, Karthik Shekhar, Joshua Sanes, Gregory W. Schwartz

**Affiliations:** Department of Ophthalmology, Feinberg School of Medicine, Northwestern University, Chicago, IL, USA; Northwestern University Interdepartmental Neuroscience Program, Northwestern University, Evanston, IL, USA; Medical Scientist Training Program, Northwestern University, Chicago, IL, USA; F.M. Kirby Neurobiology Center, Department of Neurology, Boston Children’s Hospital, Harvard Medical School, Boston, MA 02115, USA; Center for Brain Science and Department of Molecular and Cellular Biology, Harvard University, Cambridge, MA, USA; Current address: Department of Neuroscience, Brown University, Providence, RI, USA; Current address: Department of Neurosurgery, Stanford University, Stanford, CA, USA; Current address: Department of Pathology and Department of Preventive Medicine, Feinberg School of Medicine, Northwestern University, Chicago, IL, USA; Department of Pathology, Pritzker School of Medicine, University of Chicago, Chicago, IL, USA; Department of Chemical and Biomolecular Engineering, University of California Berkeley, Berkeley, CA, USA; Department of Physiology, Feinberg School of Medicine, Northwestern University, Chicago, IL, USA; Department of Neurobiology, Weinberg College of Arts and Sciences, Northwestern University, Evanston, IL, USA

## Abstract

Classification and characterization of neuronal types are critical for understanding their function and dysfunction. Neuronal classification schemes typically rely on measurements of electrophysiological, morphological, and molecular features, but aligning such datasets has been challenging. Here, we present a unified classification of mouse retinal ganglion cells (RGCs), the sole retinal output neurons. We used visually-evoked responses to classify 1859 mouse RGCs into 42 types. We also obtained morphological or transcriptomic data from subsets and used these measurements to align the functional classification to publicly available morphological and transcriptomic data sets. We created an online database that allows users to browse or download the data and to classify RGCs from their light responses using a machine learning algorithm. This work provides a resource for studies of RGCs, their upstream circuits in the retina, and their projections in the brain, and establishes a framework for future efforts in neuronal classification and open data distribution.

## Introduction

A major goal in biology is the establishment of a comprehensive atlas of cell types. Many large-scale efforts are underway to classify cells in different tissues (BRAIN Initiative Cell Census Network (BICCN), 2021; Hodge et al., 2019; Regev et al., 2017; Wilbrey-Clark et al., 2020; Yuste et al., 2020). In the central nervous system (CNS), classification efforts have relied mainly on three types of information: functional, morphological, and molecular. Functional classification involves the physiological properties of neurons, typically measured by electrophysiological recordings. Morphological classification uses the dendritic and axonal structures of neurons, measured by light or electron microscopic (EM) methods. Molecular classification was initially based on immunohistochemical or *in situ* hybridization, but has more recently relied on gene expression patterns (transcriptomics) assessed by high-throughput single-cell RNAseq and spatial transcriptomics (Close et al., 2021; Yuste et al., 2020). It has become increasingly clear that different classification methods offer complementary information and that a comprehensive classification of cell types needs to unify all three modalities (BRAIN Initiative Cell Census Network (BICCN), 2021; Scala et al., 2020; Zeng and Sanes, 2017).

The mammalian retina is especially well suited to provide a template for integrating functional, morphological, and molecular classification for three reasons. First, many retinal cell types exhibit regular spacing, called a mosaic, which ensures smooth and complete sampling of visual space (Bleckert et al., 2014; Kay et al., 2012; Reese and Keeley, 2015; Rockhill et al., 2000; Rousso et al., 2016; Wässle et al., 1981). This property means that experimentalists can sample from a sub-region of the retina and be assured that they will find cells of each type. Moreover, mosaics establish an independent metric to assess whether a set of cells comprises an authentic type. Second, because the retina responds to light *ex vivo*, functional measurements of retinal neurons include both intrinsic biophysical properties and response properties to visual stimuli. Light responses depend on the entire upstream synaptic network, creating a rich dataset. Finally, our knowledge of the morphology of retinal neurons, particularly in the mouse, is unparalleled among tissues of the mammalian CNS (Bae et al., 2018; Hoon et al., 2014; Sanes and Masland, 2015).

Here, we present a unified functional, morphological, and genetic classification of mouse retinal ganglion cells (RGCs), the output cells of the retina. We collected detailed functional data from 1859 RGCs and also obtained morphological or transcriptomic data from subsets of these cells. We then used these doubly-characterized cells to align the functional classification with publicly available large-scale datasets of RGC morphology (381 RGCs reconstructed from EM sections; Bae et al., 2018) and gene expression (35,699 single-RGC transcriptomes; (Bae et al., 2018; Tran et al., 2019), thereby generating a unified atlas. Comparison of the three datasets reveals that close relationships between cell types identified by one criterion sometimes predicts close relationships by other criteria.

Finally, we provide two tools that make the data useful to the community and suggest formats for cross-modal analyses of other populations. First, we devised a machine learning classifier that allows researchers to infer an RGC’s functional type from a small and standardized set of spike measurements. Second, we curated the data in the form of a continuously updated, open-access library (rgctypes.org) where researchers can browse single-cell-or cell type-level data and download functional, morphological, and transcriptomic datasets.

## Results

RGCs have traditionally been classified by physiological, morphological, and molecular criteria. Recent studies have used high-throughput methods to categorize mouse RGCs at large scale using all three criteria: optical imaging of visually evoked responses (Baden et al., 2016); reconstruction from serial electron microscopic sections; (Bae et al., 2018); and transcriptomic analysis of single RGCs (Tran et al., 2019). Our goal was to unify these dimensions into a single schema that was as complete as possible in representing all known RGC types in the mouse. We made our measurements in one cell at a time, allowing us to perform online functional classification followed by recovery of the same cells for morphological or transcriptomic measurements (**Figure 1**).

**Figure 1.**
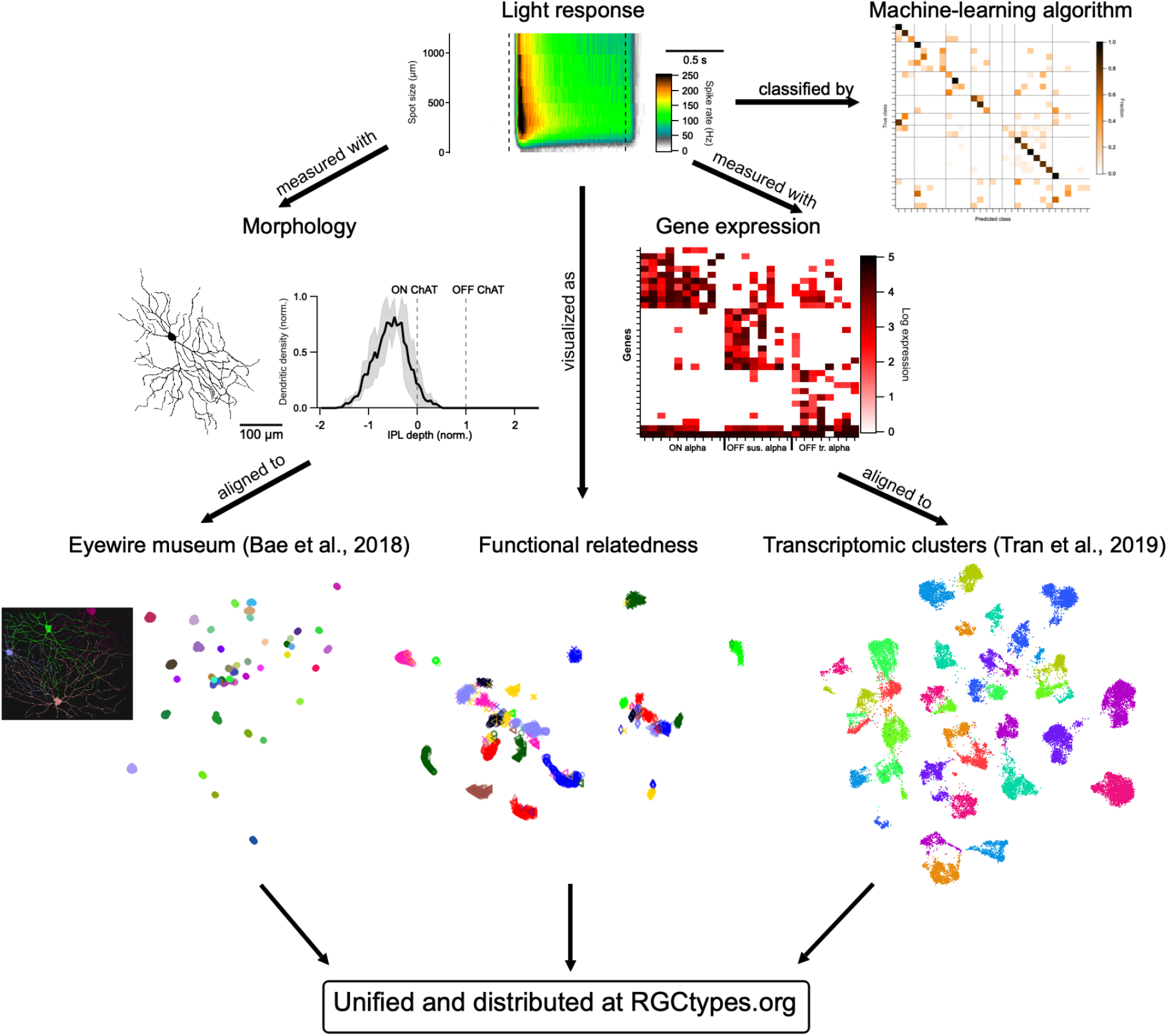
Schematic of approach. Light responses to a standard stimulus set were measured in 1859 RGCs with subsets measured for morphology or gene expression. Functional data on light responses was visualized using UMAP (see **Methods**) and classified with machine learning. Morphological and transcriptomic measurements were aligned to published datasets.

### Functional classification of RGCs

We began with physiological characterization, using a rapid and standardized light stimulus protocol for functional measurements. Experiments were performed in dark-adapted *ex vivo* preparations of the mouse retina where capacitive spikes from RGCs were recorded with cell-attached electrodes. Standard light stimuli presented to every RGC were rod-saturating (~200 isomerizations/rod/s) spots (λ = 450 nm) from darkness with diameters ranging from 30 to 1200 μm, centered on the receptive field (RF) of each individual cell. We presented additional stimuli to subsets of RGCs to test for specific forms of feature selectivity. Moving bars were used to test for direction selectivity (DS), flashed bars and drifting gratings for orientation selectivity (OS), and contrast series for contrast suppression (**Figure S1**). Background luminance values of 1000 R*/rod/s were used for drifting gratings and contrast series experiments were presented for less than 6 minutes per cell; response changes for subsequent measurements from darkness were negligible (data not shown).

Our standard stimulus paradigm differed from the full-field “checkerboard” white noise and “chirp” stimuli used in previous studies (Baden et al., 2016; Farrow and Masland, 2011; Jouty et al., 2018). Three considerations drove our stimulus choice. First, maintenance of a consistent light-adaptation state was essential because many aspects of RGC light responses change with luminance and with light adaptation (Tikidji-Hamburyan et al., 2015; Wienbar and Schwartz, 2018). High background light is unavoidable in functional two-photon imaging experiments due to excitation from the laser, limiting the period of stable light responses, especially in preparations lacking the retinal pigment epithelium (Euler et al., 2019). The use of patch electrodes allowed us to make measurements in darkness. Second, precise localization of stimuli with respect to the RF center cannot be achieved with full-field stimuli but turned out to be critical as shown below. Indeed, many RGCs that respond well to small stimuli in their RF center fail to respond to any full-field stimulus (Jacoby and Schwartz, 2017; Zhang et al., 2012). Finally, to facilitate standardization in the field, we wanted our stimulus to be simple and rapid and to correspond to those commonly used by others. For example, many previous studies have used RF-centered spots of different sizes, enabling retrospective comparisons (Jacoby and Schwartz, 2017; Johnson et al., 2018; Krieger et al., 2017; Marco et al., 2013; Rousso et al., 2016).

We assigned RGCs to 42 functional types by hand based on our iteratively updated understanding of their response patterns. 34 of these types were assigned based only on the responses to flashed spots, while the additional 8 types were subdivided by direction or orientation preference. Thus, while our classification is not free from human bias, two pieces of evidence, detailed below, strengthen our confidence that it represents an accurate typology: (1) cells that we placed in the same functional group typically had strong morphological and molecular similarities; and (2) a cross-validated algorithm successfully classified functional RGC types, including those with “external” classification data on which the algorithm was never trained. We organized the RGC types into 8 functional groups: ON sustained, OFF sustained, ON transient, OFF transient, ON OS, DS, ON-OFF small RF, and Suppressed-by-Contrast (SbC)/Other. These groups were chosen as a starting point based on previous work; a quantitative measure of functional relatedness is presented below.

In most recordings (1246/1859 cells), retinal orientation and cell locations were noted to determine whether classification varied based on retinal position. Response patterns within some RGC types have been shown to vary as a function of retinal position in photopic conditions (Joesch and Meister, 2016; Warwick et al., 2018), likely because of a pronounced cone opsin gradient along the dorsoventral axis (Nadal-Nicolás et al., 2020). In our dark-adapted preparation, however, where much of the light response was initiated in rods (Grimes et al., 2014), response variation across retinal position was minimal. We found no significant relationship between retinal position and any of the six response metrics we tested (see **Methods**). The responses of OFF transient alpha RGCs, which had previously been shown to depend on dorsoventral position in high luminance (Warwick et al., 2018) showed no position dependence under our conditions (data not shown).

Most functional RGC types were relatively uniformly distributed across retinal locations (**Figure S2**). A shuffle test revealed two RGC types with significant positional biases (**Table S1**). F-mini-ON RGCs were found in greater proportion in the ventral retina; however, we specifically targeted them in that region based on a previous report of their prevalence there in a transgenic line (Rousso et al., 2016). PixON RGCs were found in greater proportion in the dorsal retina which, to the best of our knowledge, does not represent sampling bias and has not been previously reported. Several other RGC position distributions showed trending biases that failed to meet the Bonferroni correction for multiple comparisons. These include the known prevalence of ON alpha RGCs in the temporal retina (Bleckert et al., 2014), and the previously unreported prevalence of UHD RGCs in the nasal retina.

Data from 37 RGC types are presented in three ways in **Figure 2** (only 37 of the 42 types are illustrated because DS RGCs with different directional preferences did not differ from each other in their responses to light spots). We first measured the response polarity and kinetics of light responses with a 200 μm light spot centered on the RF (marked “a” in the ON alpha RGC panel). This allowed us to assign cells according to response polarity (ON, OFF, ON-OFF or Suppressed-by-Contrast) and as having sustained or transient responses to luminance changes. Second, to assess how each RGC type’s response varied with stimulus size, we measured the total ON and OFF spike responses for spots of 12 sizes from well below the RF center diameter of the smallest RGC (30 μm) to a size that reached the far RF surround (1200 μm) (marked “b” in the ON alpha RGC panel). This information was critical in separating many types. For example, despite similar responses to the 200 μm spots, ON-OFF DS, HD1, HD2, and UHD all had different response profiles of their ON and OFF responses across spot size. For some RGC types, even the overall polarity of the light response depended on spot size. For example, HD2 RGCs are ON-OFF for small spots and ON for large spots (Jacoby and Schwartz, 2017) and the ON small OFF large RGC switches polarity entirely with spot size as its name suggests. Finally, we combined information about response amplitude and kinetics as a function of spot size into a single plot using a heatmap of firing rate over time for each spot size (marked “c” in the ON alpha RGC panel). Functional heatmaps of the variability in these responses within each functional group are presented in **Figure S3** and distributions of 6 common response metrics for each RGC type are shown in **Figure S4**.

**Figure 2.**
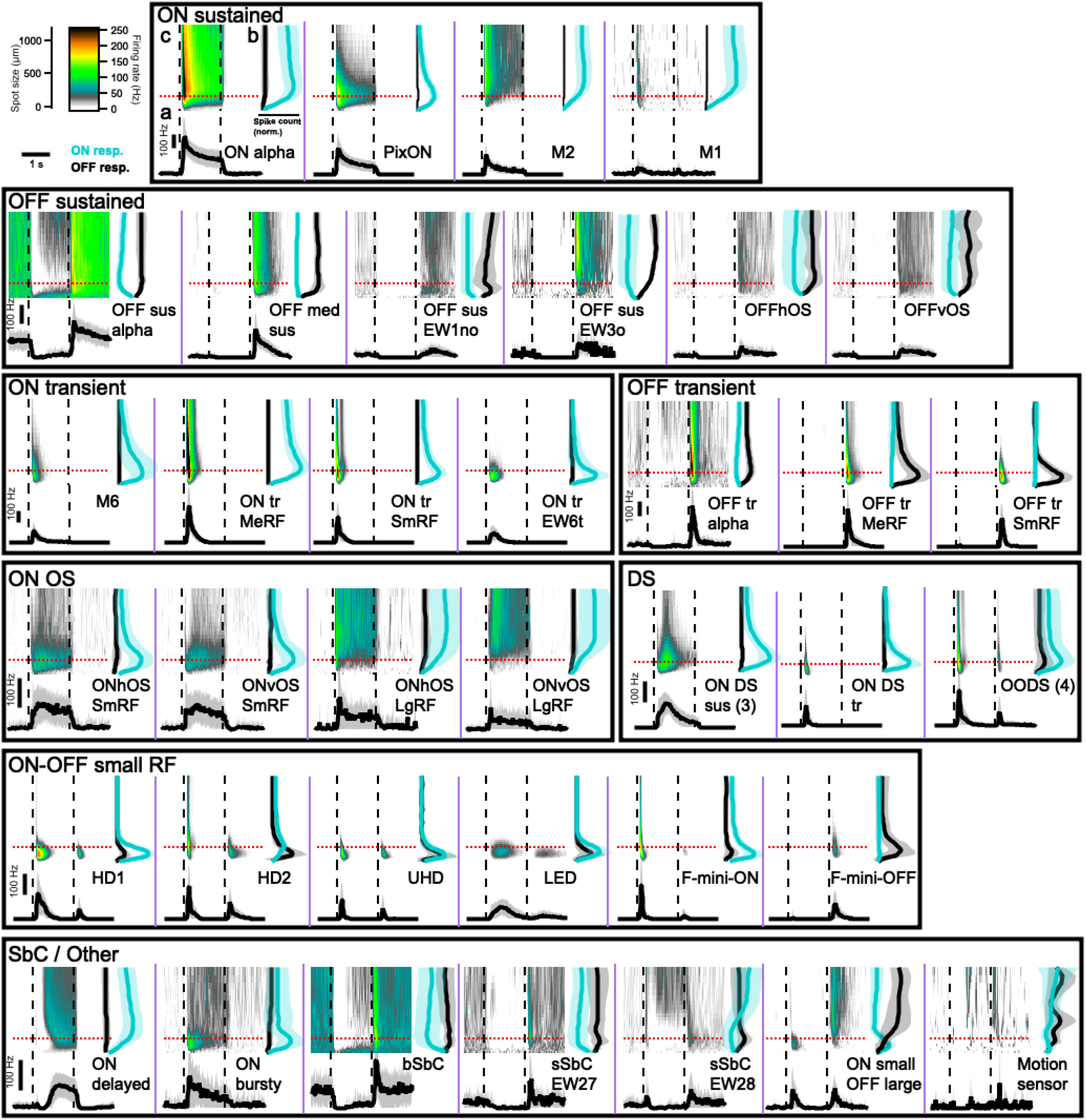
Functional diversity of mouse RGCs. Each panel (separated by purple lines) contains 3 graphs showing the light response of an RGC type to flashed spots of light (200 R*/rod/s) from darkness. The top left graph (marked ‘c’ in ON alpha panel) is a heatmap of average firing rate over time (x-axis) for spots from 30 – 1200 μm (y-axis). Dashed lines show the time of spot onset and offset. The top right graph (marked ‘b’ in ON alpha panel) shows the total spike count during flash onset (cyan) and offset (black) for each spot size. The solid lines indicate mean across cells and the shaded regions indicate standard deviation (s.d.). The bottom graph (marked ‘a’ in ON alpha panel) shows peristimulus time histogram (PSTH) plots averaging the response of each cell type to 200 μm spots, indicated in upper plots by red dotted lines. Scale bars in the upper left region are shared across all graphs. Separate scale bars for the y-axis of the PSTH plots are provided within each boxed group of cells and apply within that box. Abbreviations for cell types: sus = sustained; tr = transient; med = medium; EW = Eyewire (named based on the Eyewire museum); OS = orientation-selective; h = horizontal; v = vertical; DS = direction-selective; SmRF = small receptive field; MeRF = medium receptive field; LgRF = large receptive field; HD = high definition; UHD = ultra high definition; LED = local edge detector; (b,s)SbC = (bursty, sustained) suppressed-by-contrast.

### Functional relatedness of RGC types

To visualize the relationships between functional RGC types, we used uniform manifold approximation and projection (UMAP) (Becht et al., 2018; McInnes et al., 2018) (**Figure 3**). The UMAP algorithm assigned each cell to a point in 2D space based only on its response to spots of varying size (the data in **Figure 2**) with closely-related cells projecting to nearby locations in this space. We did not include the moving bar or drifting gratings responses as input to the UMAP algorithm since they were not measured for every RGC. Therefore, DS RGCs with different direction preferences and OS RGCs with different orientation preferences were grouped together in this representation. Most RGC types formed clear clusters in UMAP space with a few exceptions, typically for types that were sampled sparsely in our dataset (**Figure 3A**).

**Figure 3.**
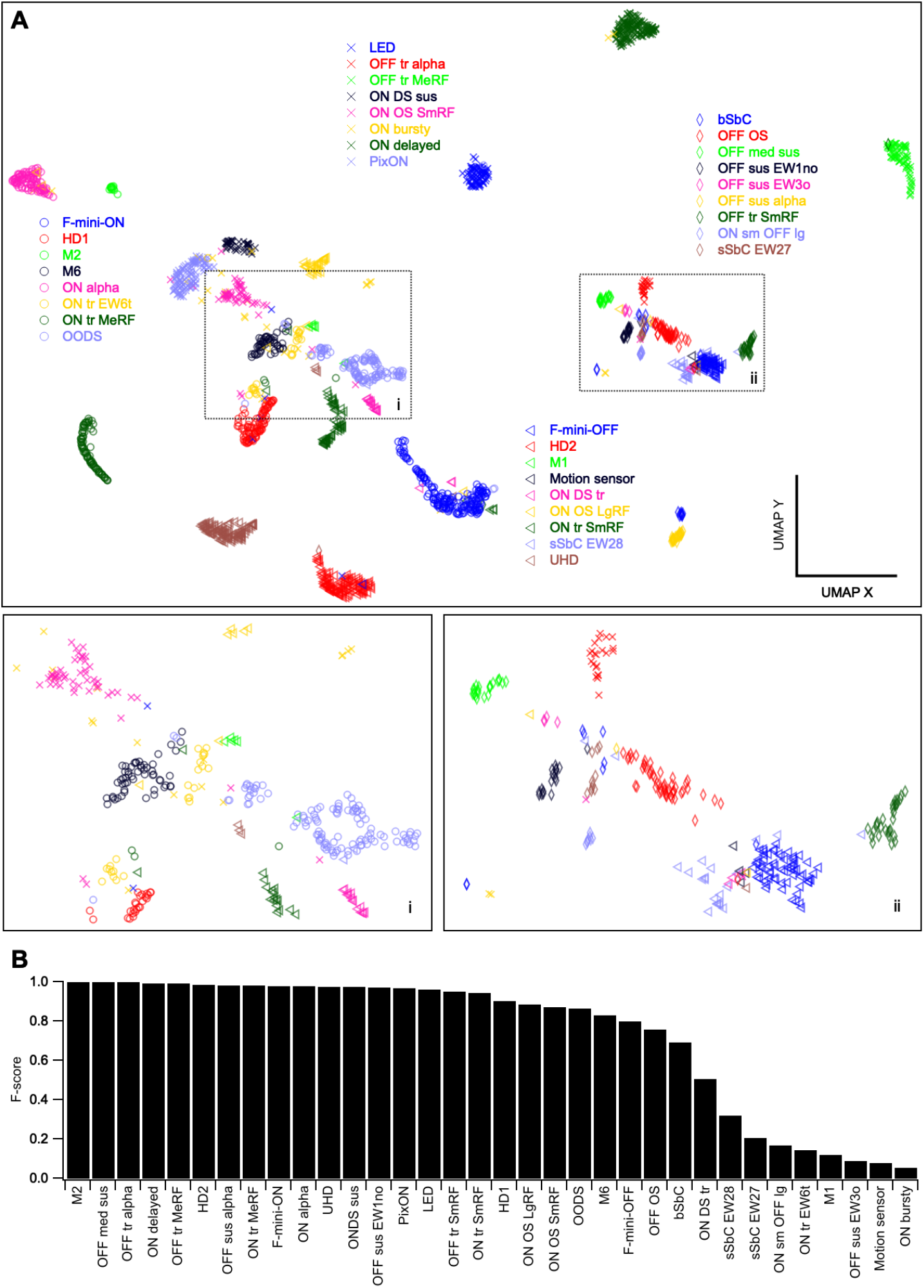
Visualization of functional relationships among RGCs. (A) UMAP projection of 1859 RGCs labeled by assigned functional type. Insets show magnified views of boxed regions. (B) F-score for each RGC type, the harmonic mean of the precision (fraction of a given cluster representing a single labeld type) and recall (fraction of our labeled cells of a given type in a single cluster) of its identification within a single DBSCAN cluster.

To assess the clustering of each of our defined functional types in this UMAP space, we subsequently clustered points in this 2D space with DBSCAN (Ester et al., 1996). F-scores, which measure the overlap between our 34 type labels and the 33 clusters identified by DBSCAN are shown in **Figure 3B**. These scores represent the degree of functional similarity (for this stimulus paradigm) within our assigned types relative to the differences between types. The types with lowest F-scores (10 types < 0.75) likely contain the majority of our labeling errors. There are also RGC types in this group (e.g. M1 and ON bursty RGCs) for which additional lines of evidence suggest that our labels are correct (see morphological and molecular data below), but for which the average spike rates for flashed spots alone are not sufficient for highly reliable functional classification. The average F-score, weighted by the number of cells of each type in the dataset, was 0.89.

### Automated functional classification

We implemented a machine learning classifier to assign RGCs to types based on a feature set comprising spike responses to spots of varying size. Since the responses to moving bars and drifting gratings were not included in the feature set, we collapsed DS and OS cells across direction and orientation, respectively. Our dataset of 1859 RGCs across 34 types, was split into a training set (n = 883), a calibration set (n = 500), and a test set for model evaluation (n = 476). Details of data split and classifier architecture are provided in **Methods**.

Following training, the performance of the classifier was evaluated on the test set (Figure 3). For each cell, the classifier outputs the probability of membership in each RGC type. Thus, the algorithm provides both a “best guess” and a confidence rating for each prediction. An advantage of probabilistic scoring is that the classifier predictions can easily be updated to include complementary sources of information (e.g. prior probabilities based on stratification depth in the inner plexiform layer, IPL, or labeling in a transgenic line) via Bayes’ rule (MacKay and Mac, 2003). Without thresholding the probability scores, classification accuracy was 59% overall (**Figures 4B,D**). The correct RGC type was among the top three choices of the classifier 75% of the time (**Figure 4A, inset**), suggesting that additional information (functional, structural or molecular), could be used to refine its predictions.

**Figure 4.**
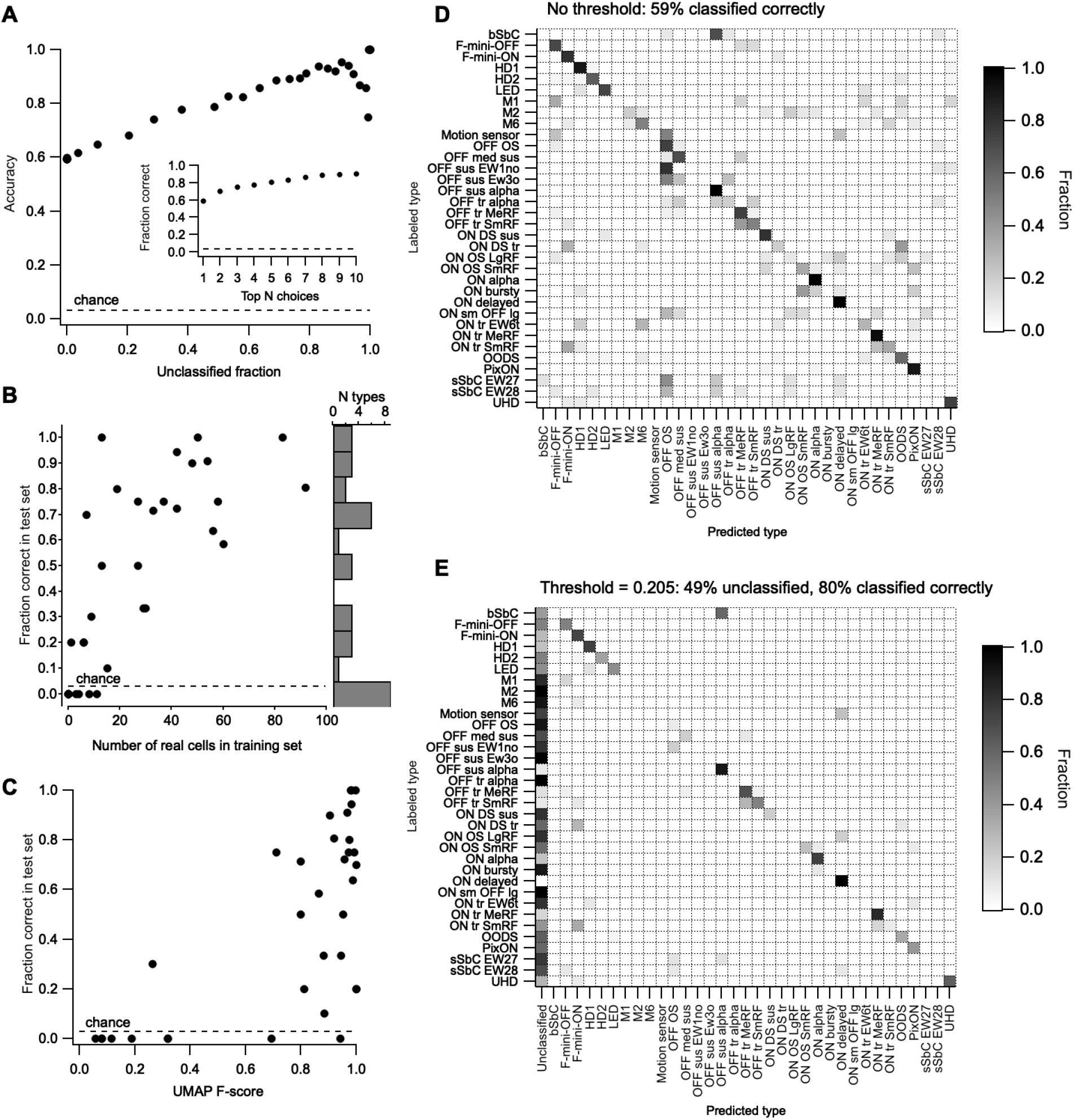
Functional classification from spot responses. (A) Overall model accuracy (y-axis) as a function of the fraction of unclassified cells in the test cells (x-axis), which increases with the classification margin. The dashed line represents the expected accuracy of a random classifier. *Inset*, fraction of instances when the correct choice was present among the top 1-10 probability scores in the classifier output. (B) Fraction of test cells of each type classified correctly versus the number of cells of that type in the training set. Histogram at the right shows the distribution of classifier accuracy across RGC types. (C) Accuracy of classification for each RGC type versus its F-score from Fig. 3B. (D) Confusion matrix (row normalized) for the classifier with no explicit classification margin set. Dotted lines separate RGC groups as in Figure 2. (E) Confusion matrix (row normalized) for the classifier with a classification margin of 0.205. The fraction of unclassified cells of each type is shown in the first column. Remaining entries in the matrix only consider classified cells.

To gain some measure of the degree to which RGC types classified by flashed spots correspond to those identified by other criteria, we used 5 types of “external” validation data that were not available to the classifier: (1) an image of the cell’s morphology, (2) fluorescent labeling in one of several transgenic lines in which small subsets of RGCs are labeled (see **Methods**), (3) synaptic currents in voltage-clamp whose profile match those in our published work on particular RGC types (Cooler and Schwartz, 2020; Jacoby and Schwartz, 2017; Jacoby et al., 2015; Mani and Schwartz, 2017; Nath and Schwartz, 2016, 2017), (4) large soma size noted at the time of recording (for the 3 alpha RGC types), (5) DS or OS as measured by moving bars and/or drifting gratings (see **Figure S1**). 634 of the RGCs in our dataset (34%) were validated by one or more of these external data types (**Table S2**). Classifier performance was slightly better for the validated cells in our test set (65% correct, n = 221 cells) than for the unvalidated cells (59% correct, n = 255 cells).

Classification accuracy varied widely across RGC types with 9 types having 0% sensitivity and the other 24 having a median accuracy of 71% (**Figure 4B**). Overall accuracy scaled linearly with unclassified fraction as we increased the classification margin, i.e. minimum probability score at which cells are assigned a type label (**Figure 4A**). Cells with maximal class probabilities below the classification margin are considered “unclassified”. Increasing the classification margin to 0.205 achieved an accuracy of 80% across the whole data set with 49% of cells unclassified (**Figure 4E**). The most significant limitation of our classifier was the size of the training set (**Figure 4C**). Thus, we expect classifier performance to improve steadily as we continue to collect more data, particularly from rare RGC types. Updated results, newly trained versions, and tutorials for formatting data and running it through the classifier will continue to be made available at rgctypes.org.

To our knowledge, this represents the first automated functional classifier designed to work on the full population of RGCs. While the overall performance might seem modest, the scale of the problem is beyond most attempts at supervised neuronal classification in the CNS. RGCs have many functional similarities, particularly when only probed with a single stimulus type, and we were attempting to classify them into 34 types, some with very little training data (in fact, 3 with none at all, guaranteeing failure: M1, Motion sensor, OFF sus EW3o). As shown next, morphological and molecular data offer large amounts of complementary information for RGC classification, so we expect a future multi-modal classifier to achieve much higher performance.

### Alignment of functional and morphological classification

After we recorded visually evoked responses from RGCs, we filled some of them with either AlexaFluor 488 for live imaging or with Neurobiotin for post-hoc imaging. 136 of these images could be effectively computationally flattened and registered to the choline acetyltransferase (ChAT) bands; ChAT is an established marker for the dendrites of starburst amacrine cells, which stratify in stereotypical, narrow strata (Sümbül et al., 2014). This alignment allowed quantitative measurements of *en face* morphology and stratification patterns within the IPL (**Figure 5**).

**Figure 5.**
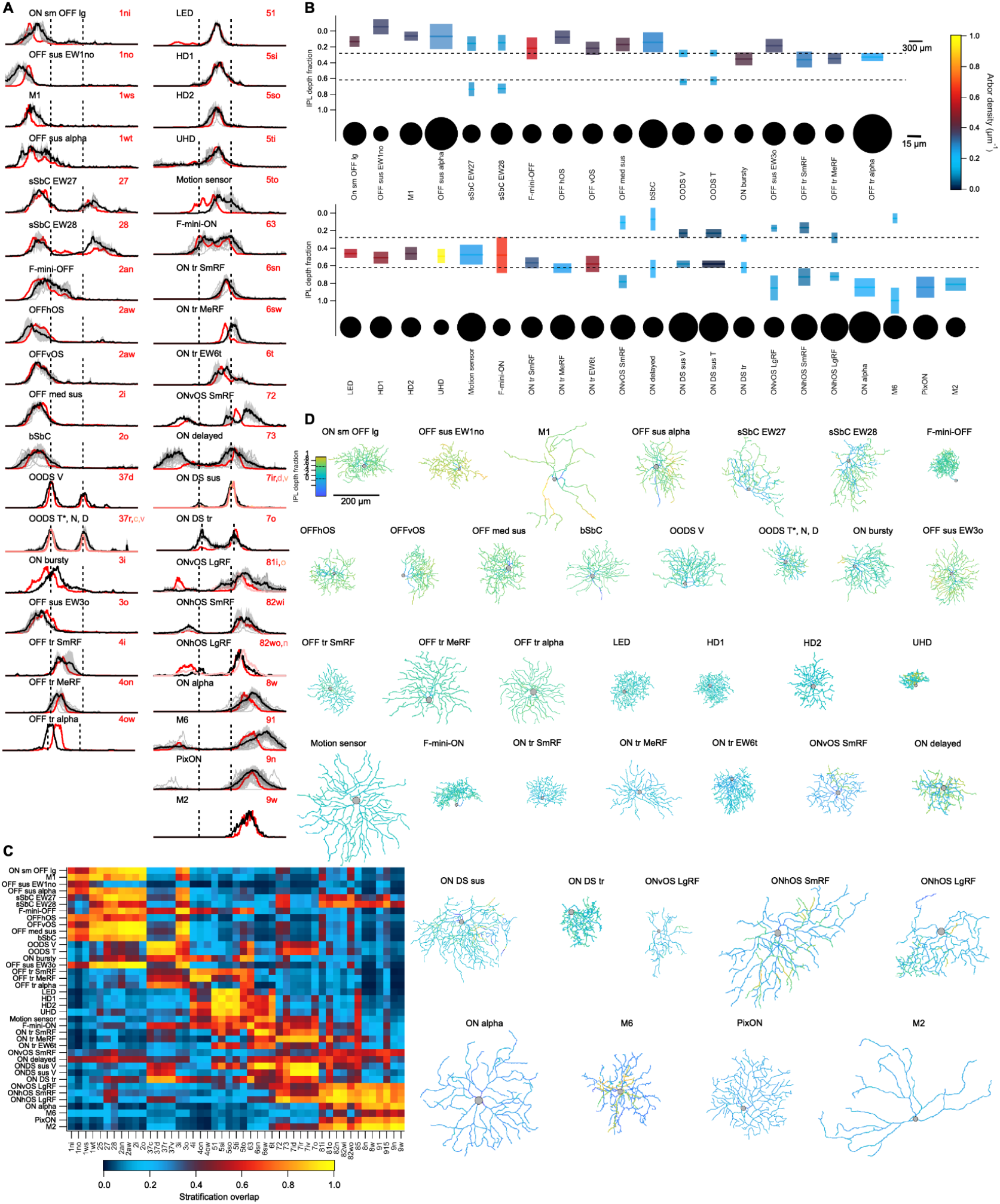
Morphological diversity of mouse RGCs. (A) Stratification profile of each RGC type along the depth of the IPL from its outer (left) to inner (right) limits. Dashed lines indicate ChAT bands. Profiles include individual cells (*thin gray lines*), the mean (*thick black line*), and s.d. (*gray shading*) as well ab ns the presumed matching type(s) in the Eyewire museum (*shades of red*). (B) Summary plot of the morphology of each RGC type. Colored rectangles depict the mean and full-width-at-half-maximum of each dendritic stratum within the IPL (*vertical scale*) and the equivalent diameter (according to its diameter) of the stratum in the plane of the IPL (*horizontal scale*). Stata are colored by arbor density. Somas are drawn as circles relative to their diameter on a separate horizontal scale, as indicated. (C) Mean overlap between the stratification profile of each measured cell and each template from the Eyewire museum as cosine similarity. (D) Gallery of *en face* skeleton example images of each RGC type colored by IPL depth. Full galleries of all skeleton images and those in the Eyewire museum can be found in the Supplemental Data.

Stratification profiles for each functionally-defined RGC type are shown in **Figure 5A** along with those of our suggested match in the Eyewire museum (Bae et al., 2018). Stratification similarity between each of our types and each type in the Eyewire museum is shown as cosine overlap in **Figure 5C.** While stratification profile was an important factor in matching our types to those in the Eyewire museum, it was not the only factor. Along with stratification location and thickness, **Figure 5B** also depicts the dendritic field diameter and density of each stratum as well as the soma size for each RGC type, as measured *en face*. Example traced images of each type are shown in **Figure 5D**, and **Supplementary Data** contains all of the traced cells in *en face* and side views along with those of our suggested Eyewire match.

We combined all of our morphological measurements and used UMAP to project the data for all 136 cells into 2D. While this dataset was not large enough for clustering into ~40 types to be feasible, we measured distances in this space to capture the morphological similarity among cells that we independently grouped together by their light responses (**Figure S5**). For 30 of the 32 types represented in this dataset (those with 2 or more members), the mean pairwise morphological distance for cells of the same functional type was less than the mean pairwise morphological distance in the entire dataset. The compactness of this morphological representation for RGCs of the same functional type varied by RGC type; 10 types were more than 10-fold more compact in this space than the mean.

### Alignment of functional and transcriptomic classification

Recent large-scale investigations of single-cell transcriptomes in the retina have identified ~45 molecularly distinct types of postnatal mouse RGCs, comparable to the number of RGC types identified through physiological and morphological analyses (Rheaume et al., 2018; Tran et al., 2019). While some clusters could be matched 1:1 with previously known types based on well-established molecular markers (Sanes and Masland, 2015), approximately 40% of clusters remained unmatched. Moreover, these methods used dissociated tissue, precluding direct harmonization of gene expression with function.

To relate functional to molecular criteria, we used a variant of the Patch-seq technique (Cadwell et al., 2016) in which RGCs were first classified based on their cell-attached light responses and then the cytoplasm was collected for RNA-seq by aspirating the soma with a clean pipette (see **Methods**). We obtained 103 high-quality single RGC transcriptomes (>2000 genes/cell). We used gradient boosted decision trees (Chen and Guestrin, 2016) to match each of our transcriptomes to a cluster in the published adult RGC dataset (Tran et al., 2019)(see **Methods**). Many of our functionally-identified cells matched the transcriptomic clusters with high concordance (**Figure 6A**) providing putative matches to previously unknown clusters. For example, the three types of ON DS sus cells all aligned to C10 (a previously uncharacterized cluster), OFF tr SmRF aligned with C21, corresponding to T-RGC S2 (Liu et al., 2018) and ON delayed (Mani and Schwartz, 2017), previously observed in *CCK-ires-Cre* mice (Jacoby and Schwartz, 2018; Tien et al., 2015) aligned with a cluster (C14), which was distinguished by the expression of the neuropeptide *Cck*.

**Figure 6.**
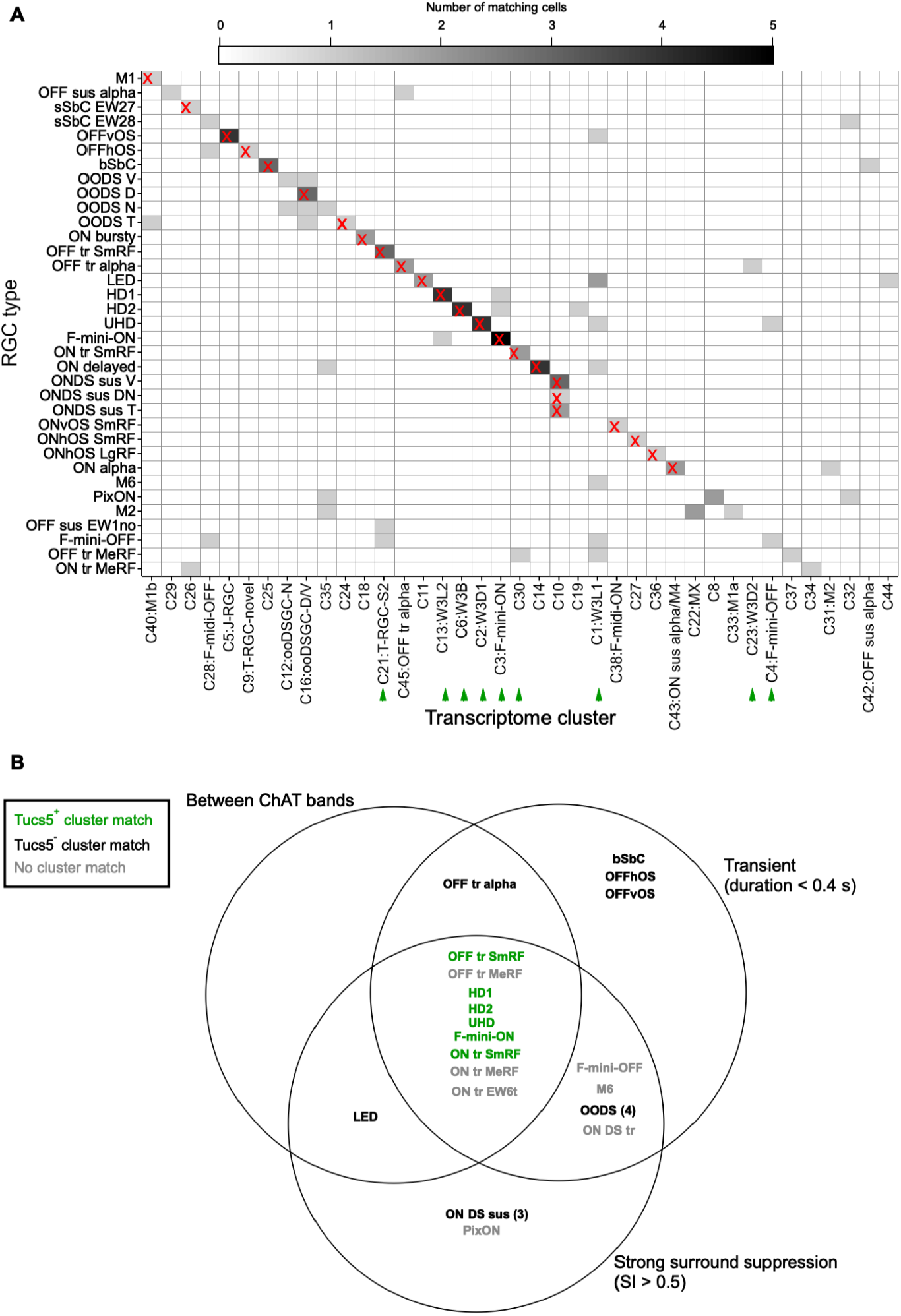
Matches between functional types and transcriptomic clusters. (A) Heatmap showing correspondence between functional types (rows) and transcriptomic clusters reported in Tran et al., 2019 (columns). Matches used in subsequent analyses are indicated by an ‘X’. Color scale indicates the number of patch-seq cells matched to each cluster. See **Methods** for matching procedure. Green arrowheads indicate T5 RGCs as described in (Tran et al., 2019). (B) Venn diagram of RGC types including one morphological characteristic (stratification between the ChAT bands) and two functional characteristics (transience and surround suppression). Green text denotes cell types matched to transcriptomic clusters identified as T5 RGCs, characterized by the specific expression of gene *Tusc5/Trarg1*, in (Tran et al., 2019).

### T5-RGCs share a functional and morphological profile

Alignment of our physiologically characterized types to transcriptomically defined RGC groups (Tran et al., 2019) enables a deeper analysis of the relationships between gene expression of RGCs and their function and morphology. One example is provided by the gene *Tusc5* (also known as *Trarg1*), which we identified as a key marker of a group of 9 mostly unidentified transcriptomic clusters termed T5 RGCs (Tran et al., 2019). Most of these RGCs are labeled by the transgene TYW3, which exhibits insertion-site dependent expression (Laboulaye et al., 2018).

Transcriptomic clusters corresponding to the T5 RGCs are labeled by green arrowheads in **Figure 6A**. Six of these clusters are matched to RGC types in our dataset, so we examined whether these types share functional or morphological characteristics. All 6 T5 RGC types lie at the intersection of two functional characteristics, transience and strong surround suppression, and one morphological characteristic, stratification between the ChAT bands (**Figure 6B**). Other subclasses of RGCs can be queried in this way, with increasing power as additional data is added to rgctypes.org.

### The question of completeness

One way to estimate the completeness of our classification is to record nearly all the RGCs in a small region of the retina and count how many can be assigned to one of our types. We performed such an experiment and then stained the tissue with the pan-RGC marker gamma-synuclein (Surgucheva et al., 2008) to confirm RGC identity post-hoc (**Figure S6**). We recorded 55 spiking cells and 25 cells for which we could not elicit spikes with our test stimuli. Of the 55 spiking cells, 48 were successfully identified in the fixed tissue. In the live tissue we had labeled 42 of these cells as RGCs matching one of our types and 6 as spiking amacrine cells. All 48 of 22 were identified in the fixed tissue: 10 were gamma-synuclein negative, presumably non-spiking amacrine cells, and 12 were gamma-synuclein positive, presumably RGCs that we failed to identify. Thus, we identified 78% (42 / 54) of the putative RGCs in this sample. While somewhat less than our estimate of 89% coverage of the types in the Eyewire museum, it is a conservative estimate because some of the non-responding RGCs these identifications were verified by the gamma-synuclein staining (42/42 gamma-synuclein positive RGCs and 6/6 gamma-synuclein negative spiking amacrine cells). Of the 25 cells for which we could not elicit spikes, were likely damaged during removal of the inner limiting membrane or by the recording procedure and did not spike (e.g. because the axon initial segment was destroyed) but survived enough structurally for gamma-synuclein staining.

### Relatedness of functional, morphological, and transcriptomic space

The main goal of our study was to directly relate physiological, structural and molecular definitions of cell type. Our suggested alignments between these three modalities are shown in **Figure 7A**. Functional types are colored by their F-scores from **Figure 4**, and the data used to infer the alignment is shown in **Figures 5, 6**, and **Supplemental Data**. With this alignment data in hand, we were able to address an additional question: to what extent do relationships among types established in one modality (e.g. function) predict those in another modality (e.g. morphology). Importantly, this is not a test of the quality of our alignment between modalities. Functionally similar RGC types might differ substantially in morphology and/or gene expression, and the degree to which local neighborhoods are similar across modalities might vary for each RGC type.

**Figure 7.**
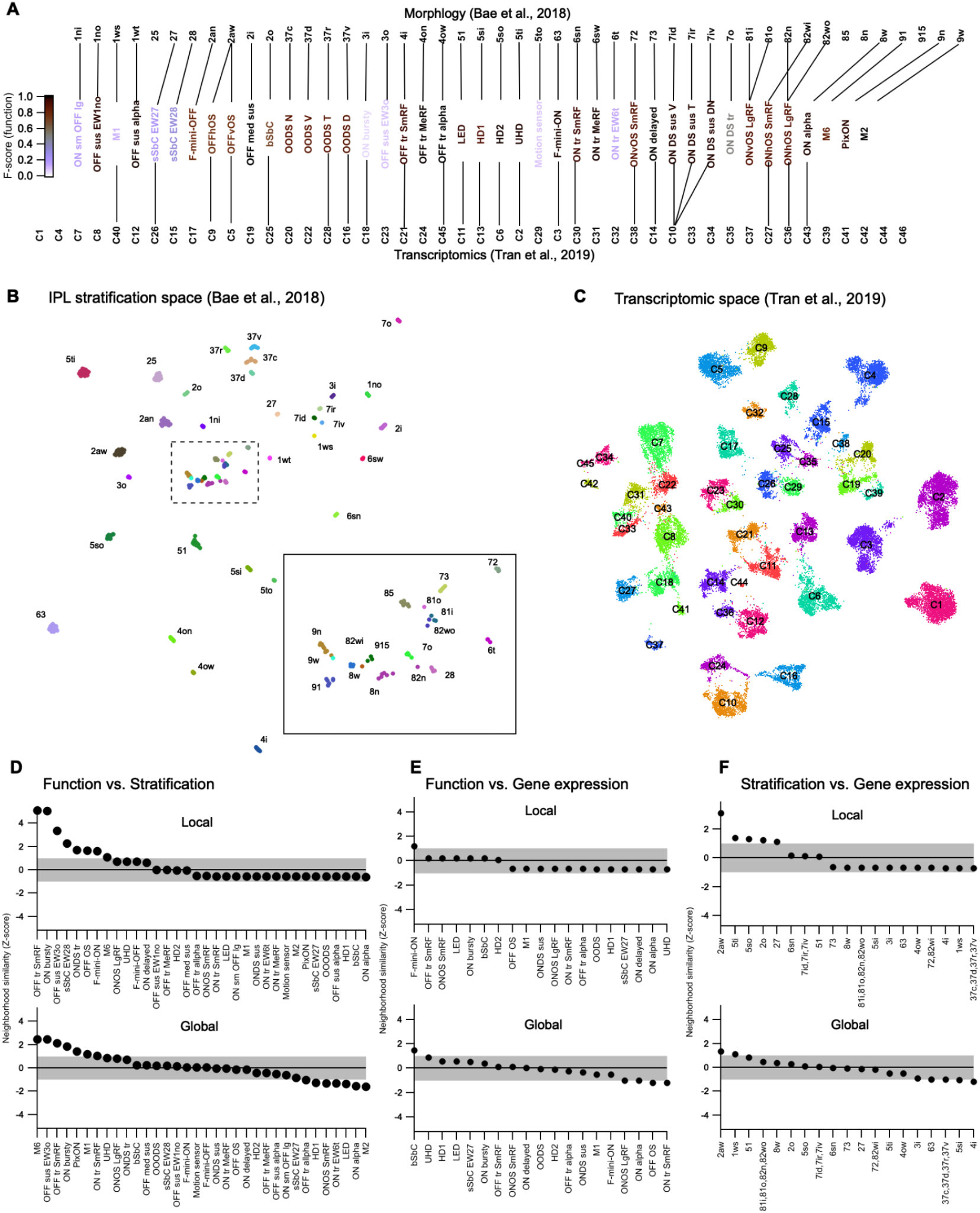
Correspondence between RGC relatedness in functional, morphological, and transcriptomic space. (A) UMAP embedding of RGC morphology constructed from the stratification profiles in the Eyewire museum (Bae et al., 2018). Inset shows boxed region at higher magnification. (B) UMAP embedding of RGC gene expression from Tran et al. (2019). Cluster labels removed for clarity. (C) Alignments between the three classification schemes that we used for subsequent analysis. Lines connect putative corresponding RGC types in each classification schema. (D) List of RGC types ranked by the z-normalized fractional overlap between functional and stratification embeddings. Shaded region indicates 1 s.d. around the expectation from the null distribution. *Top*, local neighborhood (2-4) neighbors; *bottom,* global neighborhood (5-12 neighbors). (**E, F**) Same as (**D**) but showing alignment between functional and morphological space (**E**) or morphological and gene expression space (**F**). Local neighborhood for (**E, F**) is 2-3 neighbors and global neighborhood is 4-8 neighbors.

To investigate the questions of cross-modality neighborhood similarity, we constructed a UMAP embedding of the stratification profiles of each cell in the Eyewire museum (Bae et al., 2018)(**Figure 7B**) and another UMAP embedding of gene expression from the mouse RGC transcriptomic atlas (**Figure 7C**; replotted from (Tran et al., 2019)). To measure neighborhood similarity across the 3 UMAP spaces (the functional space from **Figure 3A** and the stratification and gene expression spaces in **Figure 7B,C**), we tested whether the nearest neighbors in a reference modality were also grouped nearby in another modality.

For each RGC type, we computed the fractional overlap among the identities of its nearest neighbors in the reference embedding to that in the other two embeddings. We repeated this analysis for neighborhood sizes from 2 to 12 nearest neighbors and grouped the results into the “local” and “global” neighborhoods. To establish statistical significance on this fractional overlap measure, we used the bootstrap approach. We randomly shuffled type identities in each of the maps and recomputed the fractional overlap. Repeating this process 1000 times yielded an empirical null distribution. Fractional overlap values obtained from the real data are reported as z-scores relative to this null distribution with positive values indicating greater overlap in the real data than in the null distribution (**Figure 7D–F**).

Overall, similarity between modalities was modest; cross-modality overlap for many RGC types was within 1 s.d. of the null distribution (shaded regions in **Figure 7D–F)**. Several RGC types did show strong local neighborhood similarity between functional and morphological (IPL stratification) embeddings (**Figure 7D**), and one type (OFF OS; 2aw; C5 and C9) showed a strong correspondence between its local neighborhoods in stratification and gene expression space. Global neighborhood alignments had similar overall trends but a somewhat different set of RGCs tended to be more cross-modally aligned globally than locally.

### Integrated web-based RGC compendium

Finally, we created a resource so that labs around the world can come to a consensus on the classification of mouse RGCs. To that end, we have developed a website, rgctypes.org (**Figure 8**), with a direct pipeline to our database of functional and morphological measurements. Following a curation step and type assignment, every RGC recorded in the Schwartz lab will automatically update to rgctypes.org. Other researchers are invited to submit data for integration as well. Cells can also be reassigned to different types if evidence supports a different assignment. Full datasets are available for download immediately, regardless of publication status. We have also provided a downloadable version of our automated classifier and instructions on how to prepare a data file to obtain a type prediction and confidence score.

**Figure 8.**
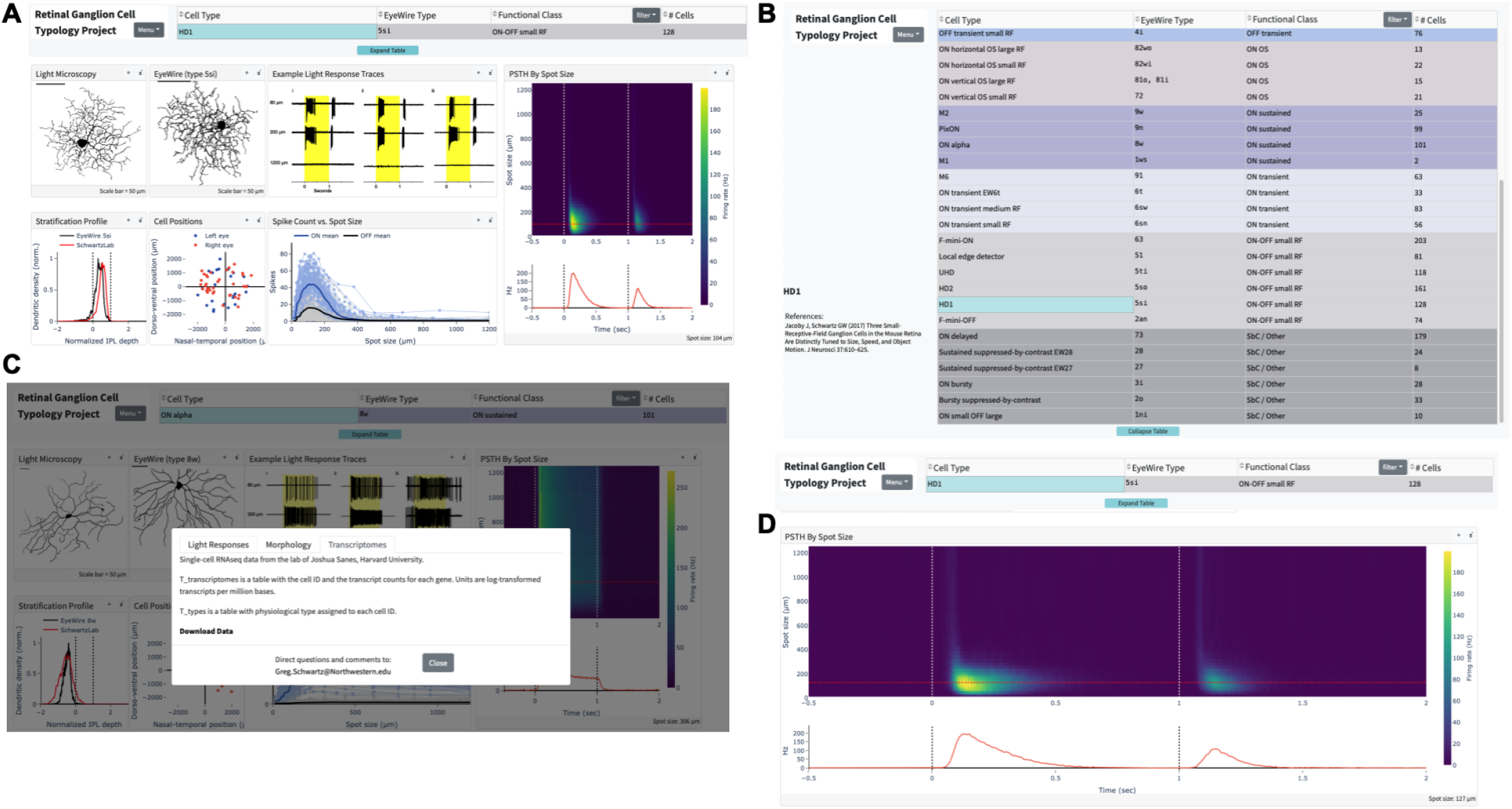
Screenshots from rgctypes.org. (A) Landing page for the HD1 RGC. (B) Table of RGC types (C) Data download area. (D) Expanded, interactive graph of HD1 RGC light responses.

## Discussion

We present a resource of physiological, morphological, and transcriptomic data aimed at establishing a comprehensive typology of mouse RGCs. A summary of our classification and its alignment with previous RGC classifications is provided in **Table S3**. As multi-modal neuronal classification efforts continue to be a major focus across many labs (BRAIN Initiative Cell Census Network (BICCN), 2021), we first consider what lessons we have learned from this approach in our dataset that might apply to other regions of the CNS before discussing what our findings have revealed about the retina.

RGCs have a distinct advantage for studies of typology since they form mosaics to tile visual space. Several lines of evidence have now converged on a number of types near 45 in the mouse (Baden et al., 2016; Bae et al., 2018; Tran et al., 2019), and we find 42 types with an estimated coverage of 89%. Each type has functional characteristics that we used to distinguish it from others (**Figure 2**), and with few exceptions, these differences were captured by supervised dimensionality reduction of the spike responses to a simple stimulus (**Figure 3**). Success in clustering responses, however, does not automatically translate into success for an automated classification algorithm (**Figure 4**). In clustering, there is typically no external ground truth data to assess the validity of the clusters as cell types. We also lacked an absolute ground truth, but we used external validation data not available to our classifier to label ⅓ of our cells (**Table S2**) and found performance to be similar (or slightly better) than on our unvalidated type labels. When external validation data is available, future studies of cell typology should report the performance of a cross-validated classifier in addition to measures of cluster separability.

As more studies employ multiple modalities, like function, morphology, and gene expression, to classify neurons, comparisons of the same cells between modalities will become more frequent. Gene expression impacts both morphology and function, and stratification within the IPL is an important factor in determining the synaptic inputs of RGCs. Thus, one might have expected an even stronger correspondence between the positions of RGC types across modalities (**Figure 7D-F**). It is worth noting that such an analysis inevitably simplifies across the large possible space of each modality by dimensionality reduction both at the level of feature selection (a single stimulus type, IPL stratification alone for morphology, gene selection for transcriptomics) and at the level of the UMAP algorithm (down to 2 dimensions for each modality). For example, specific single genes might be very important for IPL lamination (Krishnaswamy et al., 2015; Liu et al., 2018) yet might fail to group types in transcriptomic space. The high dimensionality required to fully specify a cell type in any single modality might mean that new distance metrics will be needed to study cross-modal relationships between cell types if such relationships turn out to be important principles of brain architecture or development.

### Method for functional classification

We recorded from RGCs one at a time, which allowed us to center stimuli on the receptive field of each cell. This undeniably limits throughput. On the other hand when activities of many RGCs are recorded simultaneously – for example by calcium imaging or with multielectrode arrays – it is not feasible to center stimuli on individual RGCs, so these studies have used a combination of full-field modulation, large moving objects or gratings, and spatiotemporal white noise. These stimulus choices come with a significant cost. Many RGC types, including some of the most numerous types, respond poorly or not at all to full-field stimuli or spatiotemporal white noise (Jacoby and Schwartz, 2017; Zhang et al., 2012). Other types respond both to small (RF-centered) and large stimuli, but basic response properties depend on spot size. For example, the ON small OFF large RGC would be classified as an OFF cell for full-field stimulation but responds as an ON cell for small spots in its RF center. Surround suppression differentially affects both the total spike count and response kinetics in most RGC types (**Figure 2**)(Wienbar and Schwartz, 2018), providing information that we found necessary to separate otherwise functionally similar types.

### Comparisons to previously defined RGC types

Our 42 RGC types appear to include all 28 types previously identified functionally (referenced in **Table S3** and at rgctypes.org) as well as 14 “novel” types that have not, to our knowledge, been defined previously. Remarkably, most of these types can be distinguished based on their response patterns to spots of varying size. The total is close to previous estimates (Baden et al., 2016; Bae et al., 2018; Rheaume et al., 2018; Tran et al., 2019), supporting the view that mouse RGC classification is approaching completion. Many of the “novel” types had certainly been encountered in previous studies, but we list them as such here based on our belief that they had not been identified separately as distinct functional types (e.g. multiple types had been grouped into “ON transient” and “OFF transient” categories). The novel types include several sets of functionally similar RGCs (ON tr MeRF / ON tr SmRF / ON tr EW6t, OFF tr MeRF / OFF tr SmRF, OFF med sus / OFF sus EW1no / OFF sus EW3o), all of which match 1:1 to morphological types and many to transcriptomic types.

Why did the retina evolve entire populations of RGCs that vary only subtly in function? Of many possible answers, we believe the most likely is that functionally similar types would reveal profound differences under stimulus conditions beyond those in our simple battery. A striking example is Eyewire type 25. This type is abundant (5.8% of the population), and forms a convincing and statistically validated dendritic mosaic (Bae et al., 2018), yet we were unable to find its match in thousands of recordings. A natural hypothesis is that this RGC type does not respond to our standard test stimuli, so it was consistently passed over. Supporting this idea, the calcium responses for type ‘25’ in the Eyewire museum are weak (~1% ΔF/F with low signal-to-noise ratio as opposed to some RGC types which reached 20% ΔF/F). Similarly, we failed to find a clear “trigger feature” for several RGC types (e.g. ON bursty, Motion sensor, sSbC EW27). Responses of these cells to flashed spots were inconsistent. For simplicity and reproducibility, our study omitted the vast space of light stimuli that may have differentiated these cell types, including high luminance, variations in color, and complex forms of motion.

Direction selective (DS) and orientation selective (OS) RGCs represent a substantial fraction of the RGCs in the mouse retina (14/42; 33% of types). We identified ON-OFF DS RGCs preferring all four cardinal directions (dorsal, ventral, nasal, temporal), ON DS sustained types preferring three different directions, and one ON DS transient type encountered infrequently and with a wide distribution of preferred directions (**Figure S7**). While there is broad agreement that there are four ON-OFF DS RGC types in the mouse, there is not as strong a consensus about ON DS RGCs. Some studies have reported three types (Estevez et al., 2013) while another reported four (Sabbah et al., 2017). It remains unclear whether this discrepancy is due to one of the ON DS RGC types being transient and the other three being sustained. One study reported a functionally and morphologically distinct ON DS RGC that projects to superior colliculus (SC) and not to the medial terminal nucleus (MTN) or nucleus of the optic tract (NOT) of the accessory optic system (Gauvain and Murphy, 2015). This SC-projecting ON DS RGC had transient responses and more balanced ON and OFF dendritic strata than the MTN-projecting types, consistent with type ‘7o’ in the Eyewire museum. While the previous study on these cells did not report the distribution of their preferred directions (Gauvain and Murphy, 2015), calcium responses for Eyewire type ‘7o’ consistently preferred a nearly nasal direction on the retina (Bae et al., 2018). Our sample of ON DS sustained RGCs had a distribution of preferred directions with three clusters, separated by ~120 degrees, but the sparsely sampled ON DS transient RGCs had inconsistent direction preference (**Figure S7**), and we have so far been unable to reconstruct its morphology. Thus, we have provisionally assigned the ON DS trans. RGC to Eyewire type ‘7o’, but it is one of the matches in which we have the least confidence. A more focused study on ON DS RGCs will be needed to resolve this final issue in the classification of DS RGC types.

OS RGCs, described long ago in other species (Levick, 1967; Maturana and Frenk, 1963), were only recently identified in the mouse (Nath and Schwartz, 2016, 2017). OFF OS RGCs were separated into horizontal- and vertical-preferring types based on their physiology, and the vertical-preferring type tended to have ventrally directed dendrites, while horizontally-preferring cells had a less consistent asymmetry (Nath and Schwartz, 2017). The Eyewire data did not have a corresponding type consisting only of cells with strong ventrally directed dendrites, although they note that type ‘2aw,’ with its similar range in dendritic asymmetry, has a much higher coverage factor than the other types and likely corresponds to at least two RGC types that were not separable based on morphology alone (Bae et al., 2018). Given these facts and the corresponding stratification patterns between these types, we are confident in the categorization of both OFFhOS and OFFvOS RGCs as Eyewire type ‘2aw’.

ON OS RGCs were also classified into horizontal- and vertical-preferring types when they were reported in mouse (Nath and Schwartz, 2016), but here we further subdivide each group into separate “Small RF” and “Large RF” types based on the spot size to which they respond optimally and their degree of surround suppression. All four ON OS RGC types are among the largest in the retina in terms of dendritic span, so their morphology is captured incompletely in the Eyewire dataset. Nonetheless, we have been able to assign each of these functional OS RGC types to its most likely matching morphological type.

We identified three RGC types as suppressed-by-contrast (SbC), and a fourth, the ON delayed RGC, has been classified as an SbC RGC under some conditions (Jacoby and Schwartz, 2018; Tien et al., 2015). The RGC type we originally identified as the sustained SbC (Jacoby et al., 2015) has now been split into two types (EW27 and EW28) based on both physiological and morphological criteria. The bursty SbC (bSbC) RGC is distinguished from the sustained SbC types by its much higher baseline firing rate, more transient suppression, and monostratified morphology (Wienbar and Schwartz, 2021). Overall, our data underscores the fact that, like the other three polarities (ON, OFF, and ON-OFF), SbC is a response class composed of multiple RGC types (Jacoby and Schwartz, 2018).

### Relationships between morphology, function, and gene expression

Having matched functional, morphological, and transcriptomic information for most RGC types, we were able to assess the relationships among these properties. Disappointingly, proximity of types as assessed by any single criterion failed to strongly predict proximity by either of the other two criteria. For transcriptomic relationships, one possibility is that genes expressed during development, when morphology and connectivity are being established, will need to be taken into account. On the other hand, the comparison between morphology and funuctin was valuable in highlighting three unexpected trends. First, there are many exceptions to the rule that RGCs with dendrites in the outer half of the IPL have OFF responses. The M1 ipRGC was a well-known exception, because it receives ectopic synapses from ON bipolar cells in the outer IPL (Dumitrescu et al., 2009), but it is far from the only exception to this rule. All four ON OS RGC types, the ON delayed, the M6, and both sSbC types have OFF dendrites but lack OFF spike responses. Additionally, the OFF OS RGCs and the F-mini-ON RGC receive OFF input via gap junctions but lack OFF bipolar cell input under any stimulus condition we have tested (Cooler and Schwartz, 2020; Nath and Schwartz, 2017). An important caveat is that stimuli beyond our test set could reveal OFF responses, perhaps in bright conditions (Pearson and Kerschensteiner, 2015; Tikidji-Hamburyan et al., 2015).

Second, the dendritic area of an RGC has often been associated with the size of its RF center. While this association has a strong basis in the anatomy of the vertical excitatory pathways of the retina, there are a number of exceptions in our data set. For example, “Small RF” and “Large RF” ON OS RGC types do not differ appreciably in dendritic area, and M6 RGCs have smaller RFs than ON delayed RGCs despite substantially larger dendritic area. Differential influences of inhibition and disinhibition likely explain some of these effects (Mani and Schwartz, 2017; Wienbar and Schwartz, 2018).

Finally, RGCs with dendrites near the inner and outer margins of the IPL are typically assumed to have more sustained light responses while those stratifying near the middle of the IPL are assumed to be more transient (Awatramani and Slaughter, 2000; Roska and Werblin, 2001). This association has gained support from large-scale measurements of the kinetics of glutamate release from bipolar cells throughout the IPL (Franke et al., 2017; Marvin et al., 2013). While our data generally fit this trend, there were two notable exceptions. The M6 RGC is transient despite stratification at both margins of the IPL, and the LED RGC is sustained despite stratification near the middle of the IPL (Jacoby and Schwartz, 2017).

The literature linking gene expression in particular RGC types to their morphology and function has been more fragmentary because the lack of known matches has prevented a wide view. We found that expression of the gene *Tusc5* is strongly associated with a particular physiological profile (transient light responses and strong surround suppression) and a morphological profile (stratification between the ChAT bands)(**Figure 6B**). As more information about each of the RGC types becomes available, including their projection patterns in the brain, we expect more insights into the molecular determinants of RGC wiring patterns both within the retina and to the brain. Future studies may also link biophysical properties of RGCs to the expression of ion channels.

### Limitations of the dataset and future directions

Several limitations of our dataset suggest directions for future work. First, our stimuli were limited to a single wavelength distribution, a small range of scotopic to mesopic luminance, and a simple set of artificial patterns (spots, gratings, and moving bars). These stimulus choices meant that we could not explore how RGC responses differed over the parameters of luminance or wavelength. More generally, RGCs evolved not for selectivity to the artificial parameterized stimuli we presented but to detect behaviorally relevant features of natural scenes. Second, while centering the stimulus for each RGC was important for measuring the spatial features of its response, this step complicates the recovery of locally-complete RGC mosaics. Therefore, a future step in RGC typology alignment will be needed to match our types with those in large-scale recordings using either calcium imaging or multi-electrode arrays. We hope to collaborate with other labs performing large-scale RGC recordings with some version of a sparse noise stimulus to validate the robustness of our functional classification across labs and preparations. Finally, our morphological alignment to the Eyewire dataset was not validated by a classification algorithm. The limited number of cells in both datasets and their methodological differences made such a morphology classifier impractical, but with additional data an RGC morphology classifier is a goal (Laturnus and Berens, 2021). Since our functional classification algorithm produces a posterior probability for each class, functional and morphological information could be incorporated seamlessly into a single prediction. Similarly, our improving understanding of the gene expression profiles of each RGC type could enable more accurate composite predictions from the expression of a few key genes plus functional and/or morphological data.

### Web-based resource

Standardization in the definitions of RGC types among different research groups is essential to support studies on retinal computation, circuit connectivity, and disease pathology. Additionally, there is rapidly expanding interest in the projection patterns of different RGC types throughout the brain (Dhande et al., 2015; Johnson et al., 2021; Martersteck et al., 2017), which similarly relies on standardized type definitions. For these reasons, we created an open online resource at rgctypes.org where users can search and download full datasets, use our classification algorithm, and contribute their own data to this effort. By unifying the separate functional, morphological, and molecular RGC classification schemas, this resource will allow researchers to connect data across experimental modalities. For example, a set of RGCs labeled by their projection to a certain brain region could be classified by gene expression, and our alignment between transcriptomics and function would provide insights into the functional input to that brain region without additional measurements of light responses. Or the complement of RGC types in a new transgenic mouse line could be measured by confocal microscopy, and our alignment between morphology and function could help generate hypotheses about the functional deficits that might exist if this RGC population were ablated. We expect rgctypes.org to play a central role in the fields of retinal neurobiology and vision science moving forward and, more broadly, to serve as a template for data sharing and collaboration that is applicable to neuronal classification projects throughout the CNS.

## Methods

### Animals

Wild-type mice (C57/Bl6 - JAX 000664) of either sex were dark-adapted overnight and sacrificed according to standards provided by Northwestern University Center for Comparative Medicine. 4 transgenic lines were used to target subsets of RGCs. All other mice were WT

PV-Cre (JAX #008069) x Ai14 (JAX #007908): 4 animals

Opn4-GFP (Generous gift from lab of Tiffany Schmidt, Northwestern University): 2 animals

TYW3-GFP (Lab of author J. Sanes): 9 animals

JAMB-eYFP (Lab of author J. Sanes): 3 animals

Retinal tissue was isolated under infrared illumination (900 nm) with the aid of night-vision goggles and IR dissection scope attachments (BE Meyers). Retinal orientation was identified using scleral landmarks (Wei et al., 2010), and preserved using relieving cuts in cardinal directions, with the largest cut at the ventral retina. Retinas were mounted on 12mm poly-D-lysine coated glass affixed to a recording dish with grease, with the ganglion cell layer up. Oxygenation was maintained by superfusing the dish with carbogenated Ames medium (US Biological, A1372-25) warmed to 32°C. Our dataset included 1859 recorded RGCs from 551 eyes of 544 animals.

### Visual stimuli

RGC types were identified via cell-attached capacitive spike train responses to light stimuli as previously described (Jacoby and Schwartz, 2017; Jacoby et al., 2015; Mani and Schwartz, 2017; Nath and Schwartz, 2016, 2017). Briefly, stimuli were presented using a custom designed light-projector (DLP LightCrafter; Texas Instruments) at a frame rate of 60 Hz. Spatial stimuli patterns were generated on a 912×1140-pixel digital projector using blue (450nm) LEDs focused on the photoreceptor layer. Neutral density filters (Thorlabs) were used to attenuate the light intensity of stimuli to 200 rhodopsin isomerizations per rod per second (R*/rod/s) from darkness.

The receptive field (RF) centers of individual RGCs were determined by monitoring their relative light responses to horizontal and vertical bars (200 x 40 μm, or 100 x 40 μm in the case of cells with high surround suppression) flashed at 30 μm intervals at 11 locations along each axis. Subsequent stimuli were presented at the RF center. For generic light steps, a spot of 200 μm diameter was presented for 1 s, with cell-attached responses recorded for at least 0.5 s pre-stimulus and 1s post-stimulus. For spots of multiple sizes, spots with diameters from 30-1200 μm (on a logarithmic scale) were presented in pseudorandom order, with similarly timed epochs. Direction preference of direction-selective (DS) RGCs was determined by moving bar stimuli, consisting of a rectangular bar (600 x 200 μm) passing through the receptive field center at 1000 μm/s (ON-OFF DS RGCs) or 500 μm/s (ON DS RGCs). Flashed bar stimuli for testing orientation selectivity were 800 x 50 μm and presented at 12 different orientations (Nath and Schwartz, 2016). Drifting gratings and contrast series were presented from a background luminance of 1000 R*/rod/s following protocols from previous studies (Jacoby et al., 2015; Nath and Schwartz, 2017).

### Functional response metrics

We measured 6 standard response metrics from the flashed spots data. Distributions for each metric for each RGC type are shown in **Figure S4**.

*Baseline firing rate*. Mean firing rate in darkness before spot presentation across all trials.

*Peak firing rate*. Highest firing rate (baseline subtracted) achieved in a 10 ms bin at light onset or offset across all spot sizes.

*Peak response latency*. Time from light onset or offset until the peak firing rate.

*Response duration.* Time from peak firing rate until firing rate drops below baseline +10 percent.

*Suppression index*. First we determined the dominant polarity for the cell by whether the maximum ON or OFF response (in total spike count from baseline) was larger across spot sizes. For the dominant polarity, the suppression index was the ratio of the difference between this maximum response and the response to the largest (1200 μm) spot divided by the sum of these two quantities.

*ON:OFF index.* Maximum ON response across spot sizes (spike count from baseline) minus the maximum OFF response divided by the sum of these two quantities.

### Automated classification

A classifier was trained to recognize RGC types based on cell-attached recordings of responses to spots of multiple sizes. RGC type labels were assigned manually by two of the authors (JG and GWS), and cells were further labeled as externally validated or unvalidated by GWS based on the presence of identifying data not available to the classifier, including morphological, transcriptomic, whole-cell, and physiological (response to moving bars or drifting gratings) data. Types with 5 or fewer examples were excluded from training, and OS and DS cells were condensed across orientation/direction based on the similarity of their light responses to spots of multiple sizes.

Cells were randomly assigned to a testing set (~25% of cells) and a training set, which was further subdivided into a training set for a base classifier (~50% of cells) and a calibration set (~25% of cells). The scheme favored assignment of validated cells to the calibration set and unvalidated cells to the training set for the base classifier. The classifier implements a semi-supervised learning model: a base classifier learns to recognize features of the probability distribution of RGC light responses that are useful for predicting the manual labels, subject to labeling error; this knowledge is “transferred” by the calibrator to reweight the base model’s predictions in order to better predict the labels which are influenced by external validation. Thus we minimize error propagation while maintaining a large enough training set to form robust predictions about RGC type.

The multi-class classification problem was broken down into a series of binary ones using the error-correcting output code (ECOC) scheme, such that a series of classifiers each learns to discern different combinations of RGC types. Each binary learner in the ECOC scheme was trained using Ada-boosted decision trees (Hastie et al., 2009) with initial weights set to enforce a uniform prior probability of each RGC type.

Individual trees were trained by performing elastic net logistic regression on a random subset of firing rates from peristimulus time histogram (PSTH) vs. spot size for feature reduction and choosing the threshold that minimized class uncertainty (Friedman et al., 2010; Schneider et al., 2015). Since not all PSTHs were recorded over the same time and spot size ranges, we imputed missing data using a nearest neighbor approach. Poorly sampled points were penalized in both random selection and regression: for time points the penalty was inversely proportional to their frequency of occurrence across cells (since all PSTHs were binned with the same Δt); for spot sizes we aimed to account for the nonlinearity of responses in the penalty with the following formula:

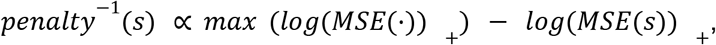

where MSE is the mean across cells of the squared error between the chosen spot size, *s*, and the nearest recorded spot size, and (·)_+_ denotes positive rectification.

To implement the calibrator, the calibration fold was used to train an isotonic regression model that transformed each binary learner score into a probability, again enforcing a uniform prior using sample weighting (Zadrozny and Elkan, 2002). The probabilities from each binary learner were then coupled to obtain a probability for each class (Zadrozny, 2002).

We used three-fold cross validation to train a Bayesian optimization model for hyperparameter tuning. Table 3 lists the hyperparameters we optimized and their final values. The classifier is available for use at rgctypes.org, and the source code is available at https://github.com/zfj1/rgc-classifier.

**Table 1.**
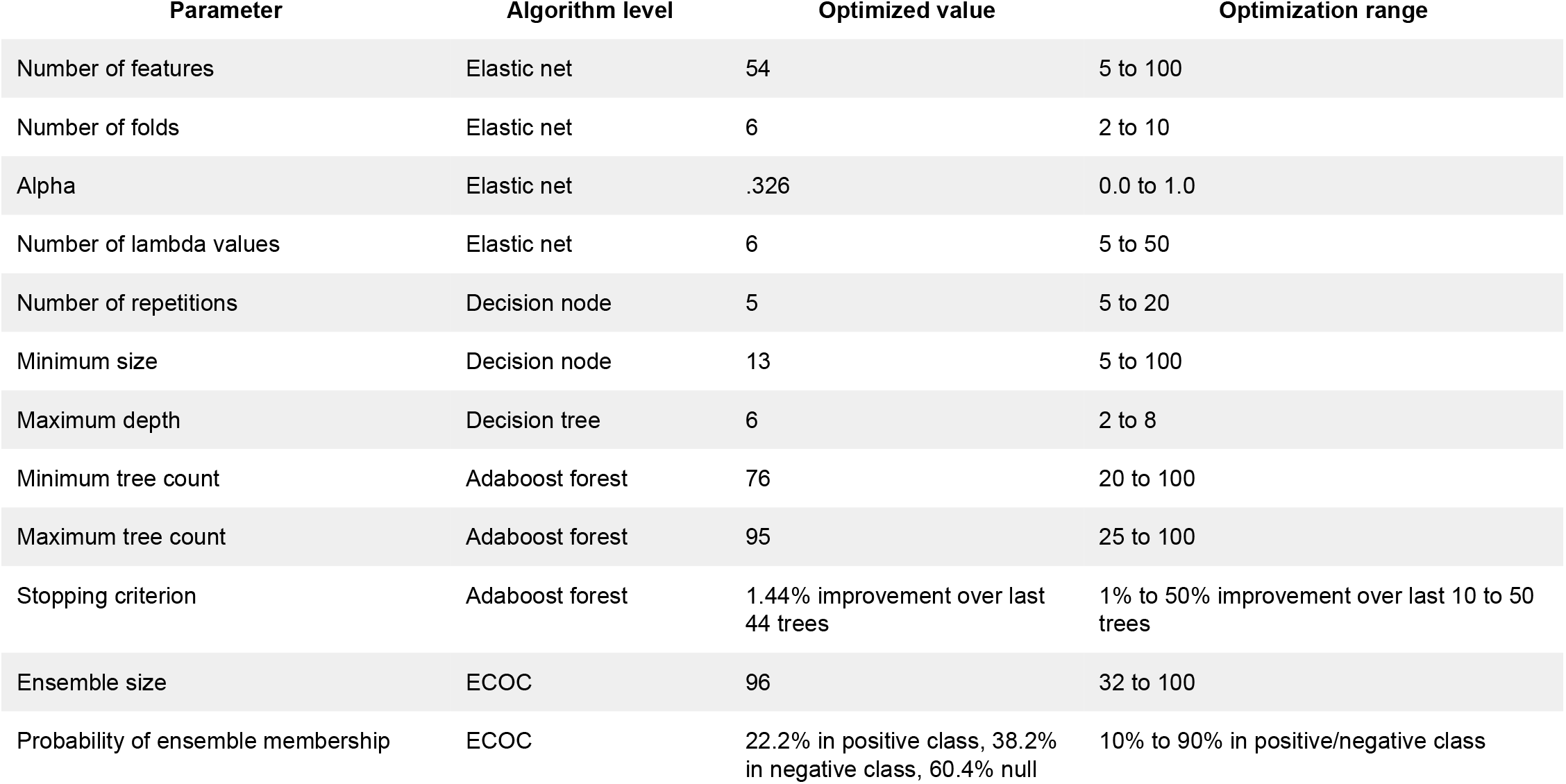
Hyperparameters for the automated RGC classifier.

| Parameter | Algorithm level | Optimized value | Optimization range |
| --- | --- | --- | --- |
| Number of features | Elastic net | 54 | 5 to 100 |
| Number of folds | Elastic net | 6 | 2 to 10 |
| Alpha | Elastic net | .326 | 0.0 to 1.0 |
| Number of lambda values | Elastic net | 6 | 5 to 50 |
| Number of repetitions | Decision node | 5 | 5 to 20 |
| Minimum size | Decision node | 13 | 5 to 100 |
| Maximum depth | Decision tree | 6 | 2 to 8 |
| Minimum tree count | Adaboost forest | 76 | 20 to 100 |
| Maximum tree count | Adaboost forest | 95 | 25 to 100 |
| Stopping criterion | Adaboost forest | 1.44% improvement over last 44 trees | 1% to 50% improvement over last 10 to 50 trees |
| Ensemble size | ECOC | 96 | 32 to 100 |
| Probability of ensemble membership | ECOC | 22.2% in positive class, 38.2% in negative class, 60.4% null | 10% to 90% in positive/negative class |

### Imaging

A subset of recorded RGCs were injected with Neurobiotin (Vector Laboratories, SP-1150, ∼3% w/v and ∼280 mOsm in potassium aspartate internal solution) using patch pipettes. Retinas were then fixed in 4% paraformaldehyde for 15 minutes at 25°C, washed three times with PBS, and incubated for 1 hour in blocking solution (1X PBS with 3% normal donkey serum, 0.05% sodium azide, 0.5% Triton X-100) including streptavidin conjugated to a fluorophore (Alexa Fluor-488 or Alexa Fluor-568). Next, retinas were incubated again in blocking solution with primary antibody against choline acetyltransferase (ChAT; Millipore, AB144P, goat anti-ChAT, 1:1000) for 5 nights at 4°C. Retinas were then rinsed in PBS three times at no less than 1 hour per wash before incubation overnight at 4°C with streptavidin (Jackson, 016-600-084) and secondary antibody (Donkey anti-Goat 647, Fisher, A11055). Retinas were then rinsed again in PBS three times at no less than 1 hour per wash before mounting on slides with Fluoromount.

RGCs filled with AlexaFluor were imaged immediately using two-photon microscopy (920 nm, MaiTai HP; SpectraPhysics) under a 60× water-immersion objective (Olympus LUMPLan FLN 60×/1.00 numerical aperture). A 520–540 nm band-pass filter was used to collect emission. After immunohistochemistry, confocal imaging was performed at the Center for Advanced Microscopy at Northwestern University Feinberg School of Medicine generously supported by NCI CCSG P30 CA060553 awarded to the Robert H Lurie Comprehensive Cancer Center. Dendrites were traced in Fiji using the SNT plugin (Arshadi et al., 2021).

### Morphology analysis

Dendrite skeleton images were flattened using a custom MATLAB tool based on the method in (Sümbül et al., 2014) and available at github.com/SchwartzNU/SymphonyAnalysis/tree/master/imageAnalysis/RGC_analyzer. In cases where we had ChAT staining, the ChAT bands were used as the reference surfaces. In cases where ChAT staining was not available, we used a smoothed version of each (hand-selected) stratum as reference surfaces and used soma position to register to IPL depth. In addition to the stratification profile, we computed 10 additional metrics from each arbor skeleton. All of these metrics were combined for the unsupervised clustering analyzed in **Fig. S5**.

*Area*. Area of the polygon connecting the tips of the dendrites in each stratum.

*Convexity Index*. Convex hull area around each stratum divided by the polygon area.

*Total length*. Linear length of dendritic tree.

*Arbor density*. Linear length divided by area for each stratum.

*Arbor complexity.* Number of branches divided by total length.

*Soma size.* Diameter of soma at its largest axial cross section.

*Branch length.* Distribution of lengths of all branches of the dendritic tree.

*Branch angle.* Distribution of angles at branch points in the arbor.

*Tortuosity*. Distribution of path length divided by Euclidean distance between endpoints for each branch.

*Depth range*. Distribution of range in depth in the IPL spanned by each branch.

### Single-cell transcriptomics

#### Library generation

Following physiological recording, a subset of RGCs was isolated for single-cell transcriptome sequencing. First, the area surrounding cells of interest was cleaned of nearby cells and visible debris by aspiration through a large (3-4um inner diameter) patch pipette. Cells were then aspirated using a freshly flame-pulled patch pipette (2.5 inner diameter) and placed into a 5 μl of lysis Buffer TCL (Qiagen, 1031576) + 1% 2-mercaptoethanol (Millipore-Sigma, 63689) before being flash-frozen on dry ice.

We generated RNA-Seq libraries using a modified Smart-seq2 method (Picelli et al., 2014) with the following minor changes: Before reverse transcription, RNA was purified using 2.2X SPRI-beads (Beckman Coulter, A3987) followed by 3 wash steps with 80% EtOH, elution in 4 μl of RT primer mix and denatured at 72 °C for 3 min. Six μl of the first-strand reaction mix, containing 0.1 μl SuperScript II reverse transcriptase (200 U/μl, Invitrogen), 0.25 μl RNAse inhibitor (40 U/μl, Clontech), 2 μl Superscript II First-Strand Buffer (5x, Invitrogen), 0.1 μl MgCl_2_ (100 mM, Sigma), 0.1 μl TSO (100 μM) and 3.45 μl Trehalose (1M), were added to each sample. Reverse transcription was carried out at 50°C for 90 min followed by inactivation at 85 °C for 5 min. After PCR preamplification, product was purified using a 0.8X of AMPure XP beads (Beckman Coulter), with the final elution in 12 μl of EB solution (Qiagen). For tagmentation the Nextera DNA Sample Preparation kit (FC-131-1096, Illumina) was used and final PCR was performed as follows: 72 °C 3 min, 95 °C 30 s, then 12 cycles of (95 °C 10 s, 55 °C 30 s, 72 °C 1 min), 72°C 5min. Purification was done with a 0.9X of AMPure XP beads. Libraries were diluted to a final concentration of 2 nM, pooled and sequenced on Next-Seq(Mid), 75bp paired end.

#### Alignment and quantification of scRNA-cell transcriptomic libraries

Gene expression levels were quantified using RNA-seq by Expectation Maximization (RSEM) (Li and Dewey, 2011). Under the hood, Bowtie 2 (Langmead and Salzberg, 2012) was used to map paired-end reads to a mouse transcriptome index (mm10/GRCm38 UCSC build). RSeQC (Wang et al., Bioinformatics, 2012) was used to quantify quality metrics for the alignment results. We only considered cells where the read alignment rate to the genome and transcriptome exceeded 85% and 35% respectively, and the total number of transcriptome-mapped reads was less than 350,000. RSEM yielded an expression matrix (genes x samples) of transcript per million counts (TPM), which were log-transformed after the addition of 1 to avoid zeros. Overall 103 RGCs, each of which carried a functional type label, were selected for further analysis.

#### Matching gene-expression clusters to cell types

To map each of the 103 RGC transcriptomes to a molecular cluster in Tran et al., 2019 we used the XGboost algorithm (Chen and Guestrin, 2016), as implemented in the R package xgboost. Briefly, we trained and validated an xgboost multi-class classifier on the atlas of 35,699 RGCs subdivided into 45 molecularly distinct groups (C1-C45). Around 50% of the data was used for training and the remaining 50% was held out and used for validation. We optimized hyperparameters (e.g. tree depth, number of features, class-specific weights) to achieve a validation set accuracy of >90% across each of the 45 transcriptomic classes. This trained classifier was then used to assign a cluster label for each of the 105 transcriptomes profiled in this study. We assigned a transcriptomic label to each RGC if a minimum of 15% of trees in the forest voted on the majority decision. This choice of voting margin was >6x higher than the random threshold of 2.3%, based on the fact that there are 45 classes. The correspondences between functional and transcriptomic labels were visualized as confusion matrices.

### UMAP and cross-modality neighborhood comparisons

The functional input data to the UMAP algorithm was a linearized version of the full matrix of the PSTH for each cell across spot sizes (as in **Figure 2**). We used a MATLAB implementation of UMAP (https://www.mathworks.com/matlabcentral/fileexchange/71902) supervised by the RGC type labels for the data set of 1859 cells. The input to the UMAP algorithm for morphology was the unnormalized stratification profile for each RGC from the Eyewire museum (381 cells) supervised by the labels in the museum. Although no attempt was made to capture details of the *en face* morphological characteristics of each cell, the unnormalized stratification data allowed the algorithm to use information about total dendritic length. The input to the UMAP algorithm for transcriptomic space was a vector of gene expression values for RGC-type-selective genes from the published dataset (~35,699 cells) as described in (Tran et al., 2019).

We measured similarities between the three UMAP spaces (function, morphology, and genetics) by comparing nearest neighbors between spaces. For each RGC type in which we established a match between the two spaces being compared, we measured the fractional overlap between the nearest neighbors in the first space and those in the second space (matching types / neighborhood size). The analysis was repeated for neighborhood sizes from 2 - 12. To assess the statistics of the measured overlap values, we created a bootstrap distribution by randomly shuffling the cluster identities in one of the spaces. Data in **Figure 7D–F** are z scores with respect to this bootstrap distribution which was Gaussian.

