## Supplemental Figures for "Unified classification of mouse retinal ganglion cells using function, morphology, and gene expression"

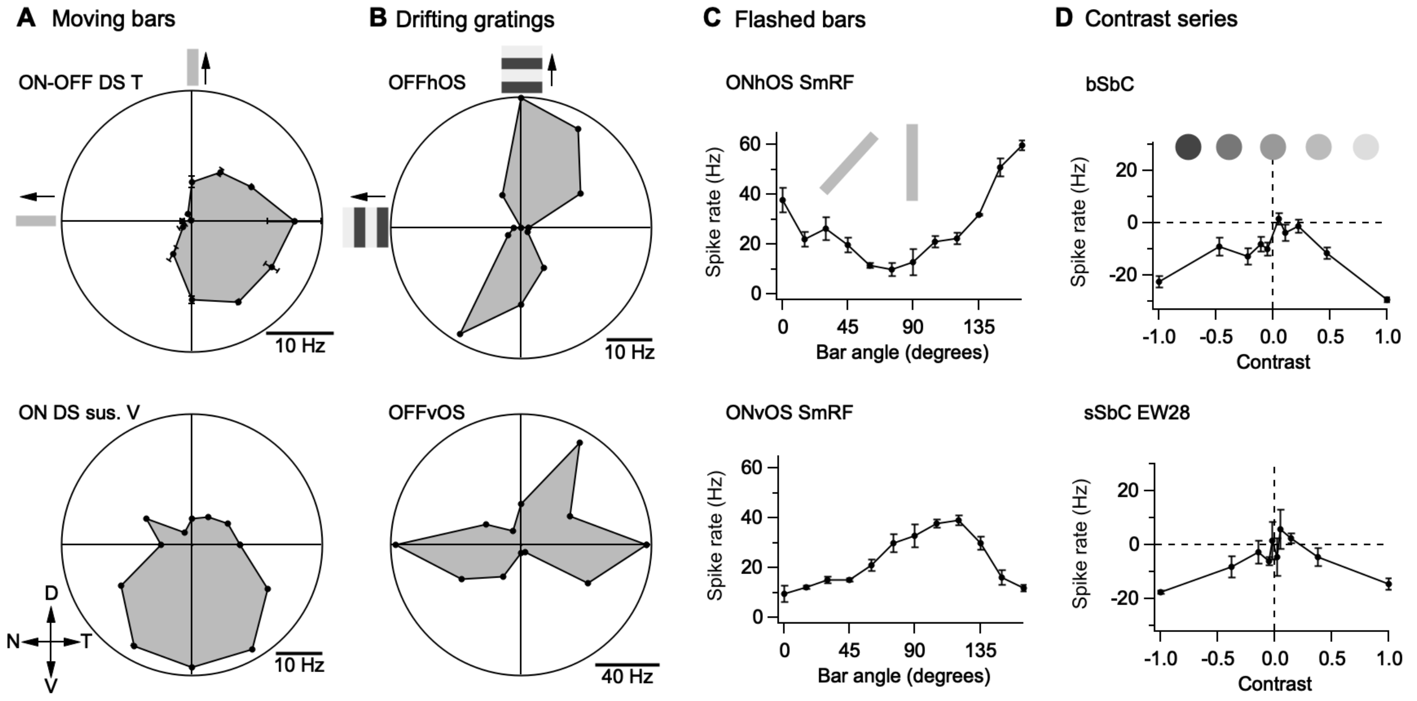


**Supplemental Figure 1. Additional stimuli for probing feature selectivity of RGCs.**

(**A**) Polar plot of spike responses to moving bar stimuli for an ON-OFF DS Temporal RGC (top) and an ON DS sustained Ventral RGC (bottom). Arrows in the lower left corner show dorsal (D), ventral (V), nasal (N), and temporal (T) directions on the retina. See **Methods** for parameters pertaining to each of the visual stimuli.

(**B**) Polar plot of spike responses to drifting grating stimuli for Horizontal (top) and Vertical (bottom) OFF OS RGCs. Note, angles on polar plots correspond to grating movement direction which is orthogonal to bar orientation.

(**C**) Spike responses for bars flashed at different orientations for a Horizontal (top) and Vertical (bottom) ON OS SmRF RGC.

(**D**) Baseline subtracted spike responses for contrast series in two different subtypes of suppress-by-contrast RGCs, the bSbC (top), and the sSbC EW28 (bottom).

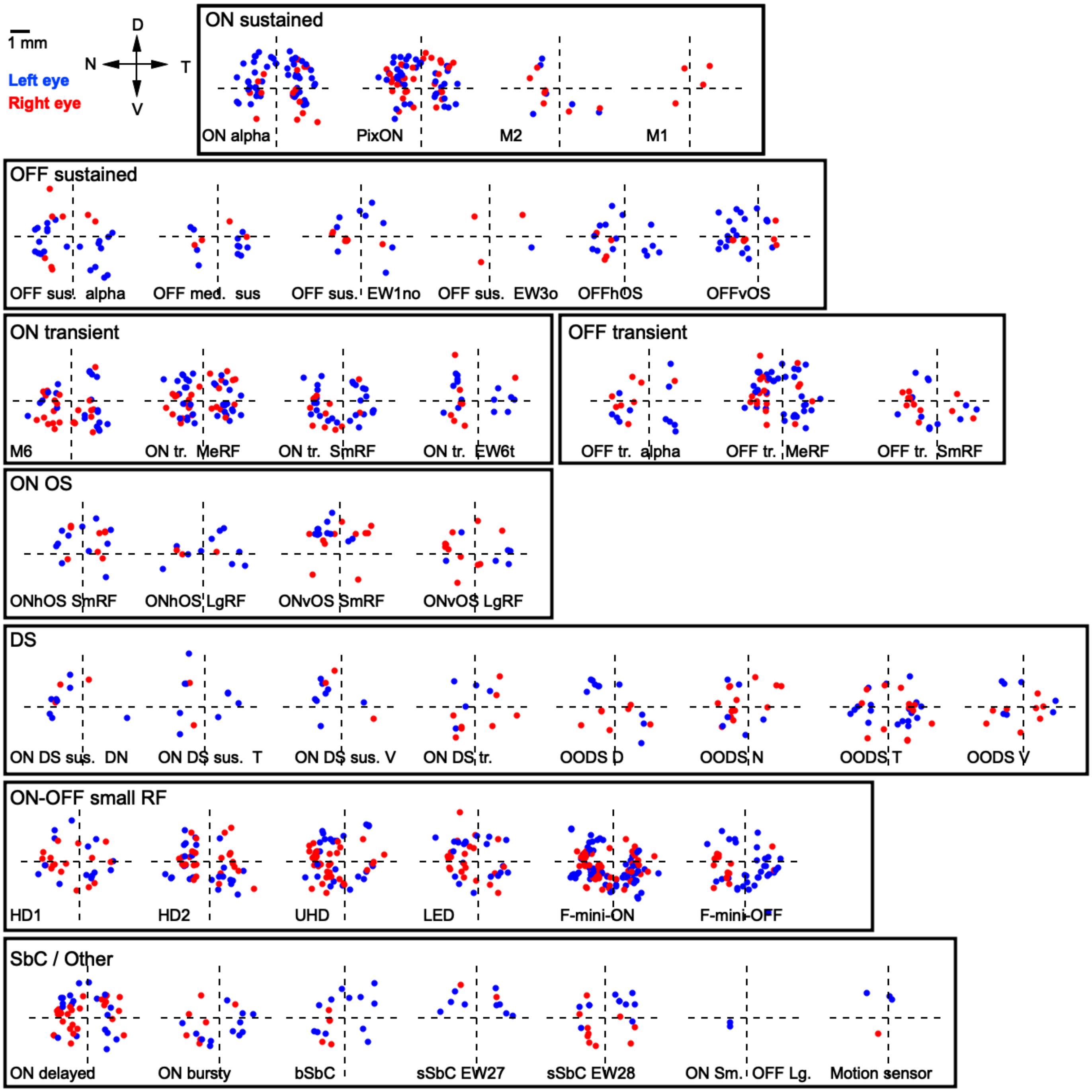


**Supplemental Figure 2. Position map of physiologically-typed RGCs.**
The location of physiologically-typed RGCs are plotted in absolute retinal space, with (0,0) representing the optic nerve. Typology was consistent across retinal space and from both left (blue) and right (red) eyes.

| RGC type | N | NT bias | p | DV bias | p |
|---|---|---|---|---|---|
| bSbc | 17 | N | 0.3719 | V | 0.4504 |
| F-mini-OFF | 55 | T | 0.0673 | V | 0.148 |
| F-mini-ON | 126 | N | 0.5344 | V | 0 |
| HD1 | 30 | T | 0.5272 | V | 0.252 |
| HD2 | 58 | N | 0.5046 | V | 0.0248 |
| LED | 42 | N | 0.0882 | D | 0.5227 |
| M1 | 5 | T | 0.3597 | D | 0.2126 |
| M2 | 13 | N | 0.298 | V | 0.1155 |
| M6 | 50 | T | 0.0551 | V | 0.1087 |
| Motion sensor | 4 | T | 0.5546 | D | 0.336 |
| OFFhOS | 19 | N | 0.121 | V | 0.2797 |
| OFF med sus | 15 | T | 0.1302 | V | 0.1306 |
| OFF sus EW1mo | 15 | N | 0.1684 | V | 0.2598 |
| OFF sus EW3o | 4 | T | 0.5598 | V | 0.6702 |
| OFF sus alpha | 29 | N | 0.3878 | V | 0.1059 |
| OFF tr alpha | 19 | N | 0.4144 | V | 0.1466 |
| OFF trr MeRF | 56 | T | 0.5119 | D | 0.0595 |
| OFF tr SmRF | 25 | N | 0.1007 | V | 0.1728 |
| OFFvOS | 29 | N | 0.0334 | V | 0.2921 |
| ON DS sus DN | 11 | N | 0.0879 | D | 0.1329 |
| ON DS sus T | 14 | T | 0.3718 | D | 0.1053 |
| ON DS sus V | 11 | N | 0.0923 | D | 0.0379 |
| ON DS tr | 17 | T | 0.4262 | D | 0.3561 |
| ON alpha | 81 | T | 0.0008 | D | 0.0198 |
| ON bursty | 20 | T | 0.4818 | V | 0.1041 |
| ON delayed | 49 | N | 0.3735 | D | 0.0157 |
| ONhOS LgRF | 13 | T | 0.2761 | D | 0.1547 |
| ONhOS SmRF | 20 | T | 0.1743 | D | 0.0071 |
| ON sm OFF lg | 2 | N | 0.3349 | V | 0.2353 |
| ON tr EW6t | 23 | N | 0.1719 | D | 0.2519 |
| ON tr MeRF | 59 | T | 0.2415 | V | 0.407 |
| ON tr SmRF | 45 | T | 0.2136 | V | 0.4265 |
| ONvOS LgRF | 17 | T | 0.4274 | D | 0.3615 |
| ONvOS SmRF | 21 | N | 0.448 | D | 0.0004 |
| OODS D | 18 | T | 0.5201 | V | 0.5395 |
| OODS N | 20 | N | 0.1934 | V | 0.3559 |
| OODS T | 31 | T | 0.0498 | V | 0.1885 |
| OODS V | 16 | T | 0.5463 | V | 0.3617 |
| PixON | 78 | T | 0.5408 | D | 0.0001 |
| sSbC EW 27 | 10 | T | 0.2033 | D | 0.0009 |
| sSbc EW28 | 23 | T | 0.2183 | D | 0.2342 |
| UHD | 72 | N | 0.0003 | V | 0.4558 |

**Supplemental Table 1. Statistical tests of positional bias in RGC type distributions.** Columns list the number of cells of each type with position information, whether there was a bias for more cells to be found on the nasal (N) or temporal (T) half of the retina or the dorsal (D) or ventral (V) half of the retina. P values are for one-sided Student's t-tests. Yellow highlights indicate significance below the bonferroni corrected threshold of $2.9 \times 10^{-4}$. Orange highlights indicate potentially significant trends. Note: the F-mini-ON bias in the ventral retina was heavily influenced by experimental sampling bias based on the results of (Rousso et al. 2016).

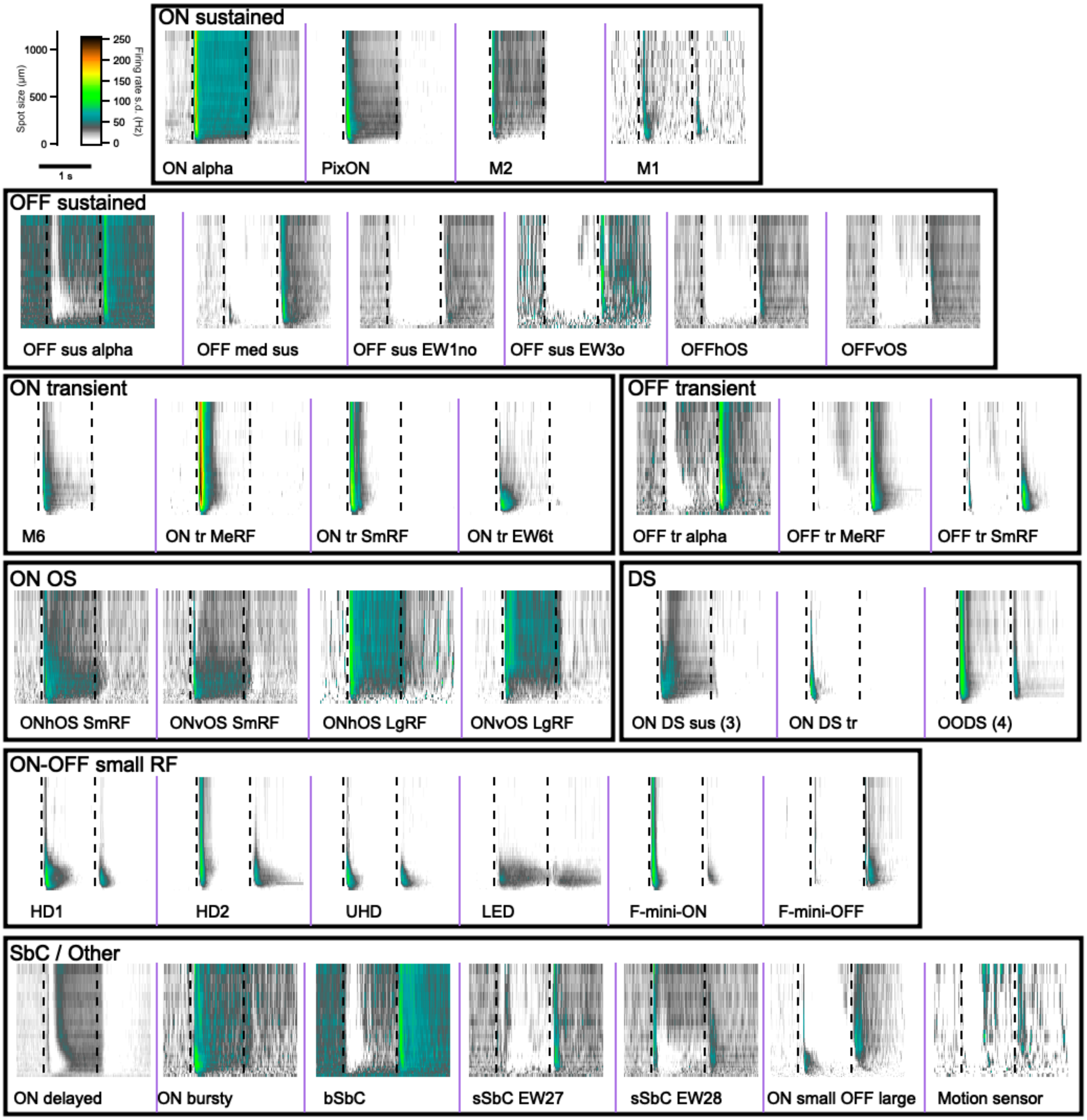


**Supplemental Figure 3. Standard deviation of light responses within each functional RGC type.**

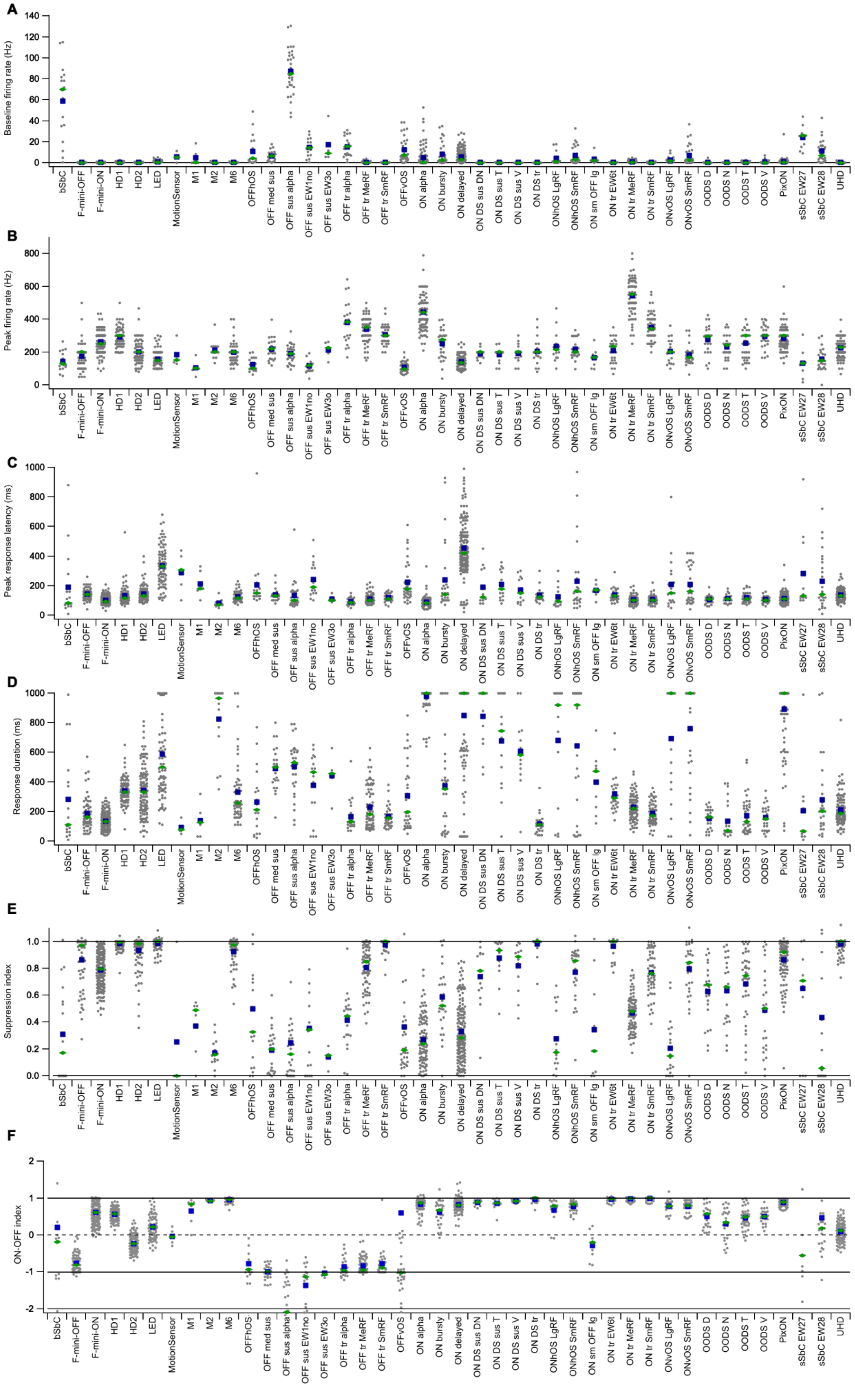


**Supplemental Figure 4. Distributions of six response metrics across RGC types.** Points are individual cells. Blue squares are means and green diamonds are medians for each metric.

| RGC type | Physiology (N) | Validated (N) | % | Confocal image (N) | 2P image (N) | Transgenic (N) | Vclamp (N) | Soma size (N) | DS (N) | OS (N) |
|---|---|---|---|---|---|---|---|---|---|---|
| bSbc | 17 | 8 | 47.1 | 5 | 0 | 0 | 3 | 0 | 0 | 0 |
| F-mini-OFF | 65 | 20 | 30.8 | 2 | 1 | 13 | 7 | 0 | 0 | 0 |
| F-mini-ON | 176 | 32 | 18.2 | 3 | 4 | 4 | 24 | 0 | 0 | 0 |
| HD1 | 94 | 12 | 12.8 | 2 | 0 | 0 | 9 | 0 | 0 | 0 |
| HD2 | 108 | 12 | 11.1 | 1 | 2 | 1 | 8 | 0 | 0 | 0 |
| LED | 83 | 3 | 3.6 | 1 | 0 | 0 | 2 | 0 | 0 | 0 |
| M1 | 6 | 2 | 33.3 | 1 | 1 | 0 | 1 | 0 | 0 | 0 |
| M2 | 14 | 2 | 14.3 | 0 | 1 | 1 | 0 | 0 | 0 | 0 |
| M6 | 54 | 18 | 33.3 | 3 | 8 | 8 | 5 | 0 | 0 | 0 |
| Motion sensor | 4 | 2 | 50.0 | 1 | 0 | 0 | 0 | 0 | 0 | 0 |
| OFFhOS | 19 | 19 | 100 | 1 | 0 | 0 | 0 | 0 | 0 | 19 |
| OFF med sus | 23 | 4 | 17.4 | 1 | 3 | 0 | 2 | 0 | 0 | 0 |
| OFF sus EW1no | 18 | 2 | 11.1 | 0 | 0 | 0 | 0 | 0 | 0 | 0 |
| OFF sus EW3o | 4 | 2 | 50.0 | 1 | 1 | 0 | 0 | 0 | 0 | 0 |
| OFF sus alpha | 31 | 5 | 16.1 | 0 | 0 | 0 | 4 | 0 | 0 | 0 |
| OFF tr alpha | 22 | 19 | 86.4 | 0 | 0 | 0 | 0 | 19 | 0 | 0 |
| OFF trr MeRF | 73 | 8 | 11.0 | 4 | 2 | 0 | 2 | 0 | 0 | 0 |
| OFF tr SmRF | 32 | 5 | 15.6 | 0 | 1 | 4 | 1 | 0 | 0 | 0 |
| OFFvOS | 35 | 35 | 100 | 0 | 1 | 0 | 1 | 0 | 0 | 34 |
| ON DS sus DN | 13 | 10 | 76.9 | 0 | 0 | 0 | 0 | 0 | 10 | 0 |
| ON DS sus T | 14 | 13 | 92.9 | 1 | 0 | 0 | 0 | 0 | 13 | 0 |
| ON DS sus V | 13 | 12 | 92.3 | 1 | 0 | 0 | 0 | 0 | 11 | 0 |
| ON DS tr | 21 | 13 | 61.9 | 1 | 1 | 0 | 0 | 0 | 13 | 0 |
| ON alpha | 96 | 88 | 91.7 | 1 | 9 | 4 | 16 | 86 | 0 | 0 |
| ON bursty | 29 | 2 | 6.9 | 2 | 0 | 0 | 0 | 0 | 0 | 0 |
| ON delayed | 161 | 41 | 25.5 | 2 | 6 | 0 | 38 | 0 | 0 | 0 |
| ONhOS LgRF | 14 | 12 | 85.7 | 0 | 1 | 0 | 0 | 0 | 0 | 12 |
| ONhOS SmRF | 27 | 24 | 88.9 | 0 | 2 | 0 | 2 | 0 | 0 | 24 |
| ON sm OFF lg | 9 | 2 | 22.2 | 1 | 1 | 0 | 1 | 0 | 0 | 0 |
| ON tr EW6t | 25 | 2 | 8.0 | 2 | 0 | 0 | 0 | 0 | 0 | 0 |
| ON tr MeRF | 83 | 5 | 6.0 | 2 | 1 | 0 | 2 | 0 | 0 | 0 |
| ON tr SmRF | 57 | 6 | 10.5 | 3 | 1 | 0 | 1 | 0 | 0 | 0 |
| ONvOS LgRF | 20 | 18 | 90.0 | 4 | 0 | 0 | 0 | 0 | 0 | 18 |
| ONvOS SmRF | 31 | 30 | 96.8 | 0 | 1 | 0 | 2 | 0 | 0 | 29 |
| OODS D | 25 | 22 | 88.0 | 0 | 0 | 0 | 0 | 0 | 22 | 0 |
| OODS N | 25 | 21 | 84.0 | 0 | 0 | 0 | 1 | 0 | 21 | 0 |
| OODS T | 41 | 36 | 87.8 | 2 | 0 | 0 | 0 | 0 | 36 | 0 |
| OODS V | 25 | 22 | 88.0 | 1 | 0 | 0 | 0 | 0 | 22 | 0 |
| PixON | 105 | 30 | 28.6 | 6 | 15 | 6 | 17 | 0 | 0 | 0 |
| sSbC EW 27 | 11 | 2 | 18.2 | 2 | 0 | 0 | 0 | 0 | 0 | 0 |
| sSbc EW28 | 24 | 1 | 4.2 | 1 | 0 | 0 | 0 | 0 | 0 | 1 |
| UHD | 112 | 12 | 10.7 | 2 | 2 | 1 | 7 | 0 | 0 | 0 |

**Supplemental Table 2. External validation of RGC typology.** For each RGC type, columns list the number of cells with one or more validation datasets separate from the core spots typology data used by the ML classifier. Following the column listed the percent validation, numbers of cells are listed by each validation data type.

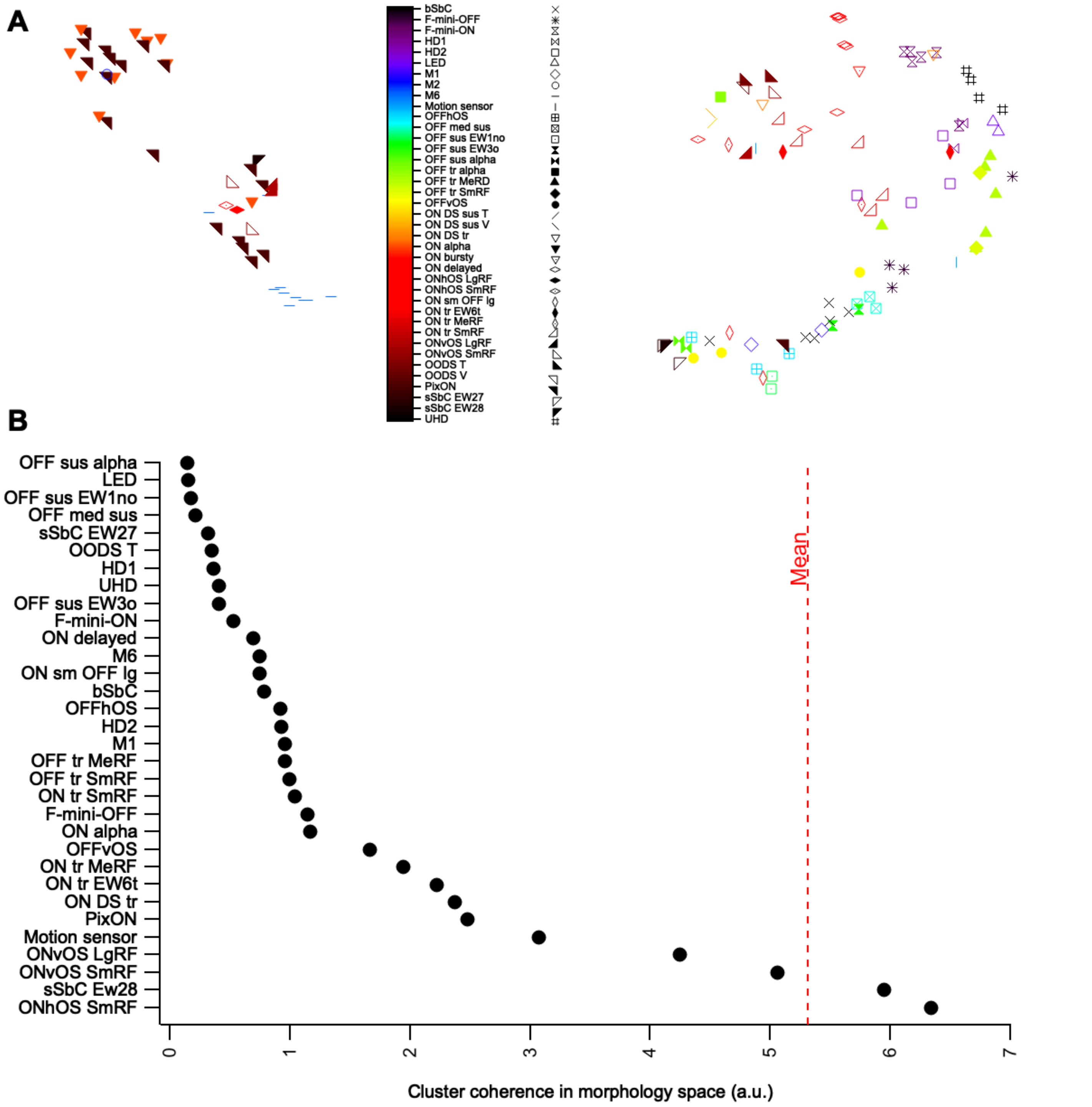


**Supplemental Figure 5. Concordance between functional and morphological clusters at the level of single cells.**
(**A**) UMAP of morphology dataset (n = 136 RGCs) using all morphological parameters described in Methods.
(**B**) The X-axis shows the mean pairwise distance in this UMAP space between cells belonging to each functionally-defined type. Red dashed line is the mean pairwise distance between all cells. RGC types are ordered by cluster coherence.

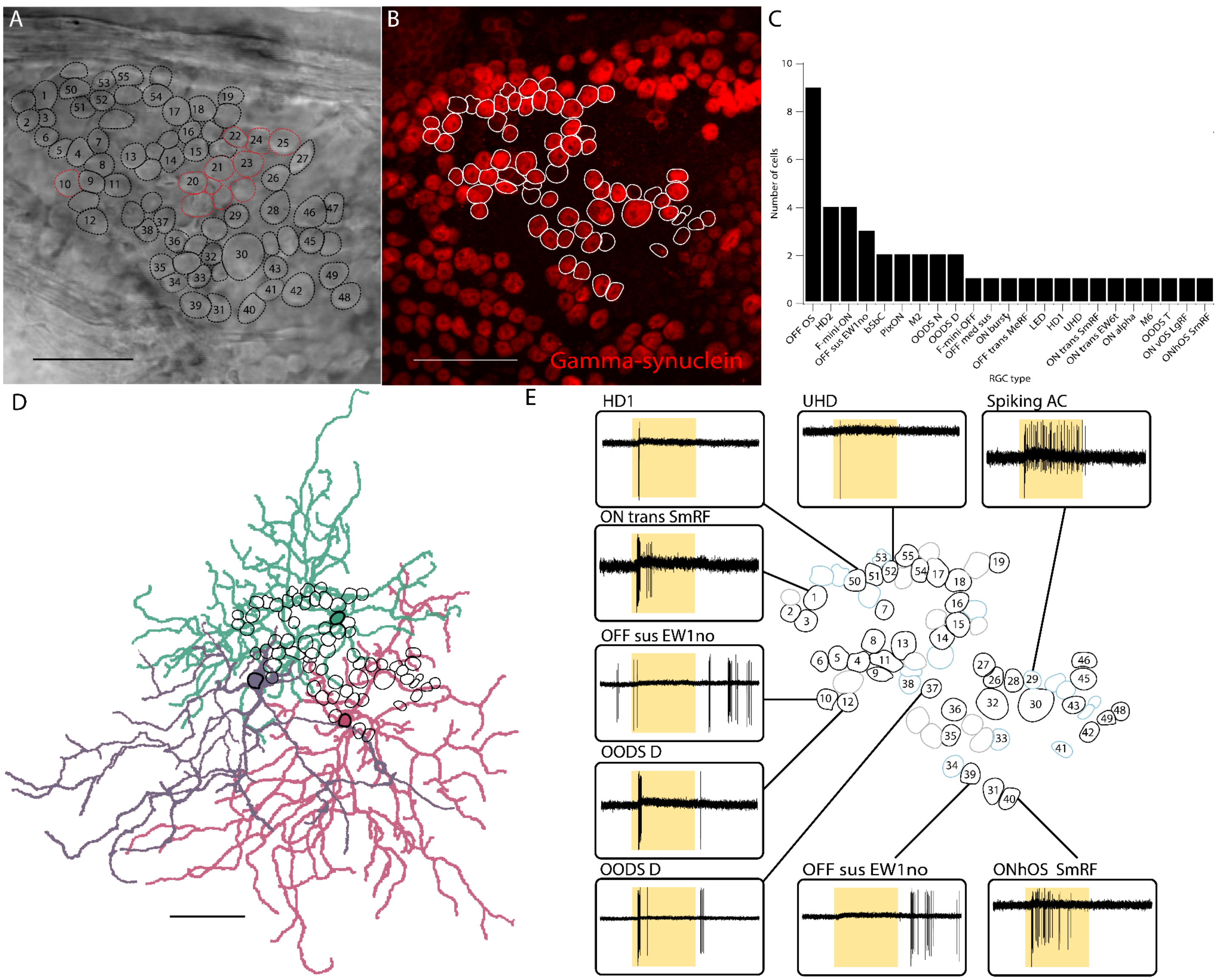


**Supplemental Figure 6. Classifying most RGCs in a retinal region**

(**A**) Image with recorded cells outlined. Numbered cells had spike responses to our standard stimuli. Red outlines indicate cells for which the soma was not visible in the fixed tissue.

(**B**) Confocal image of the same region stained with an antibody against the pan-RGC marker gamma-synuclein.

(**C**) Histogram of RGC types identified in the region in (A,B).

(**D**) Reconstructions of 3 dye-filled RGC's of type 1no from this region. Soma outlines from (B) are included.

(**E**) Examples of light responses from some of the recorded cells in this region. Cell outlines are colored according to the outcome of classification and gamma-synuclein staining. Black outlines indicate cells identified as RGCs that were synuclein-gamma positive (n = 42). Cyan outlines indicate gamma-synuclein negative cells that were either identified as spiking amacrine cells (n = 6, numbered) or non-spiking (n = 10, not numbered). Gray outlines indicate gamma-synuclein positive cells for which we were unable to elicit spikes (n = 12).

Scale bars in (**A**), (**B**), and (**D**) indicate 50 μm.

| RGC type | Eyewire type | Tran et al. (2019) cluster | Baden et al. (2016) group | Baden et al. (2016) match confidence | References |
|---|---|---|---|---|---|
| ON alpha | 8w | C43 | 24 | high | (Krieger et al., 2017) |
| PixON | 9n | | 22a,b | low | (Johnson et al., 2018) |
| M2 | 9w | | | | (Schmidt and Kofuji, 2009) |
| M1 | 1ws | C40 | | | (Berson et al., 2010; Do et al., 2009) |
| OFF sus alpha | 1wt | | 5a,b,c | high | (Krieger et al., 2017) |
| OFF med sus | 2i | | 3 | low | |
| OFF sus EW1no | 1no | | 4a | medium | |
| OFF sus EW3o | 3o | | 4b | medium | |
| OFFhOS | 2aw | C9 | 1, 14 | medium | (Nath and Schwartz, 2017) |
| OFFvOS | 2aw | C5 | 1, 2, 6, 14 | medium | (Kim et al., 2008; Nath and Schwartz, 2017) |
| M6 | 91 | | 20 | low | (Quattrochi et al., 2019) |
| ON tr MeRF | 6sw | | 18a | low | |
| ON tr SmRF | 6sn | C30 | 18b | low | |
| ON tr EW6t | 6t | | 21 | low | |
| OFF tr alpha | 4ow | C45 | 8a,b | high | (Krieger et al., 2017) |
| OFF tr MeRF | 4on | | 9 | low | |
| OFF tr SmRF | 4i | C21 | 9 | low | |
| ONhOS SmRF | 82wi | C27 | 17a,b,c | medium | (Nath and Schwartz, 2016) |
| ONhOS LgRF | 82n, 82wo | C36 | 30 | medium | |
| ONvOS SmRF | 72 | C38 | 17a,b,c | medium | (Nath and Schwartz, 2016) |
| ONvOS LgRF | 81o, 81i | | 30 | medium | |
| ON DS sus DN | 7iv | C10 | 25 | high | (Estevez et al., 2013) |
| ON DS sus T | 7ir | C10 | 26,29 | high | (Estevez et al., 2013) |
| ON DS sus V | 7id | C10 | 26,29 | high | (Estevez et al., 2013) |
| ON DS tr | 7o | | 16 | medium | (Gauvain and Murphy, 2015) |
| OODS D | 37v | C16 | 12a,b,13 | high | (Kay et al., 2011) |
| OODS T | 37r | C24 | 12a,b,13 | high | (Kay et al., 2011) |
| OODS V | 37d | | 12a,b,13 | high | (Trenholm et al., 2011) |
| OODS N | 37c | | 12a,b,13 | high | (Kay et al., 2011) |
| HD1 | 5si | C13 | 10,11a,b | low | (Jacoby and Schwartz, 2017) |
| HD2 | 5so | C6 | 10,11a,b | low | (Jacoby and Schwartz, 2017) |
| UHD | 5ti | C2 | 10,11a,b | low | (Jacoby and Schwartz, 2017) |
| LED | 51 | C11 | 10,11a,b | low | (Jacoby and Schwartz, 2017) |
| F-mini-ON | 63 | C3 | 10,11a,b | low | (Cooler and Schwartz, 2020; Rousso et al., 2016) |
| F-mini-OFF | 2an | | | | (Cooler and Schwartz, 2020; Rousso et al., 2016) |
| ON delayed | 73 | C14 | 27,28a,b | low | (Mani and Schwartz, 2017) |
| ON bursty | 3i | C18 | 27 | low | |
| bSbC | 2o | C25 | 32a,b,c | medium | (Wienbar and Schwartz, 2021) |
| sSbC EW27 | 27 | C26 | 31a,b,c,d,e | low | (Jacoby et al., 2015) |
| sSbC EW28 | 28 | | 31a,b,c,d,e | low | (Jacoby et al., 2015) |
| ON sm OFF lg | 1ni | | | | |
| Motion sensor | 5to | | | | |

**Supplemental Table 3. Mouse RGC types and their alignment to previous classifications.**

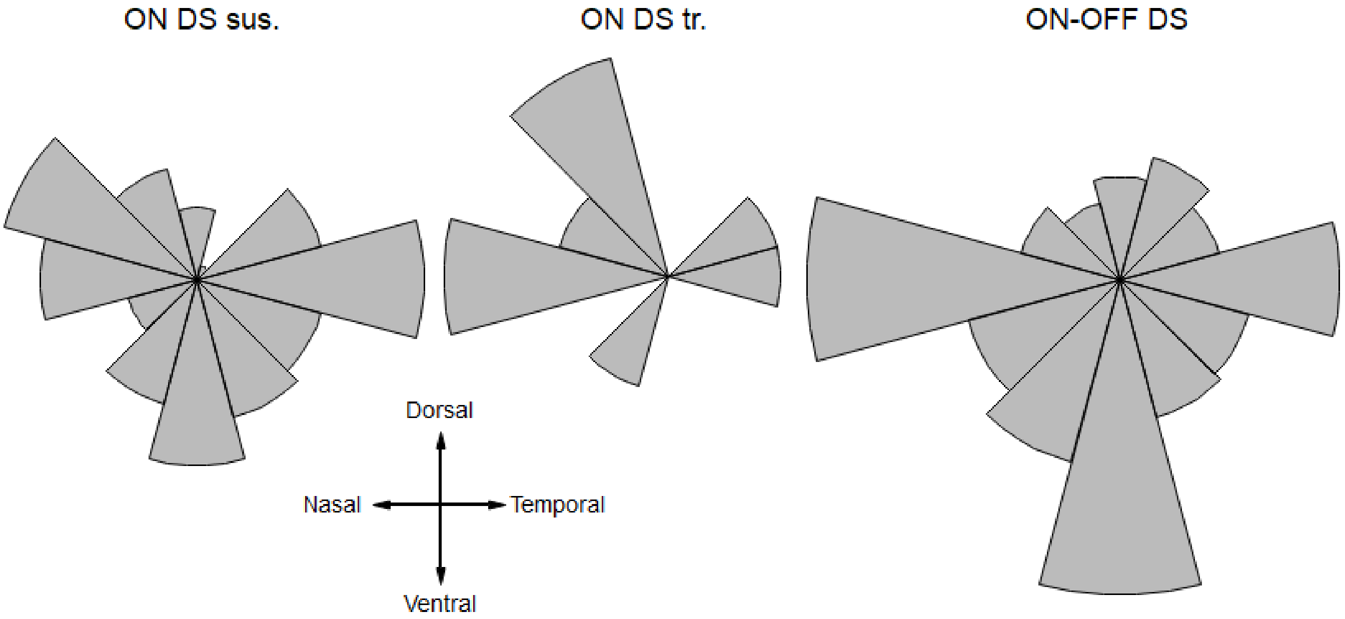


**Supplemental Figure 7. DS rose plots**
Histograms showing the direction preferences of each type of DS using the optic nerve and a ventral cut to orient retinas (n = 110 ON DS sus., 8 ON DS tr., 252 OODS). We observe three major categories of ON DS sus. (Dorsonasal, Temporal, and Ventronasal). ON DS tr. were rarely observed, with a tendency to prefer dorsonasal movement. There are known to be four major categories of OODS cells preferring each cardinal direction; while we observe fewer OODS-D than expected, all four types are seen in our dataset.
