## Supplemental Data for "Unified classification of mouse retinal ganglion cells using function, morphology, and gene expression"

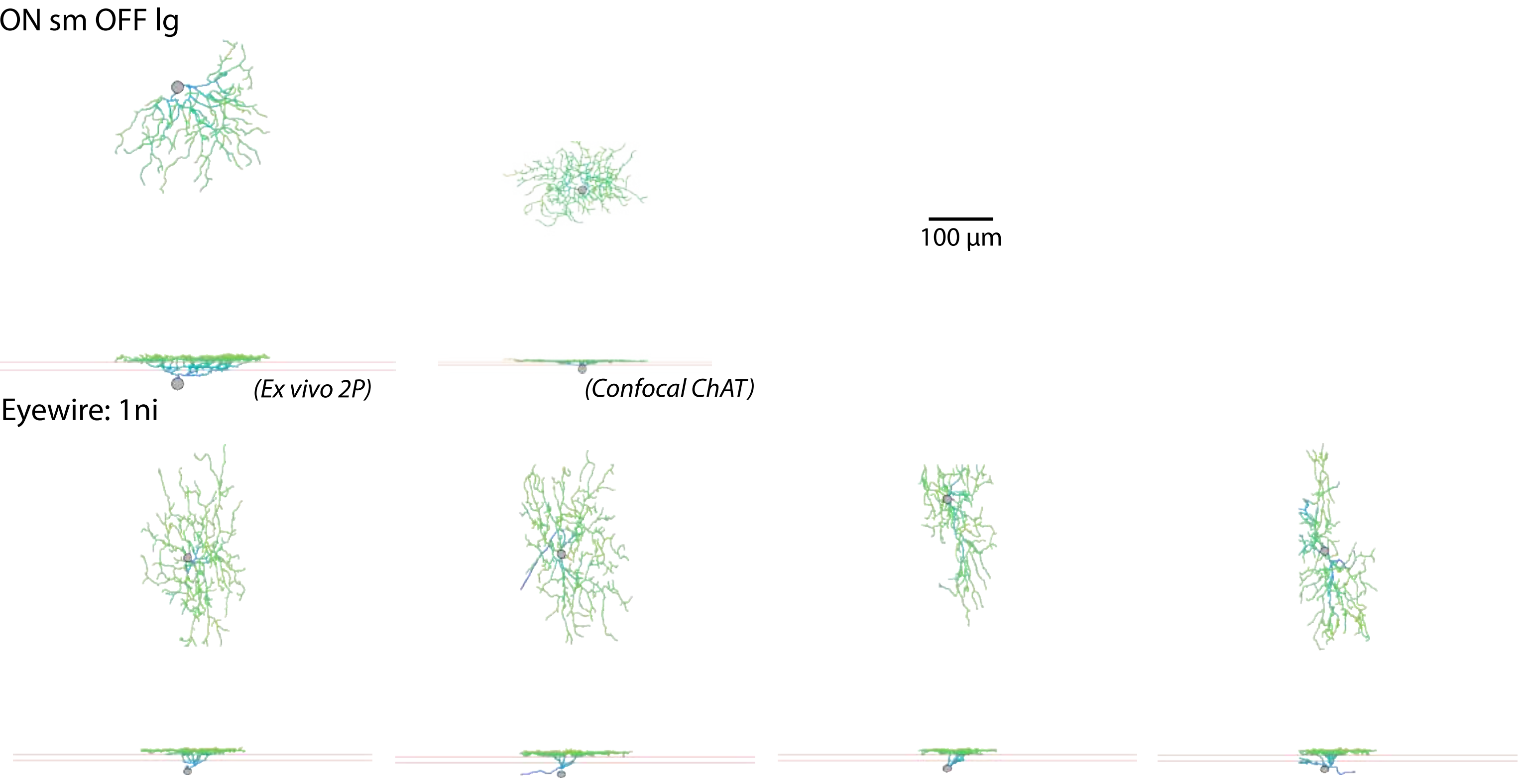
ON sm OFF lg
100 µm
(Ex vivo 2P)
(Confocal ChAT)
Eyewire: 1ni

### OFF sus EW1no

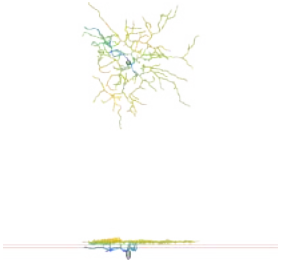

(Confocal ChAT)

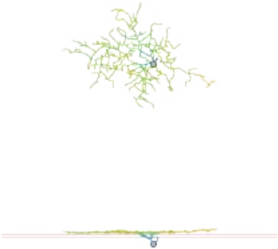

(Confocal ChAT)

### Eyewire: 1no

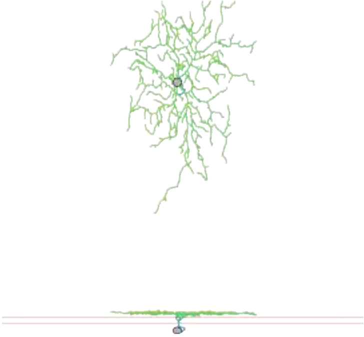

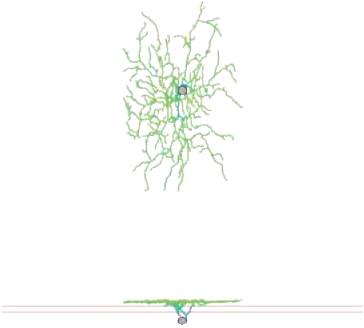

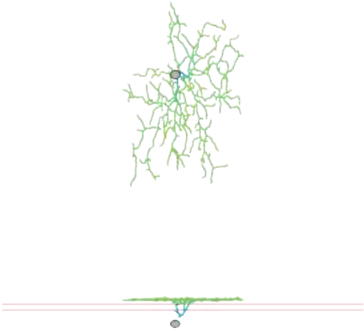

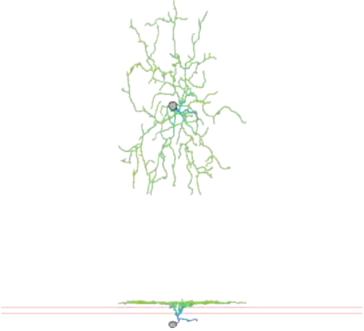

M1

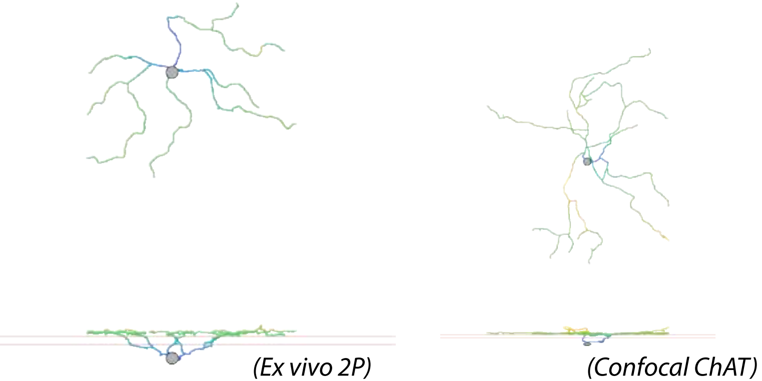


Eyewire: 1ws

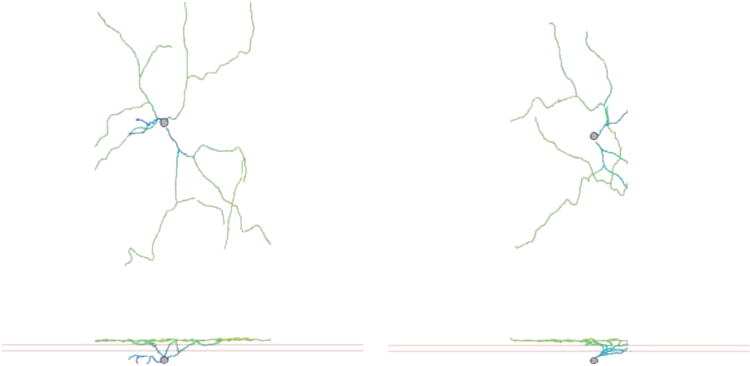

OFF sus alpha

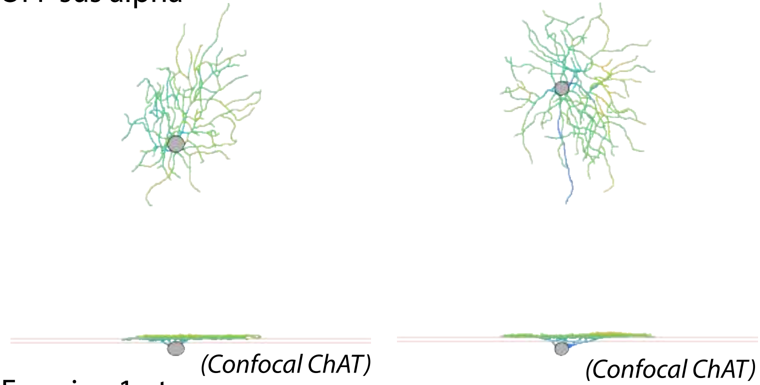


Eyewire: 1wt

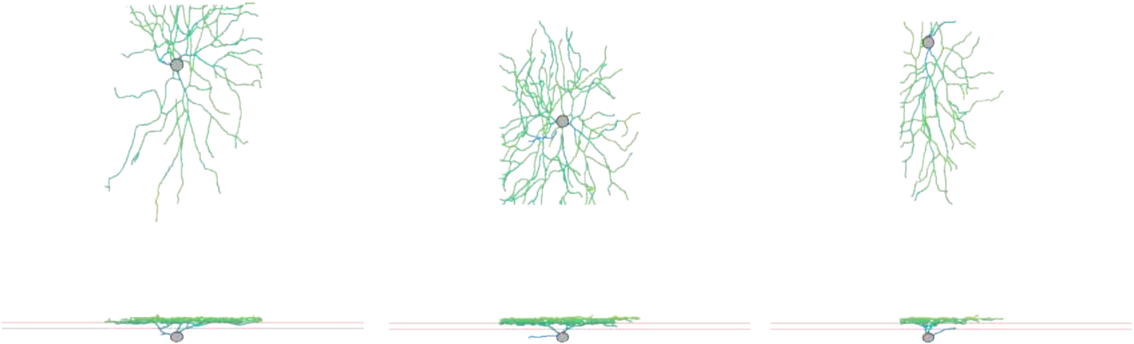

sSbC EW27

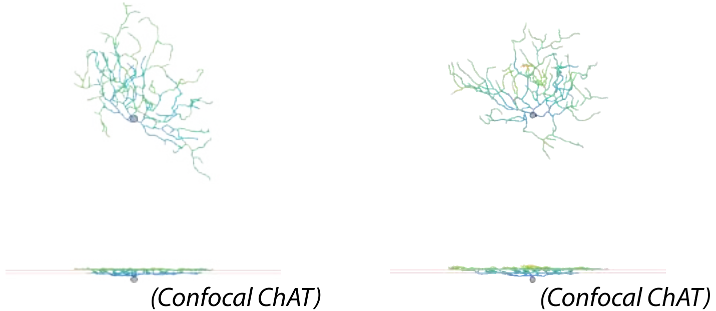


Eyewire: 27

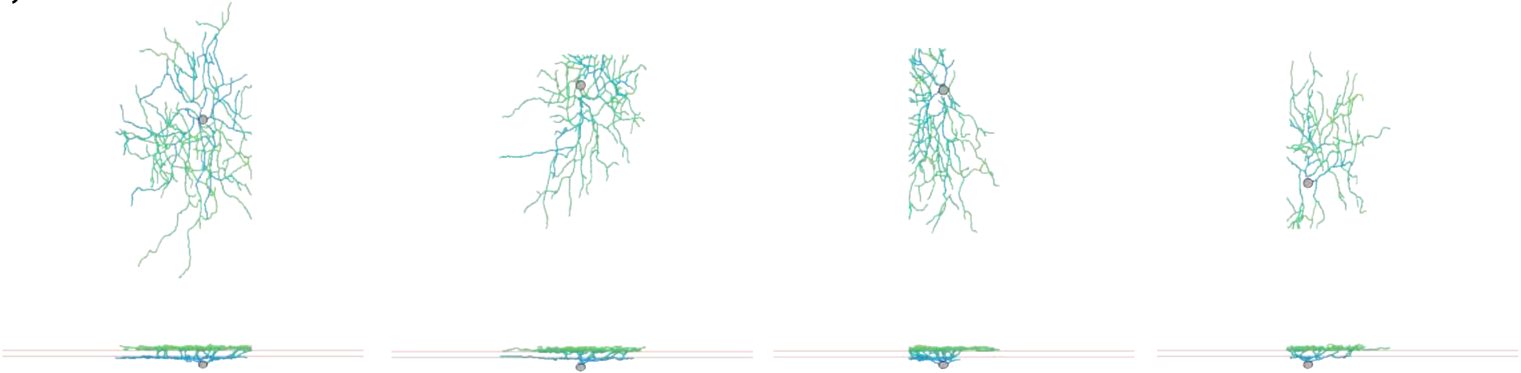

sSbC EW28

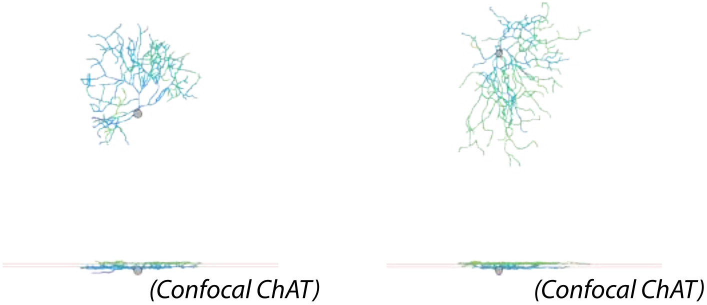


Eyewire: 28

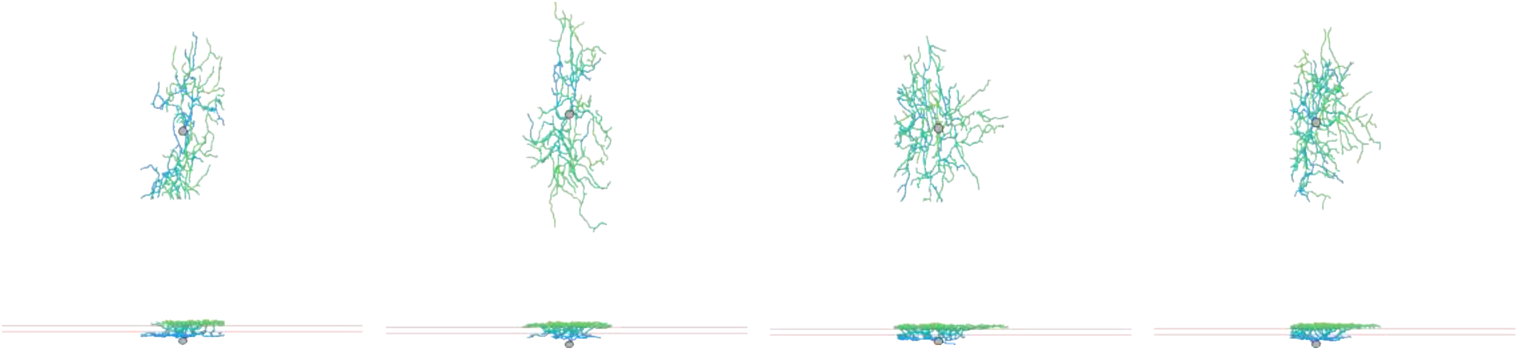

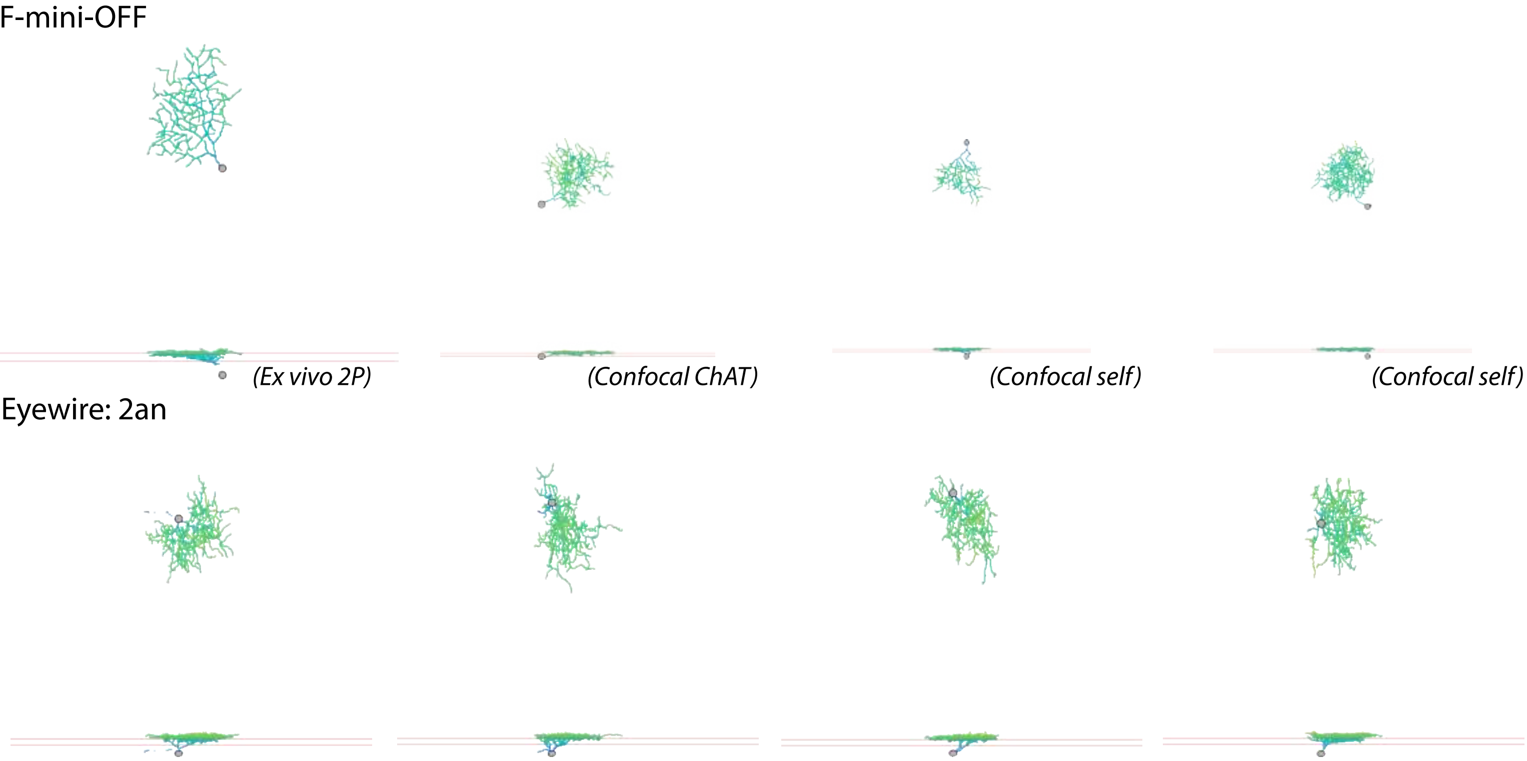
F-mini-OFF
(Ex vivo 2P)
(Confocal ChAT)
(Confocal self)
(Confocal self)
Eyewire: 2an

OFFhOS

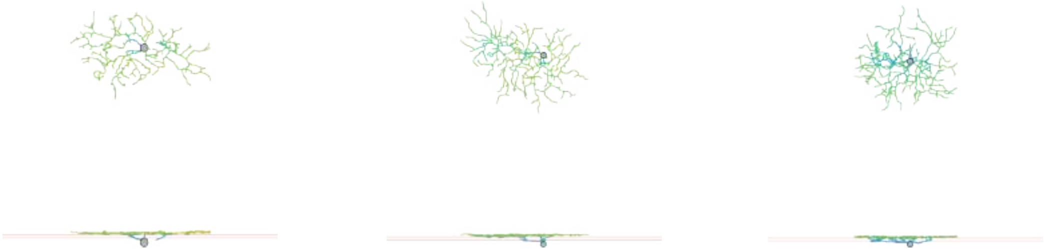

*(Confocal ChAT)* *(Confocal ChAT)* *(Confocal ChAT)*

Eyewire: 2aw

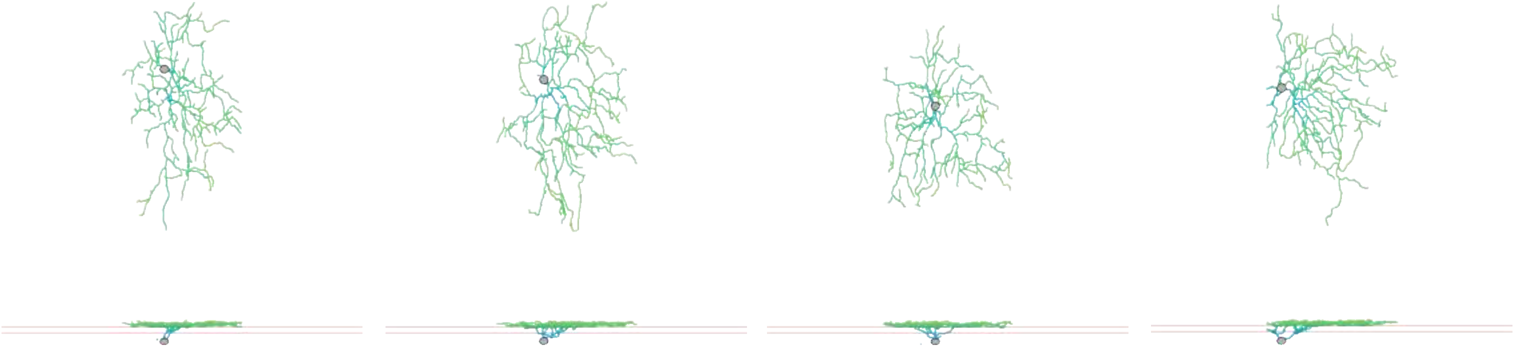

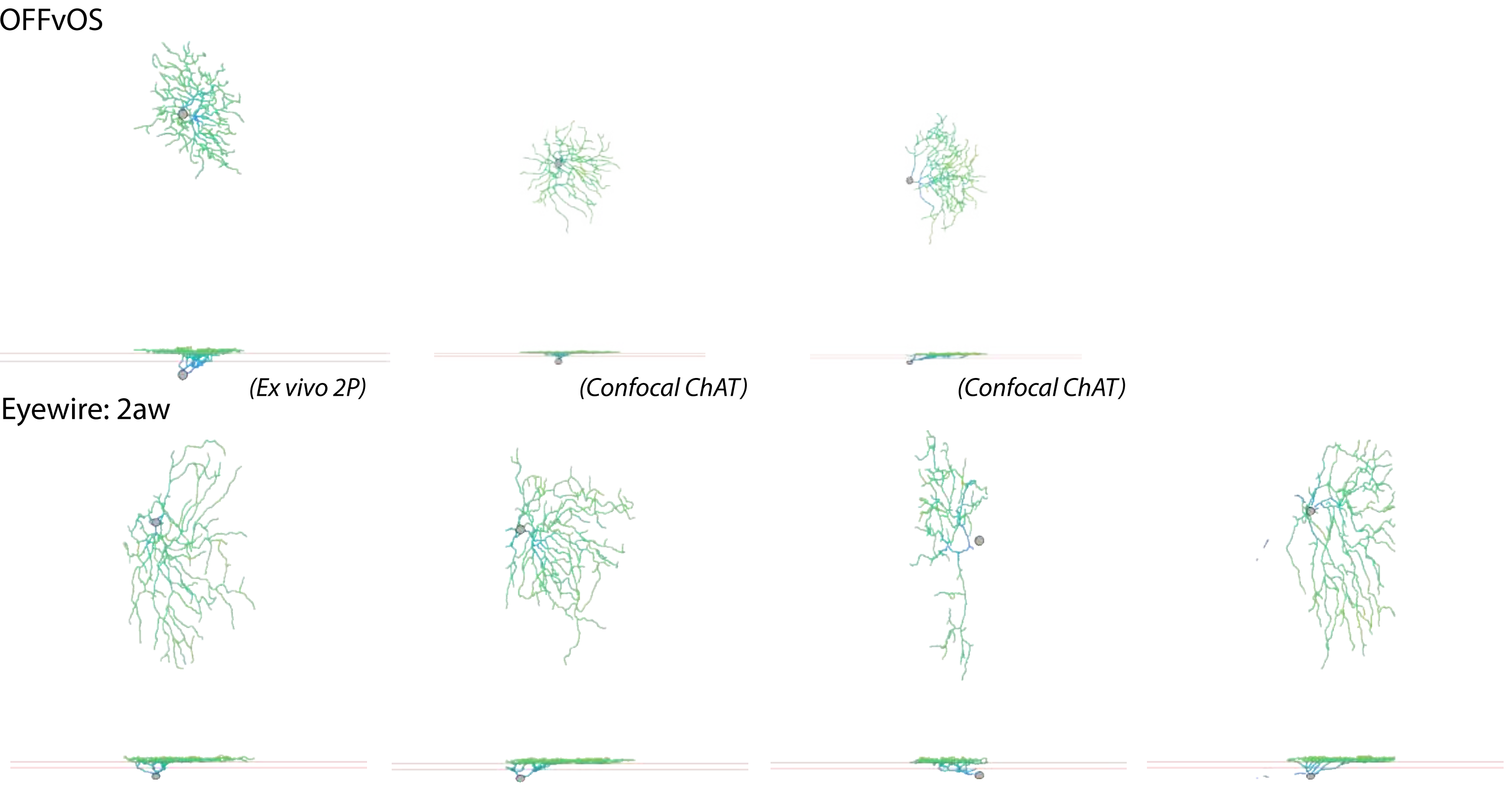
OFFvOS
(Ex vivo 2P)
(Confocal ChAT)
(Confocal ChAT)
Eyewire: 2aw

OFF med sus

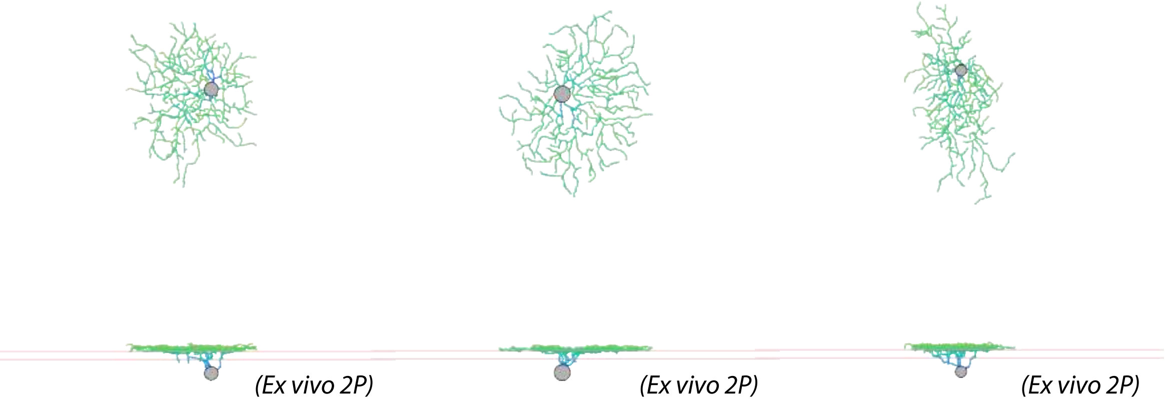


Eyewire: 2i

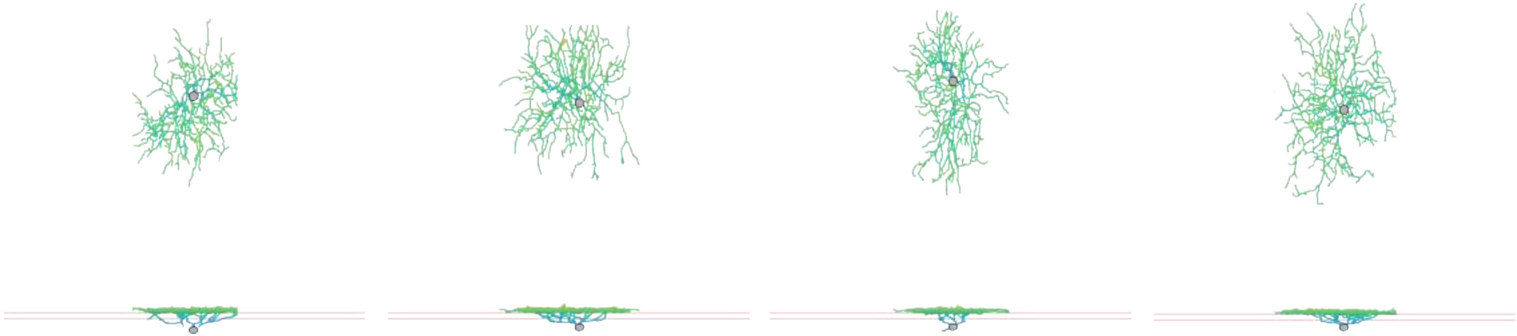

bSbC

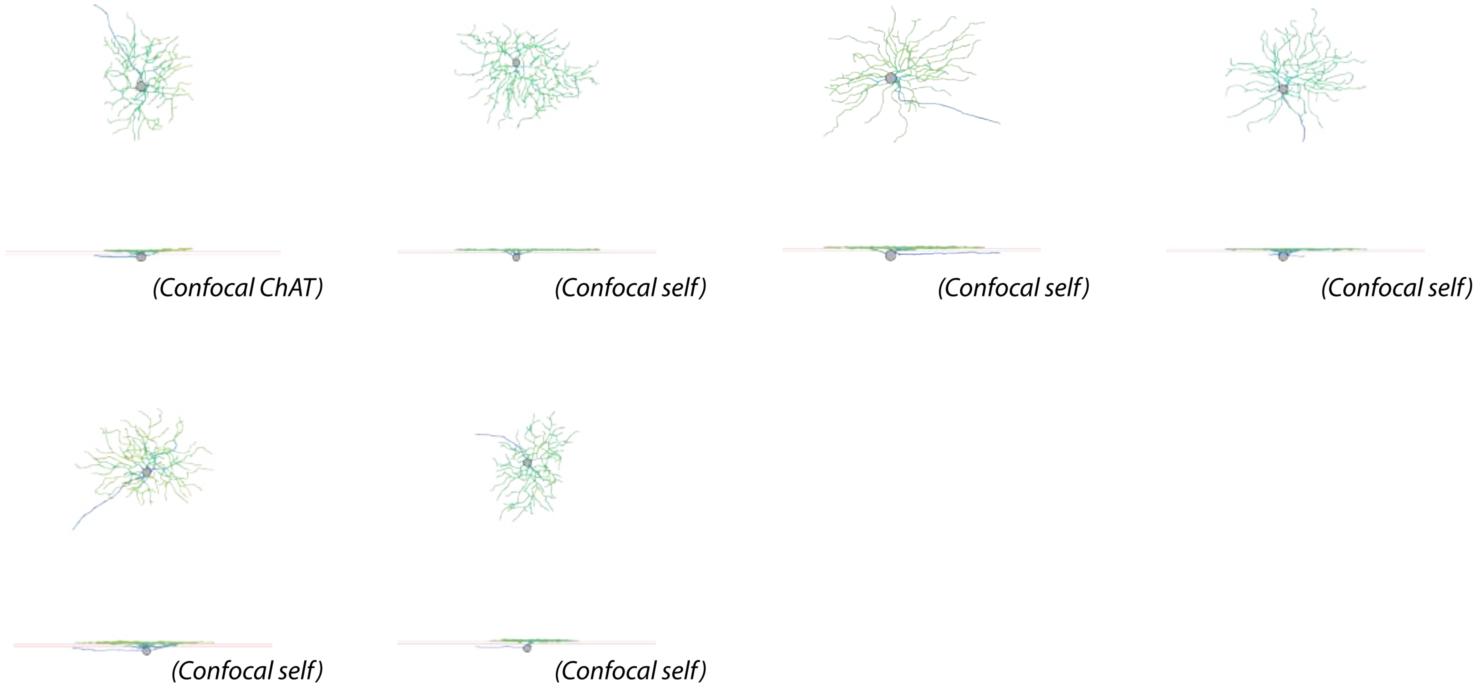


Eyewire: 2o

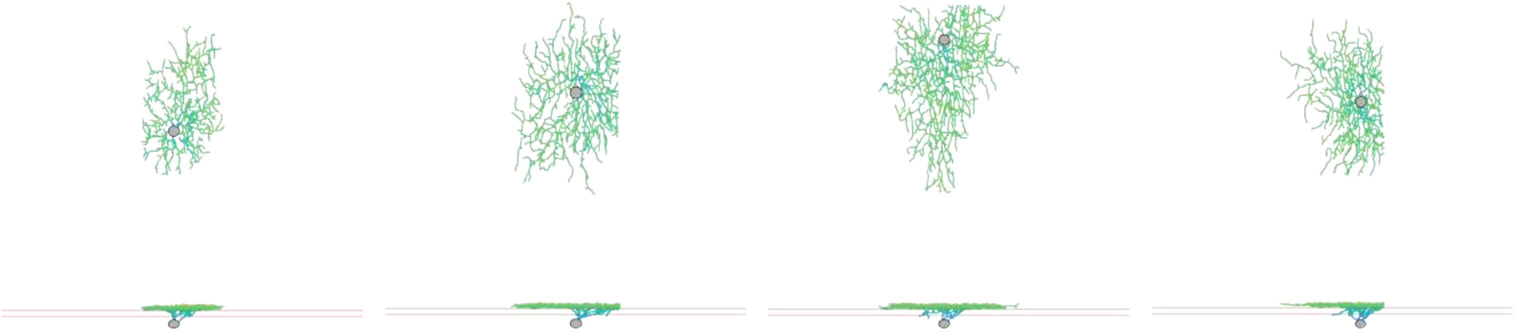

OODS V

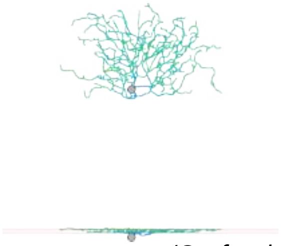

*(Confocal self)*

Eyewire: 37d

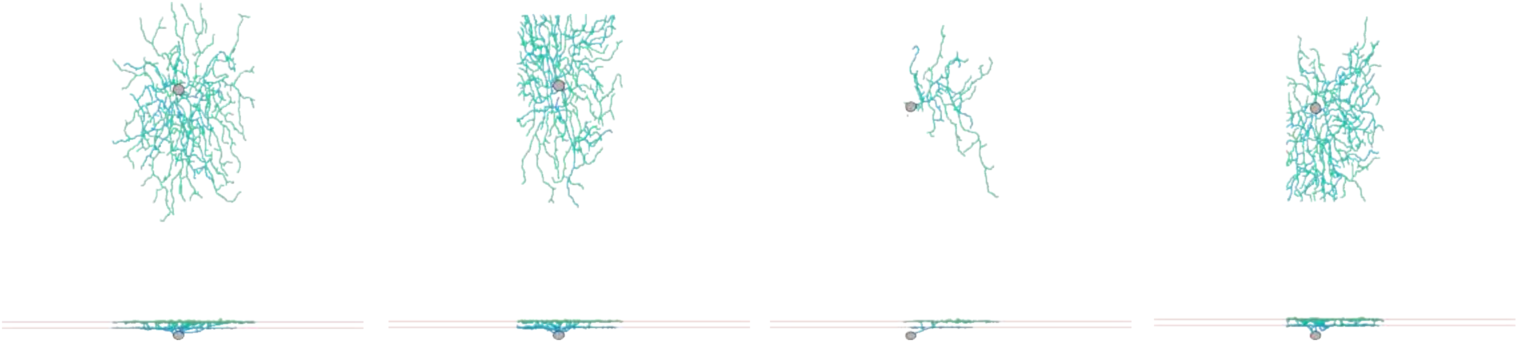

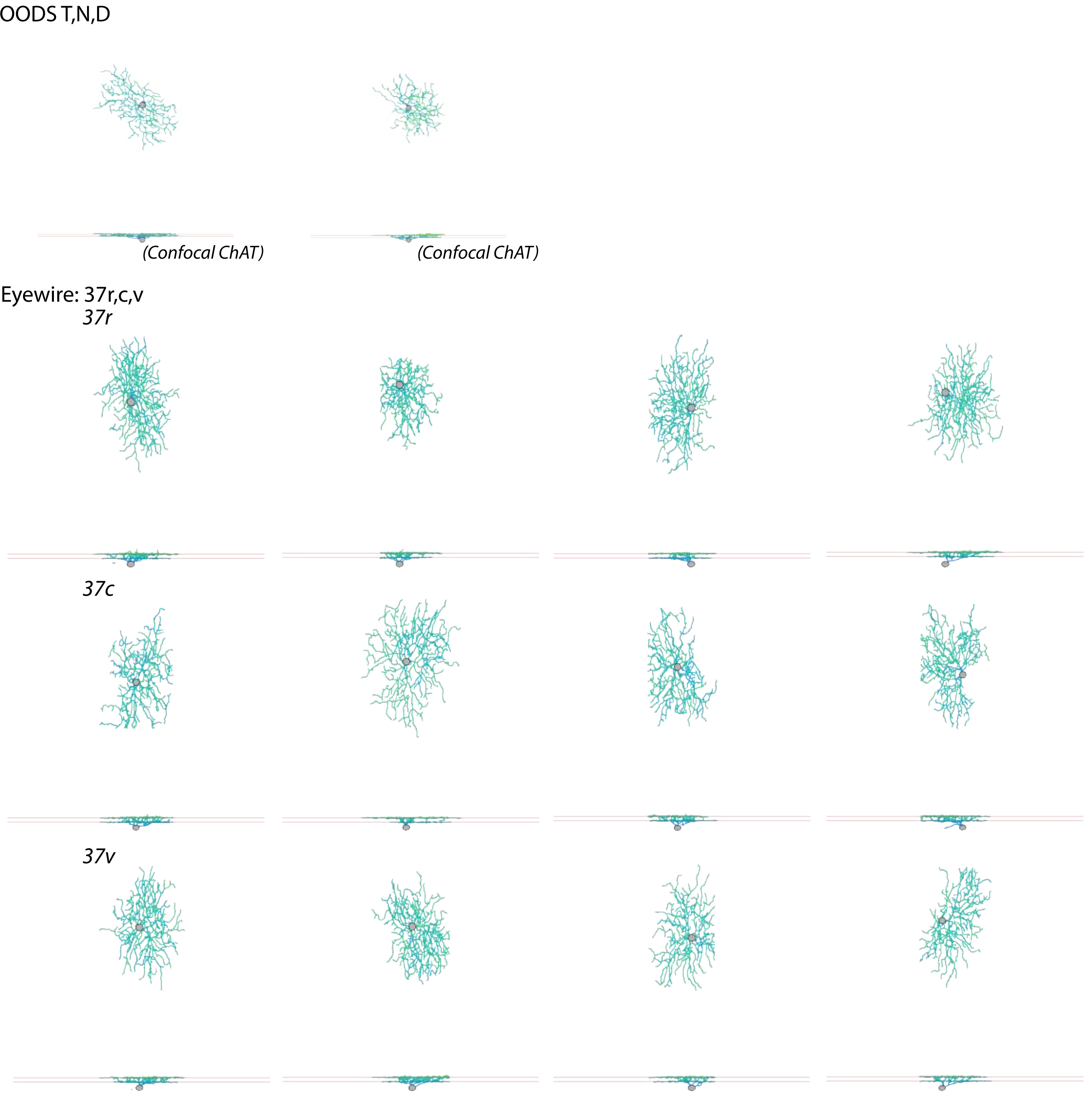
OODS T,N,D
(Confocal ChAT)
(Confocal ChAT)
Eyewire: 37r,c,v
37r
37c
37v

ON bursty

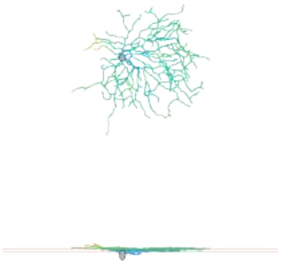

(Confocal ChAT)

Eyewire: 3i

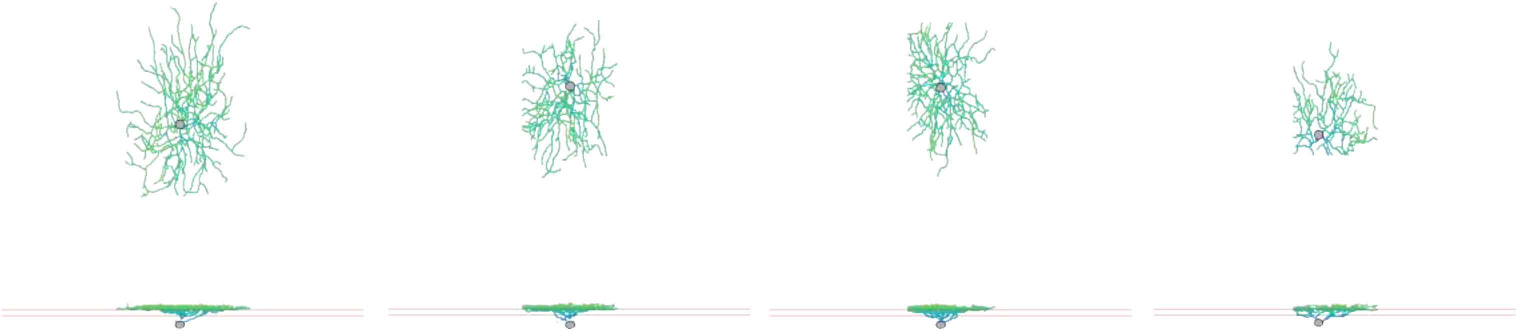

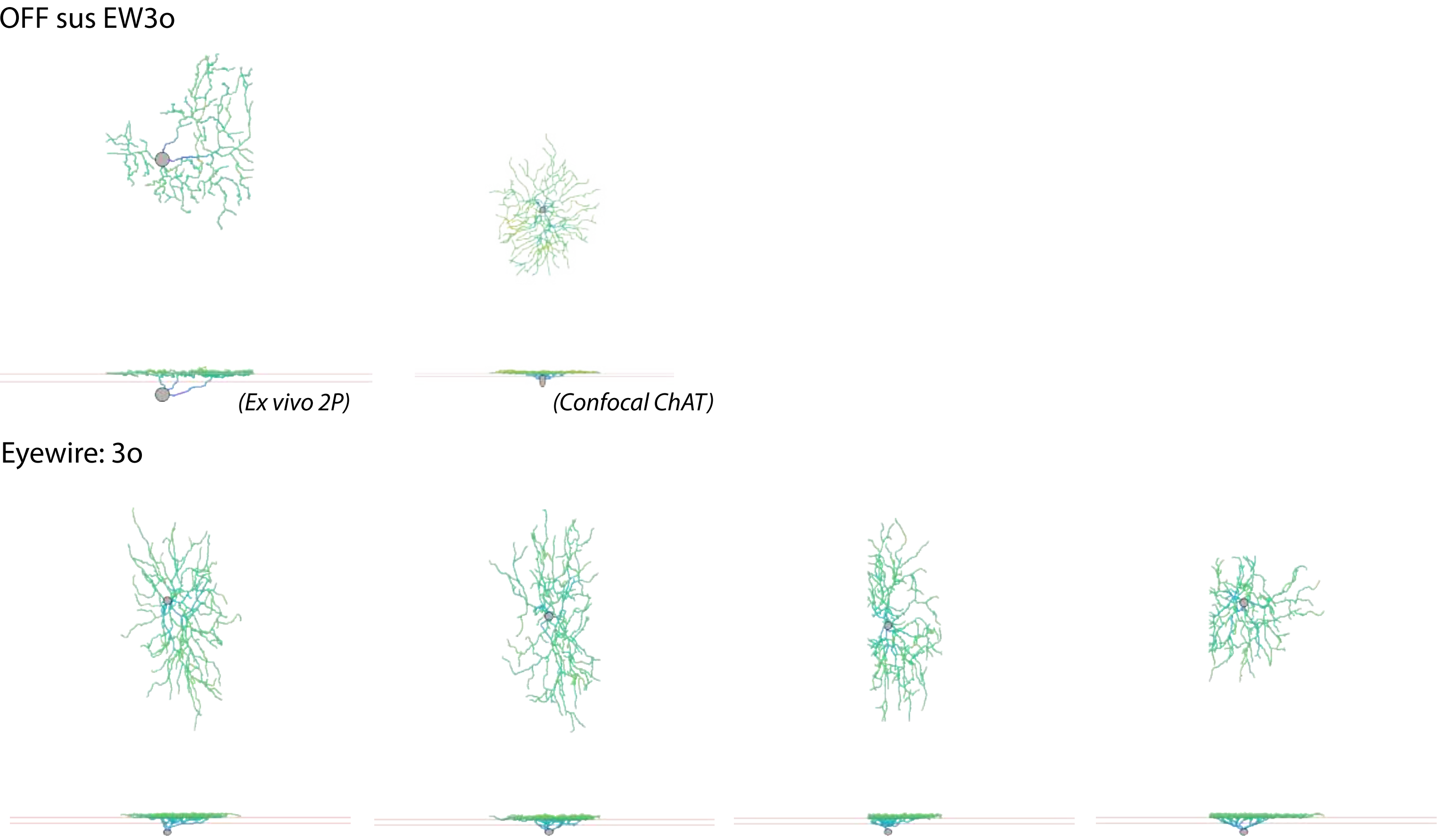
OFF sus EW3o
(Ex vivo 2P)
(Confocal ChAT)
Eyewire: 3o

OFF tr SmRF

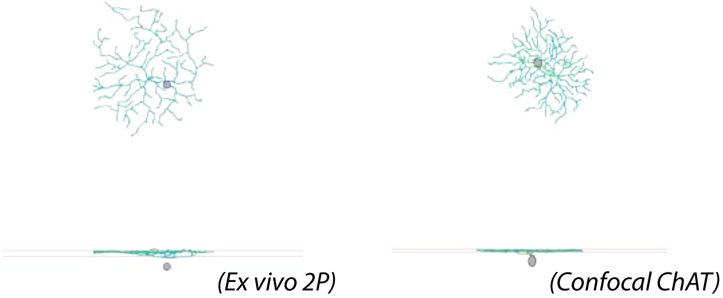


Eyewire: 4i

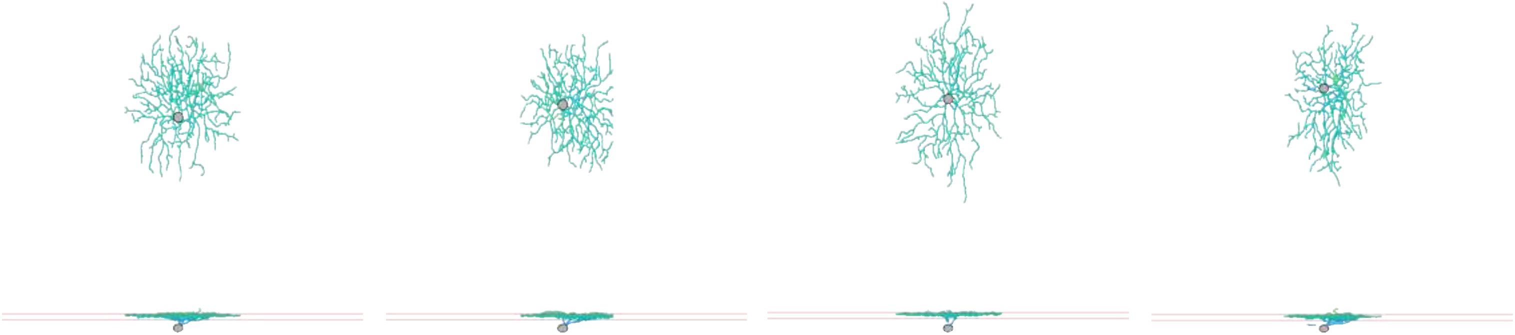

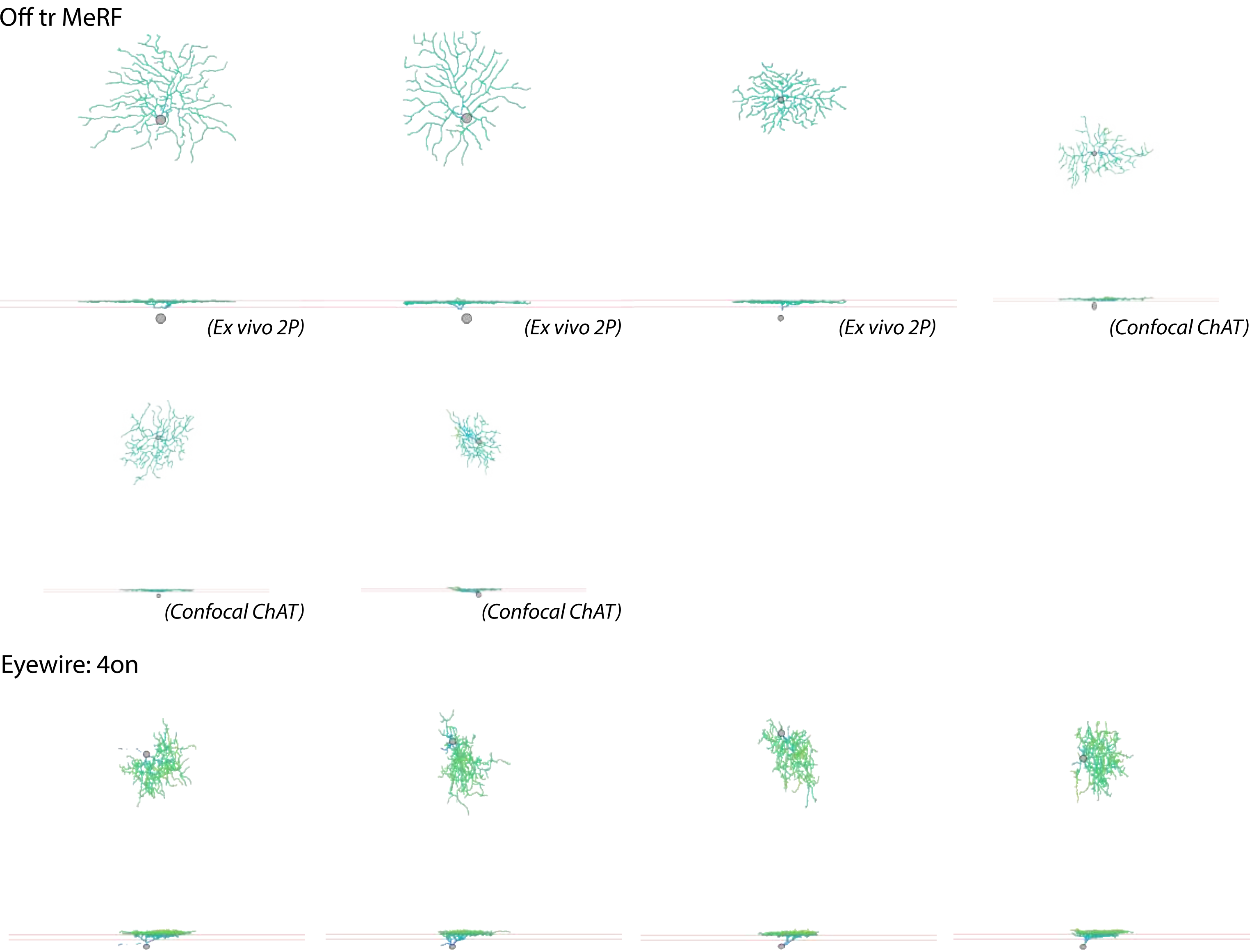
Off tr MeRF
(Ex vivo 2P)
(Ex vivo 2P)
(Ex vivo 2P)
(Confocal ChAT)
(Confocal ChAT)
(Confocal ChAT)
Eyewire: 4on

OFF tr alpha

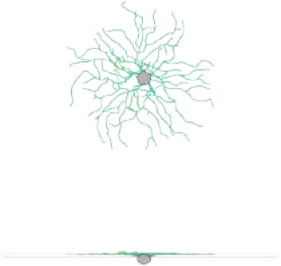

*(Confocal ChAT)*

Eyewire: 4ow

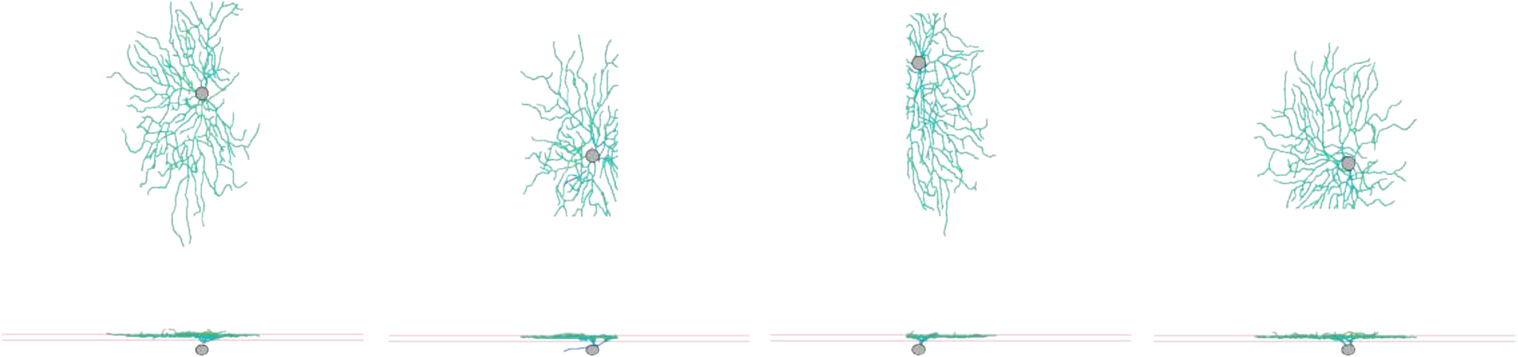

LED

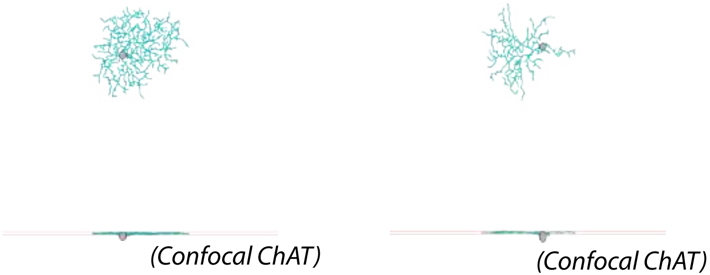


Eyewire: 51

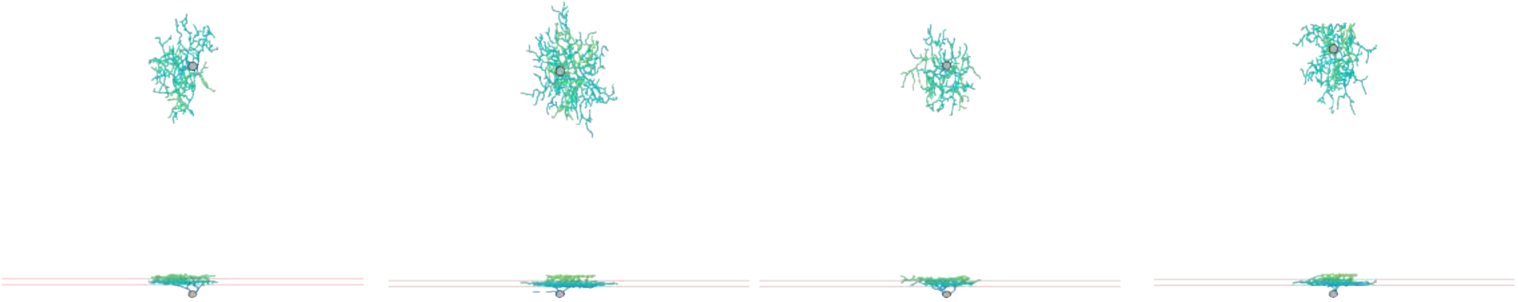

HD1

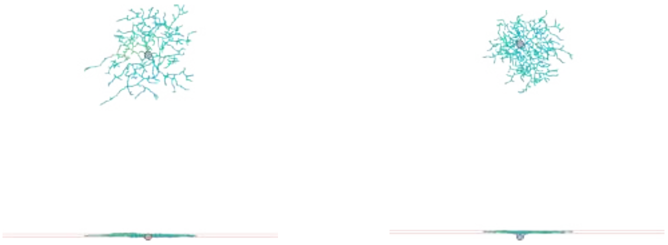

(Confocal ChAT) (Confocal ChAT)

Eyewire: 5si

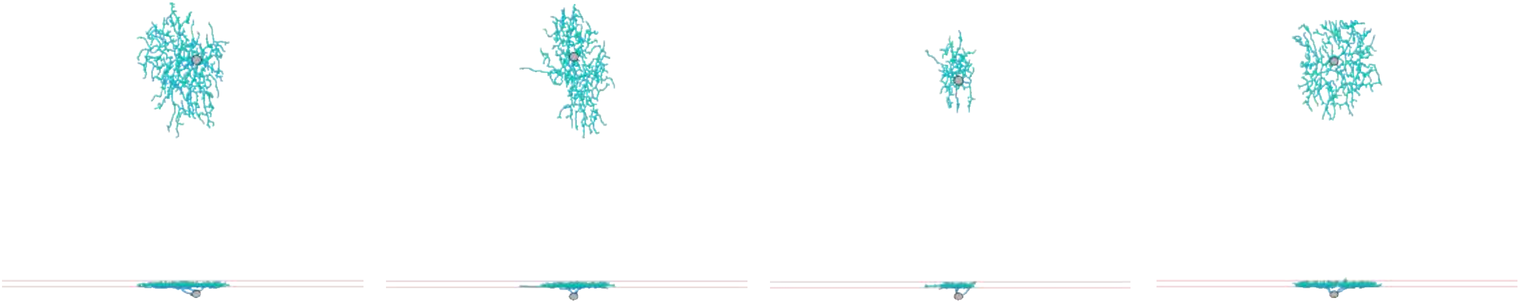

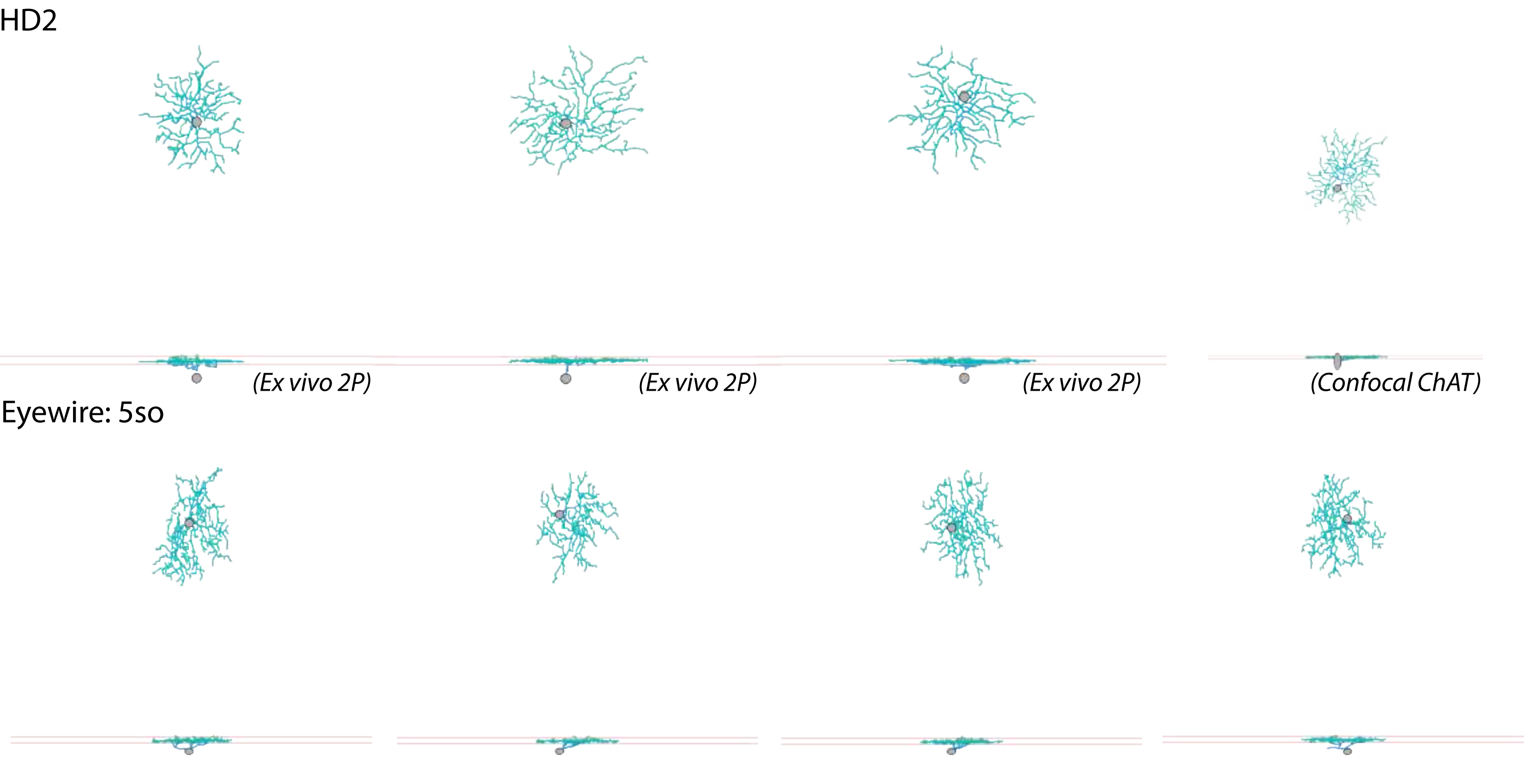
HD2
(Ex vivo 2P)
(Ex vivo 2P)
(Ex vivo 2P)
(Confocal ChAT)
Eyewire: 5so

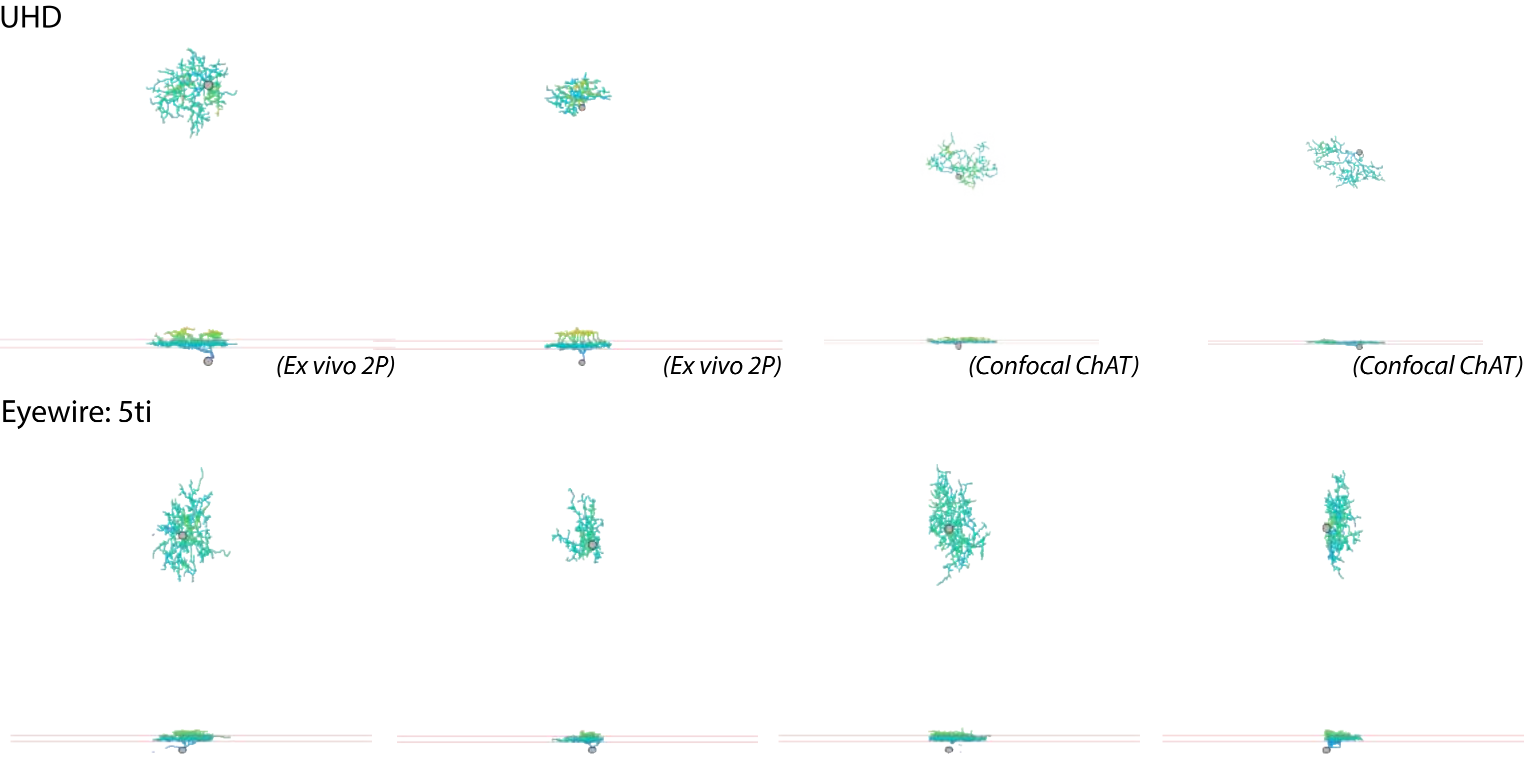
UHD
(Ex vivo 2P)
(Ex vivo 2P)
(Confocal ChAT)
(Confocal ChAT)
Eyewire: 5ti

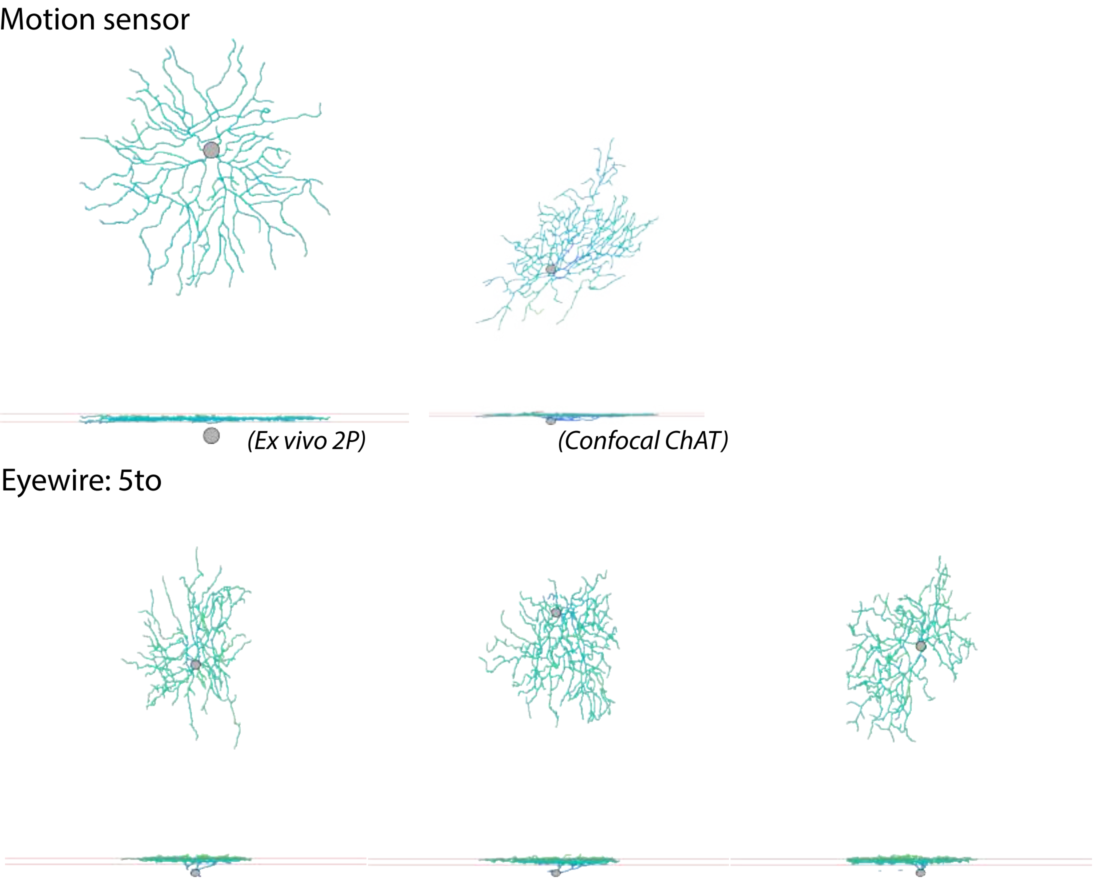
Motion sensor
(Ex vivo 2P)
(Confocal ChAT)
Eyewire: 5to

F-mini-ON

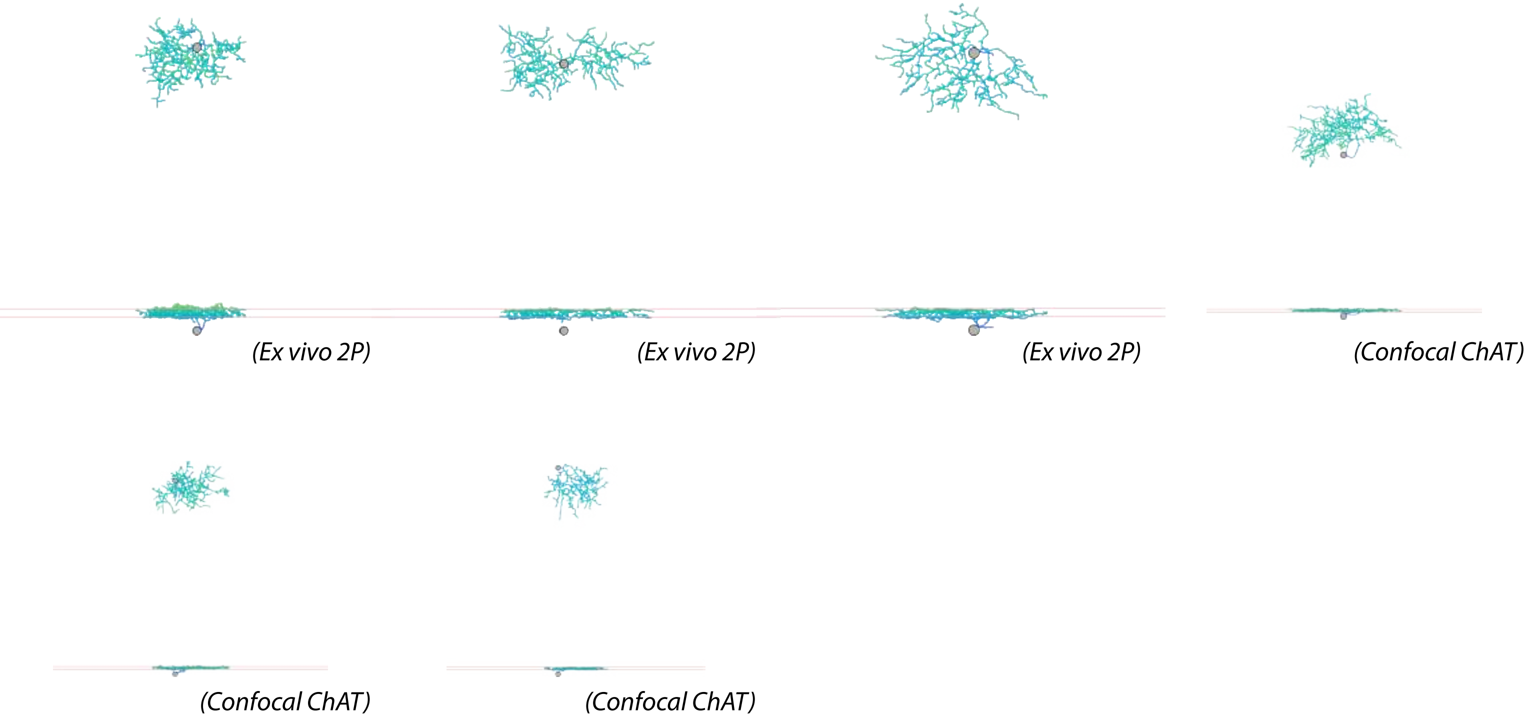


Eyewire: 63

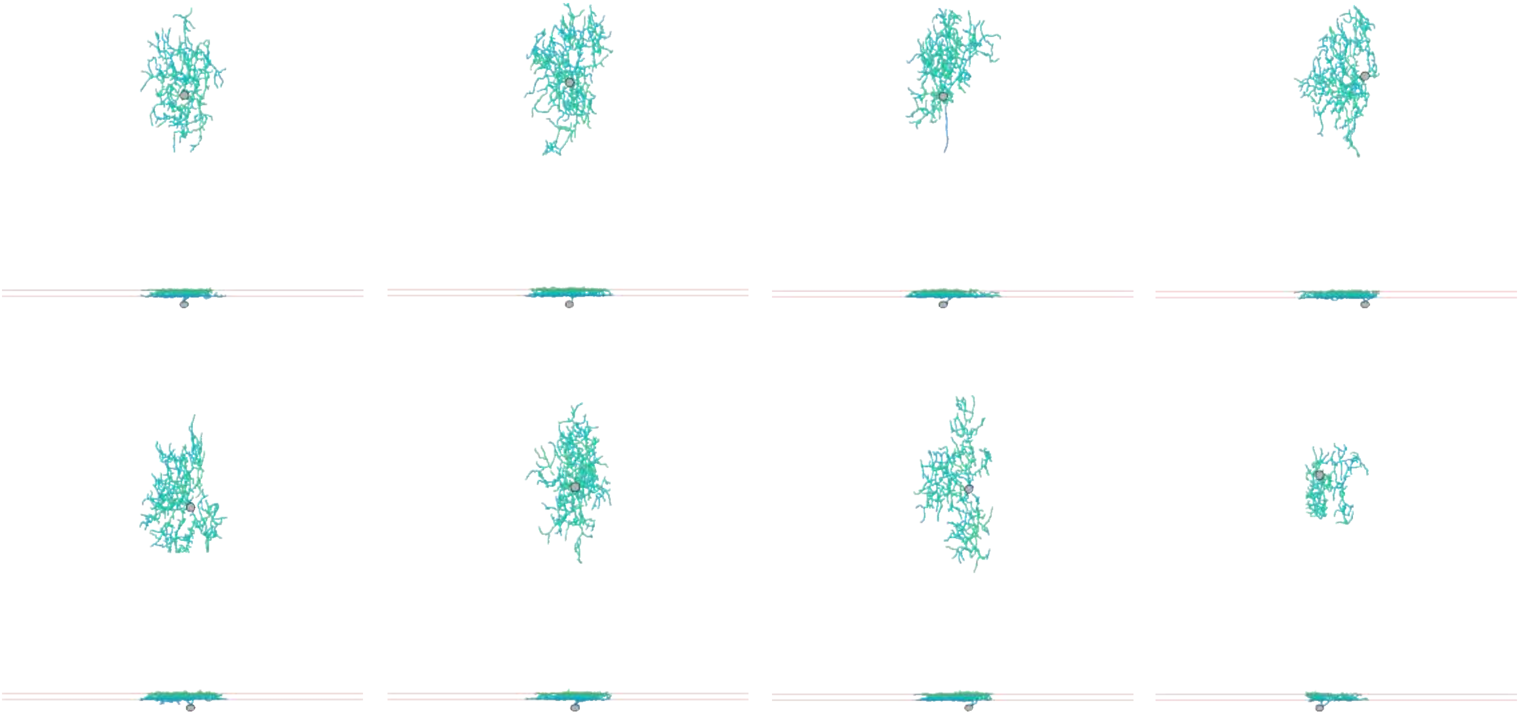

ON tr SmRF

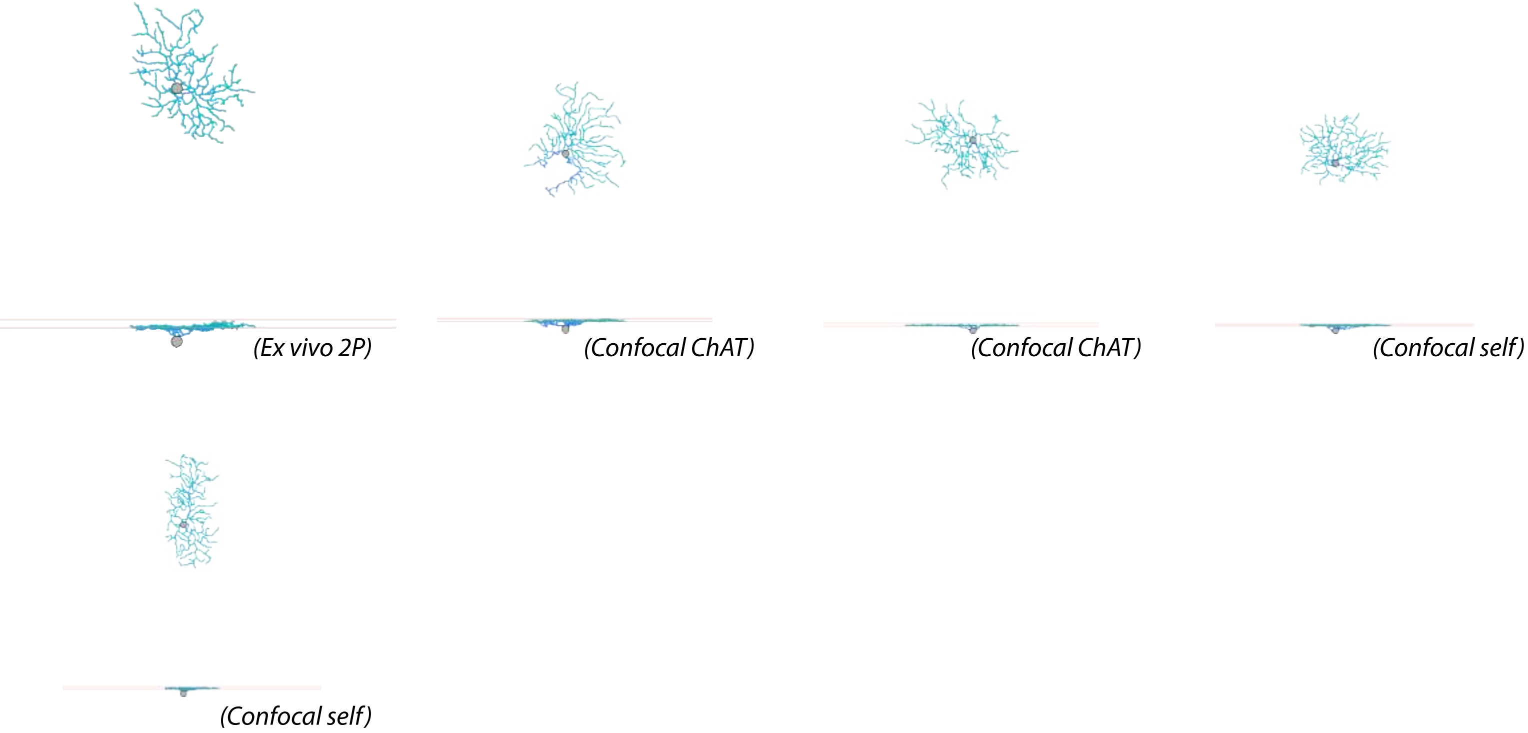


Eyewire: 6sn

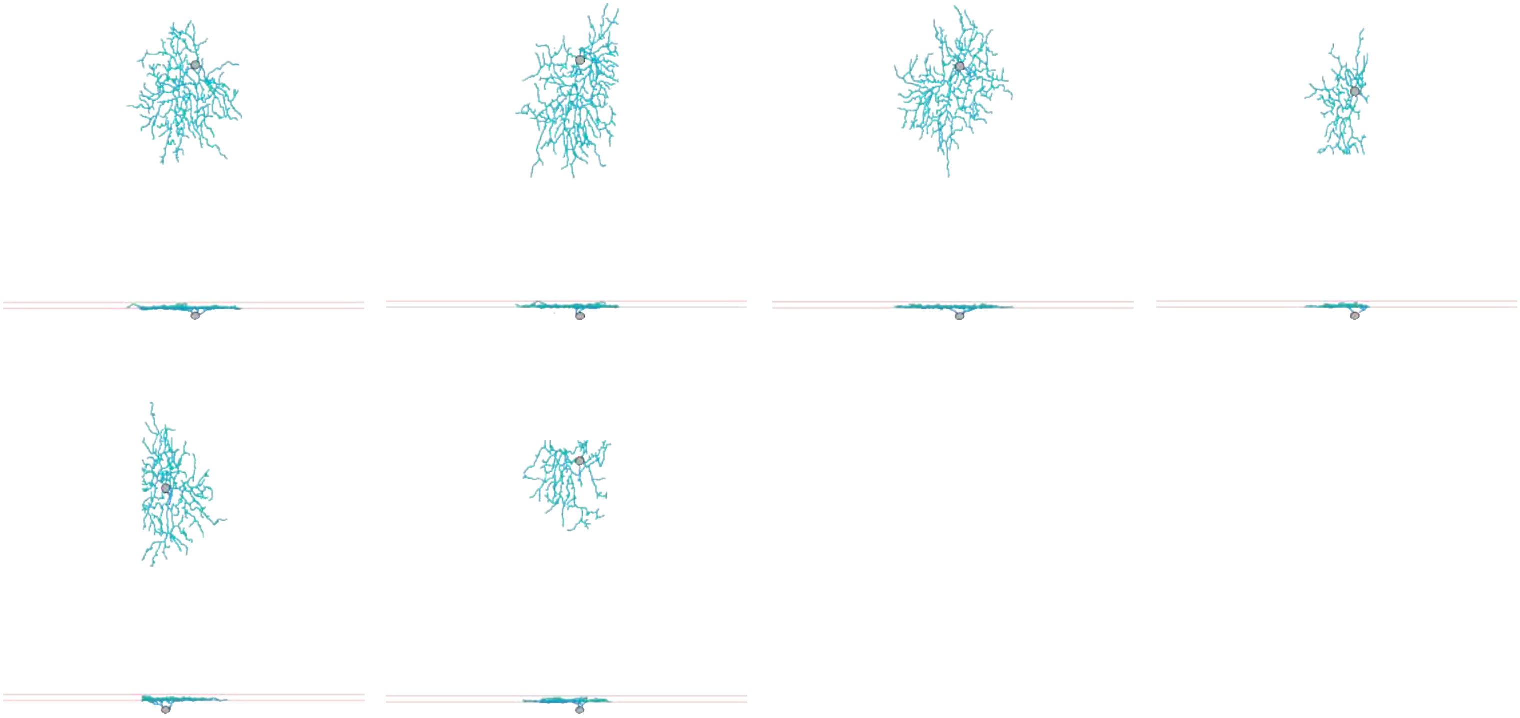

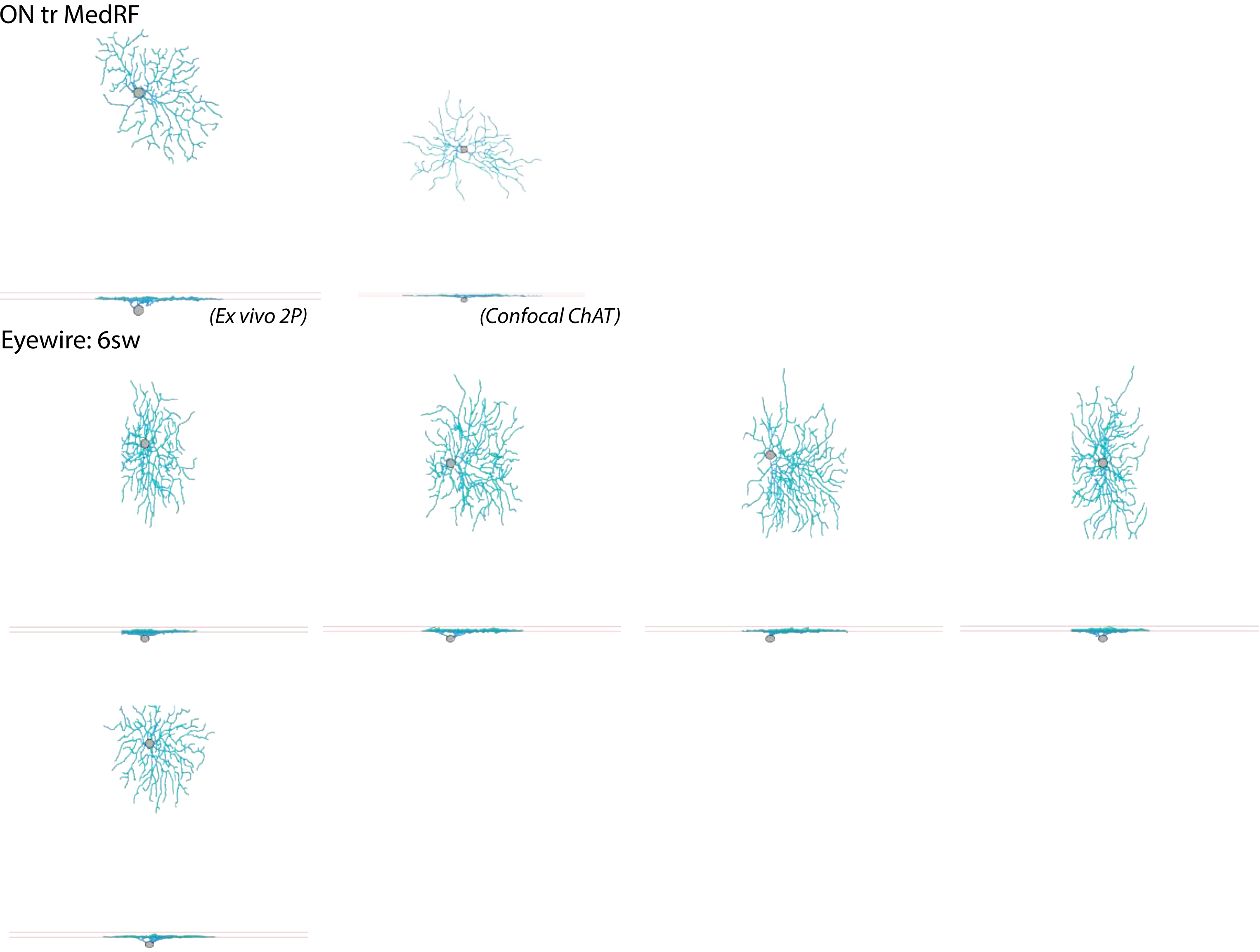
ON tr MedRF
(Ex vivo 2P)
(Confocal ChAT)
Eyewire: 6sw

ON tr EW6t

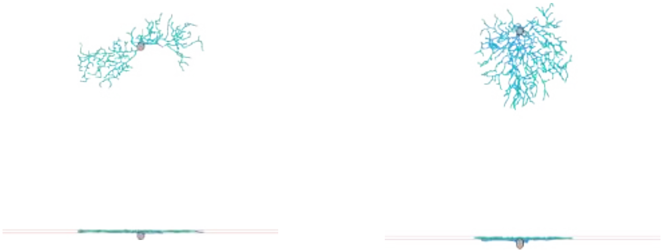

(Confocal ChAT) (Confocal ChAT)

Eyewire: 6t

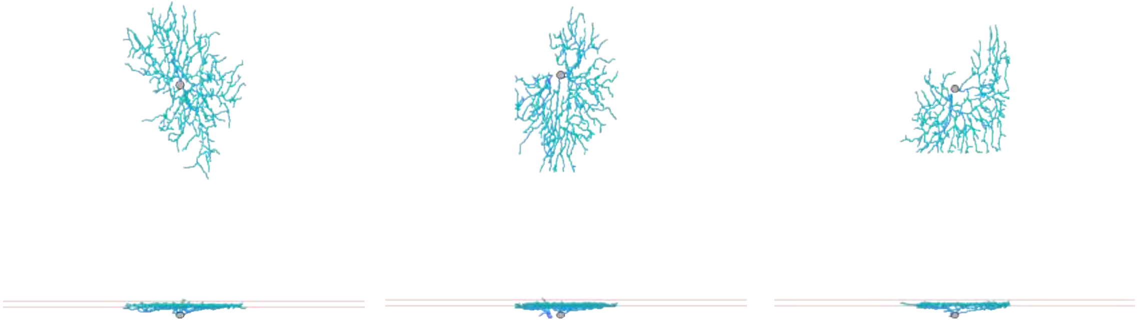

ON vOS SmRF

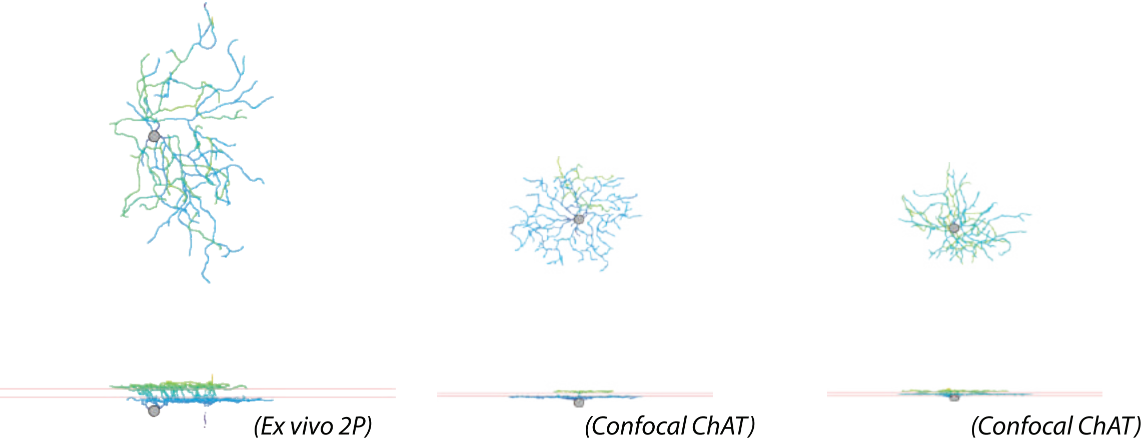


Eyewire: 72

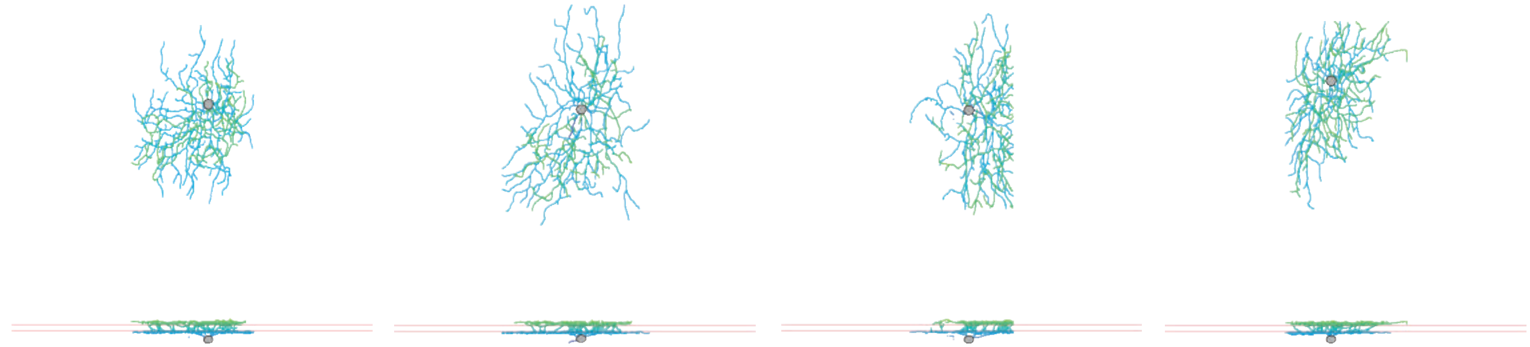

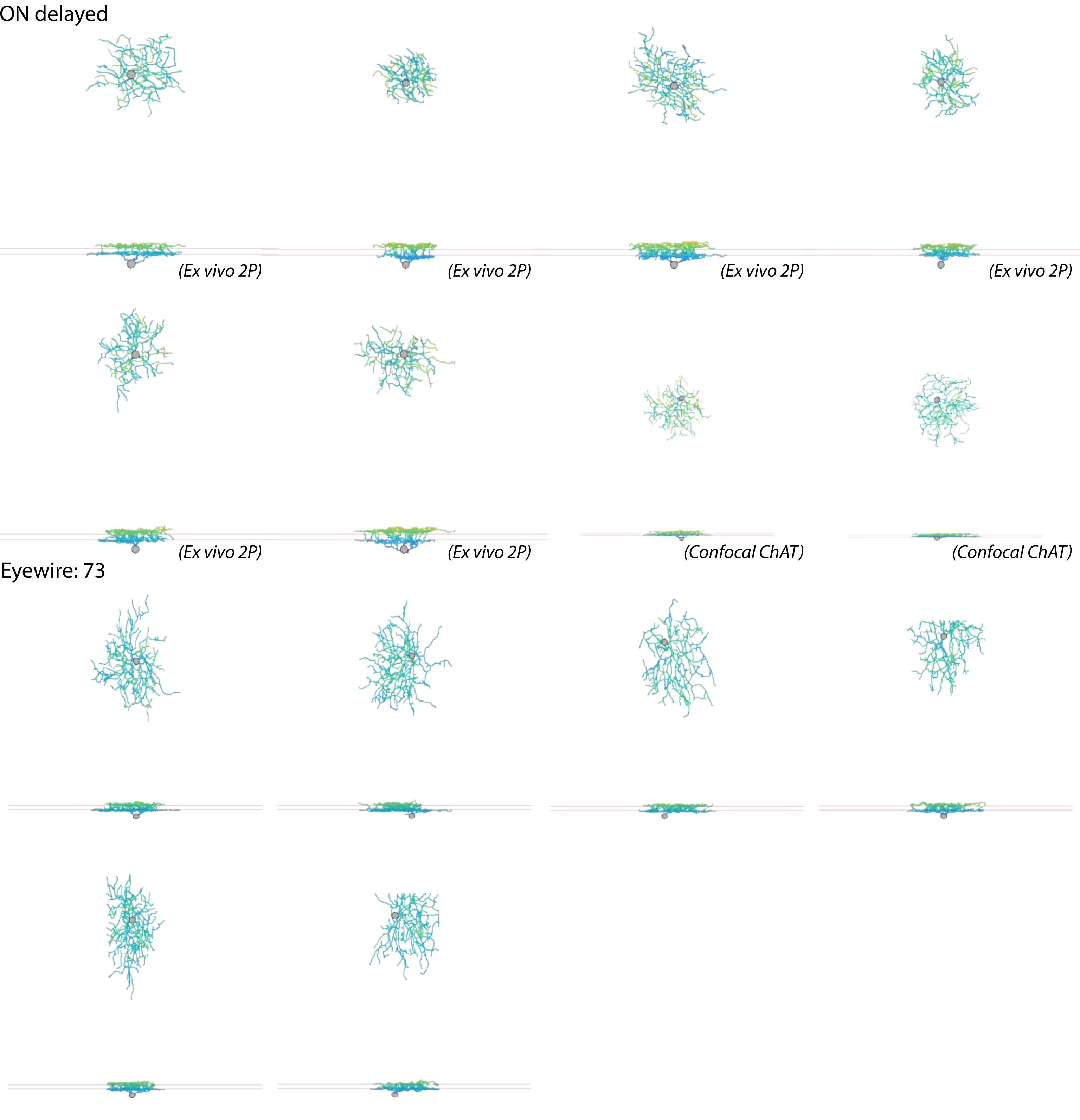
ON delayed
(Ex vivo 2P)
(Ex vivo 2P)
(Ex vivo 2P)
(Ex vivo 2P)
(Ex vivo 2P)
(Ex vivo 2P)
(Confocal ChAT)
(Confocal ChAT)
Eyewire: 73

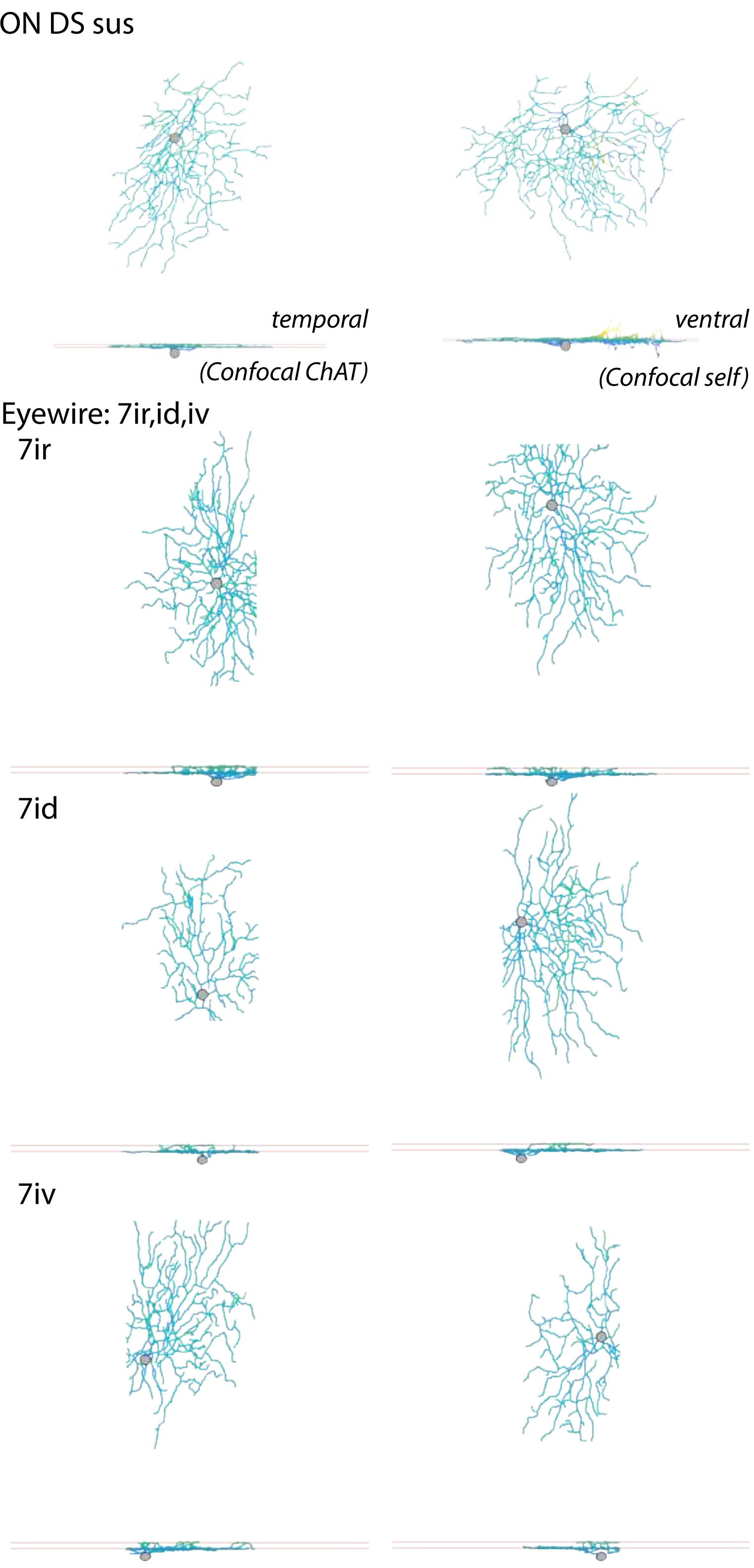
ON DS sus
temporal
(Confocal ChAT)
ventral
(Confocal self)
Eyewire: 7ir,id,iv
7ir
7id
7iv

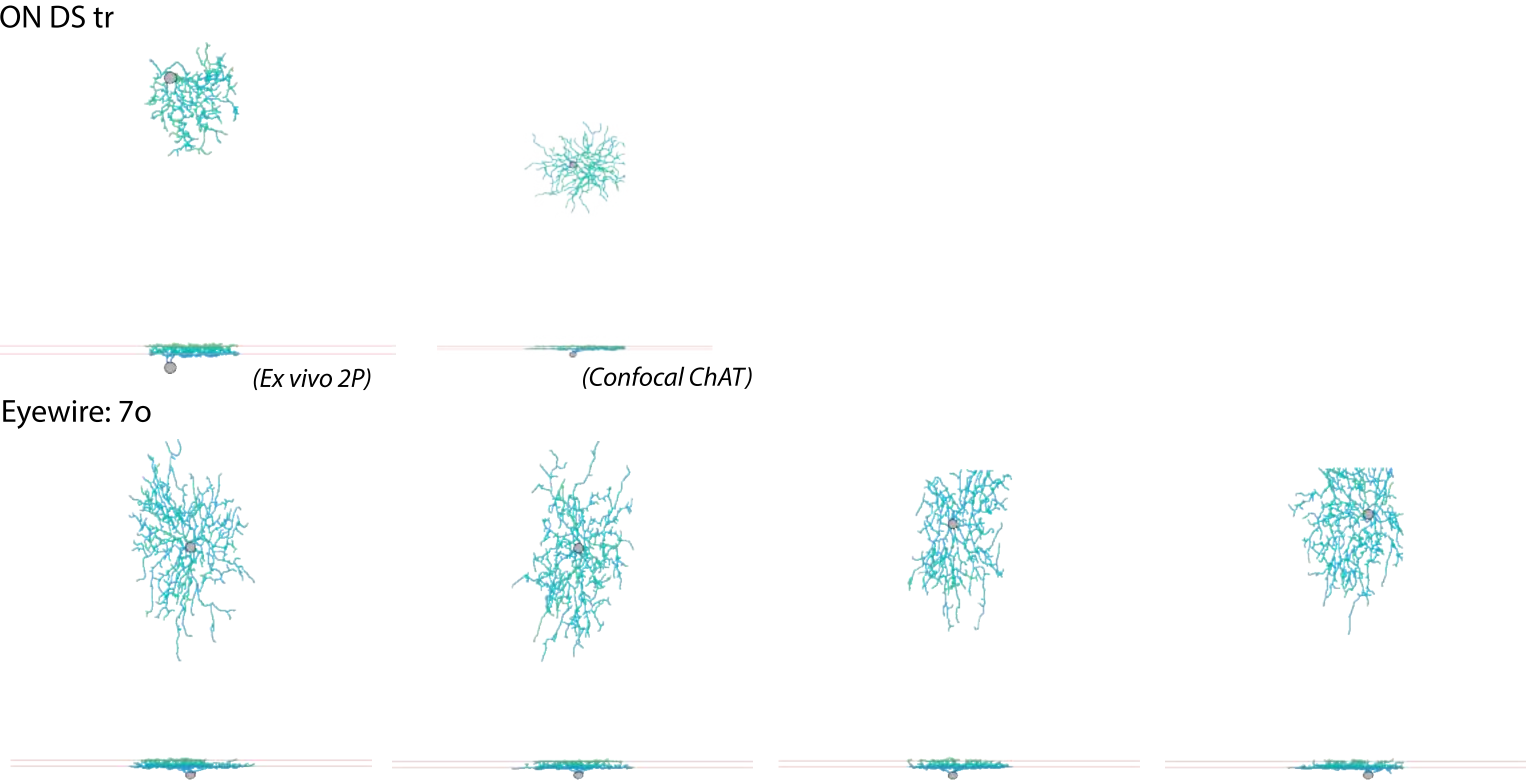

ON DS tr
(Ex vivo 2P)
(Confocal ChAT)
Eyewire: 7o

ONvOS LgRF

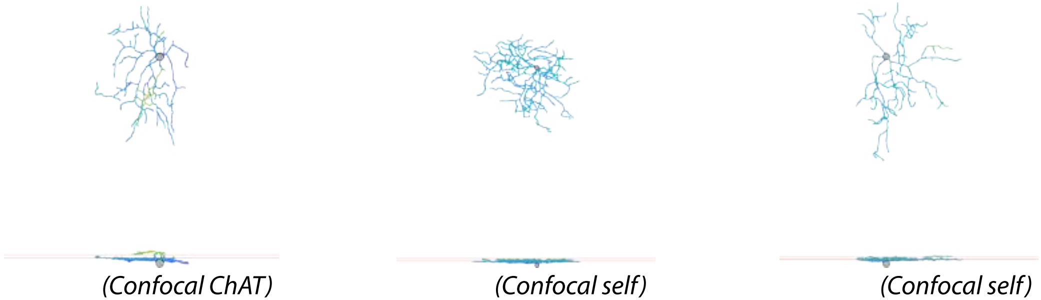


Eyewire: 81i,o

81i

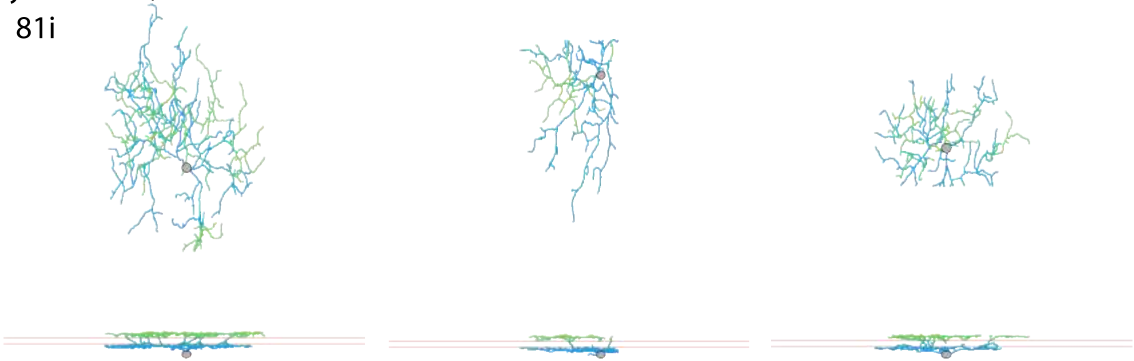

81o

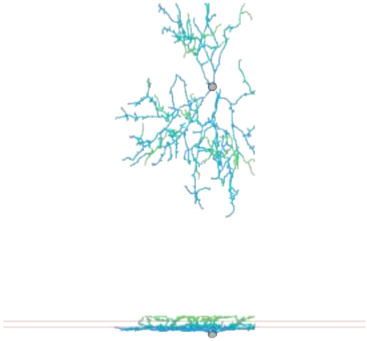

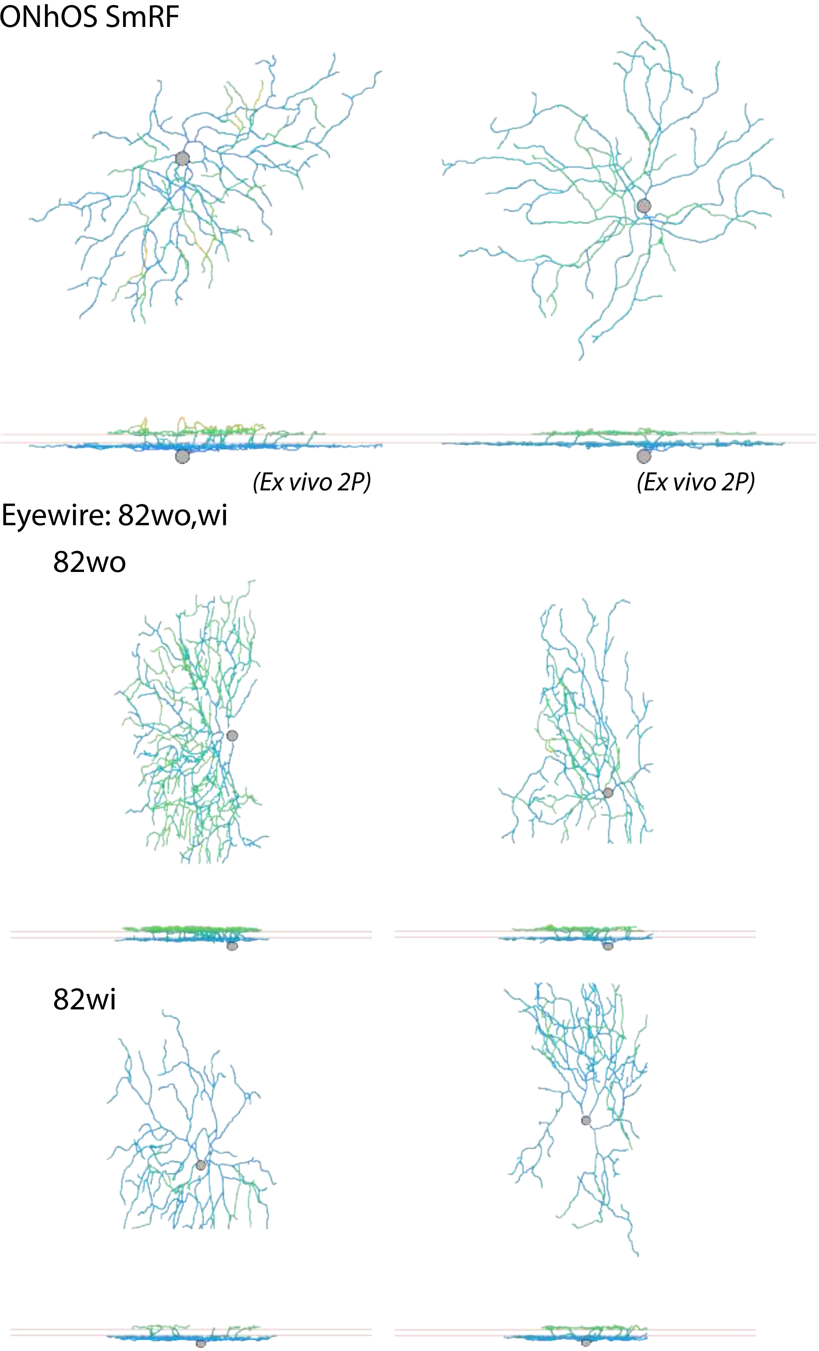
ONhOS SmRF
(Ex vivo 2P)
(Ex vivo 2P)
Eyewire: 82wo,wi
82wo
82wi

ONhOS LgRF

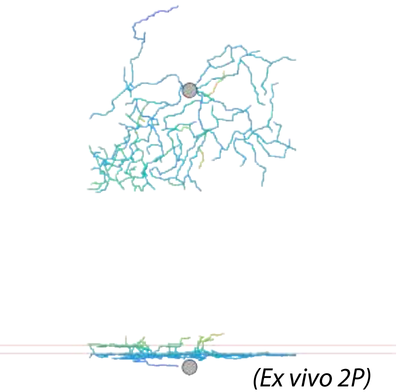


Eyewire: 82n

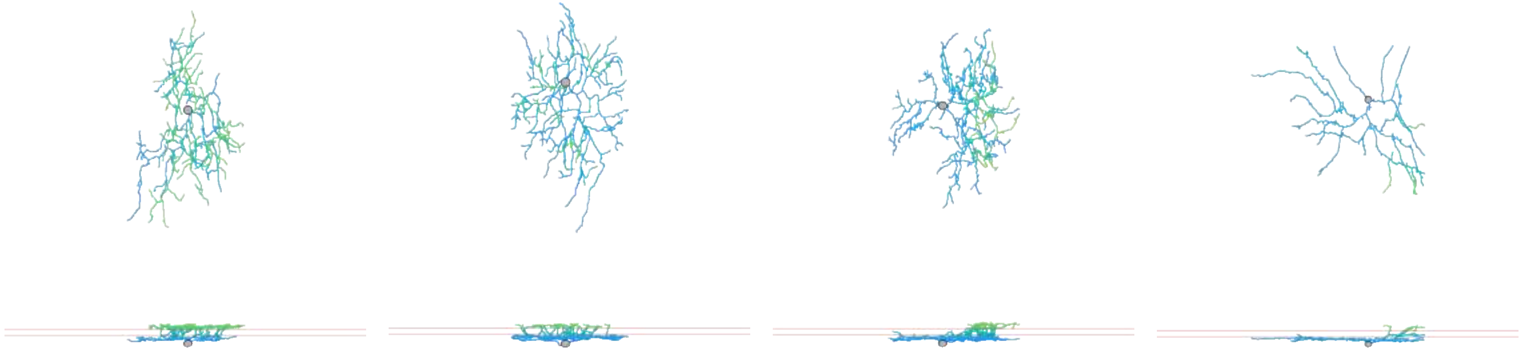

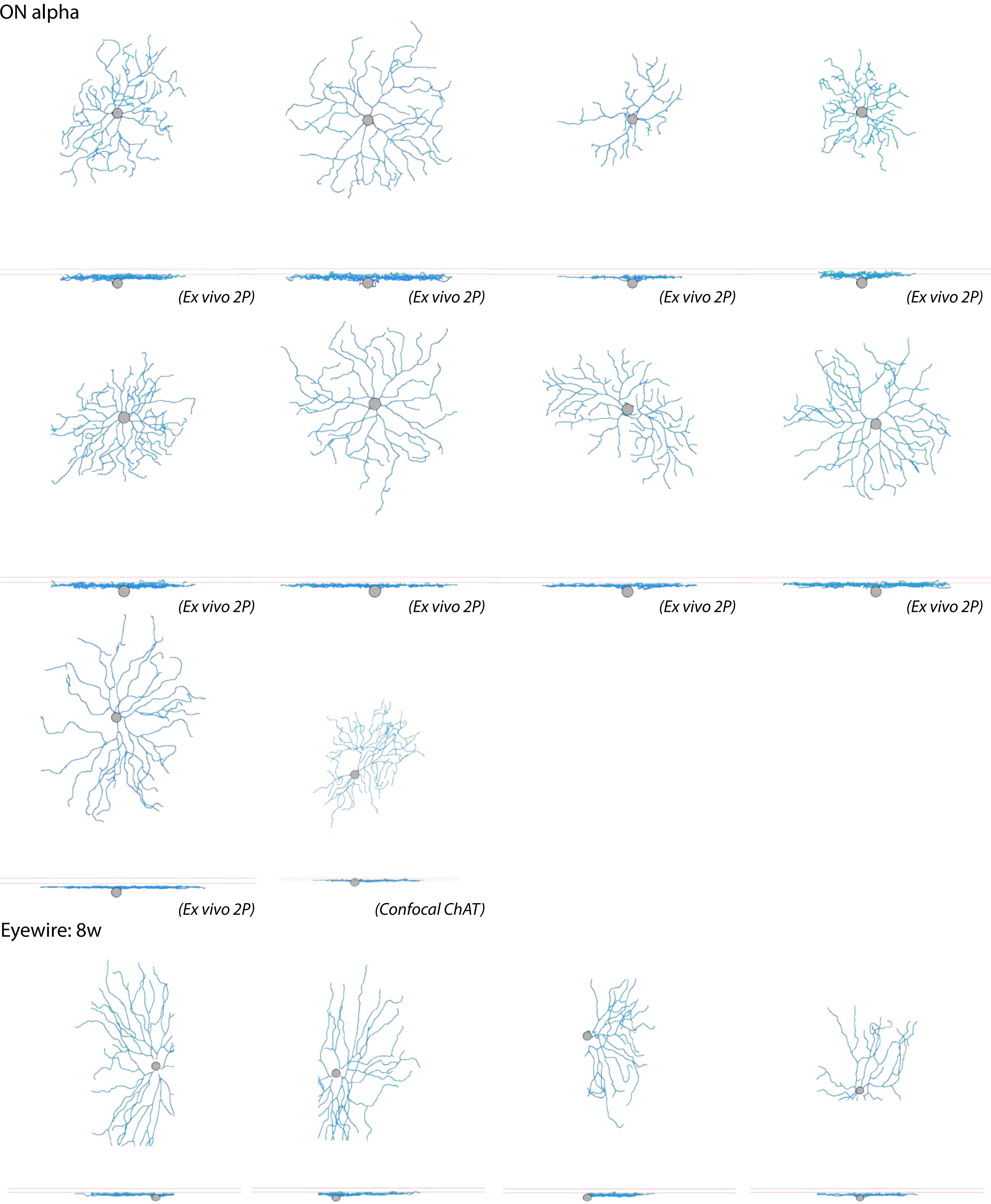
ON alpha
(Ex vivo 2P)
(Ex vivo 2P)
(Ex vivo 2P)
(Ex vivo 2P)
(Ex vivo 2P)
(Ex vivo 2P)
(Ex vivo 2P)
(Ex vivo 2P)
(Ex vivo 2P)
(Confocal ChAT)
Eyewire: 8w

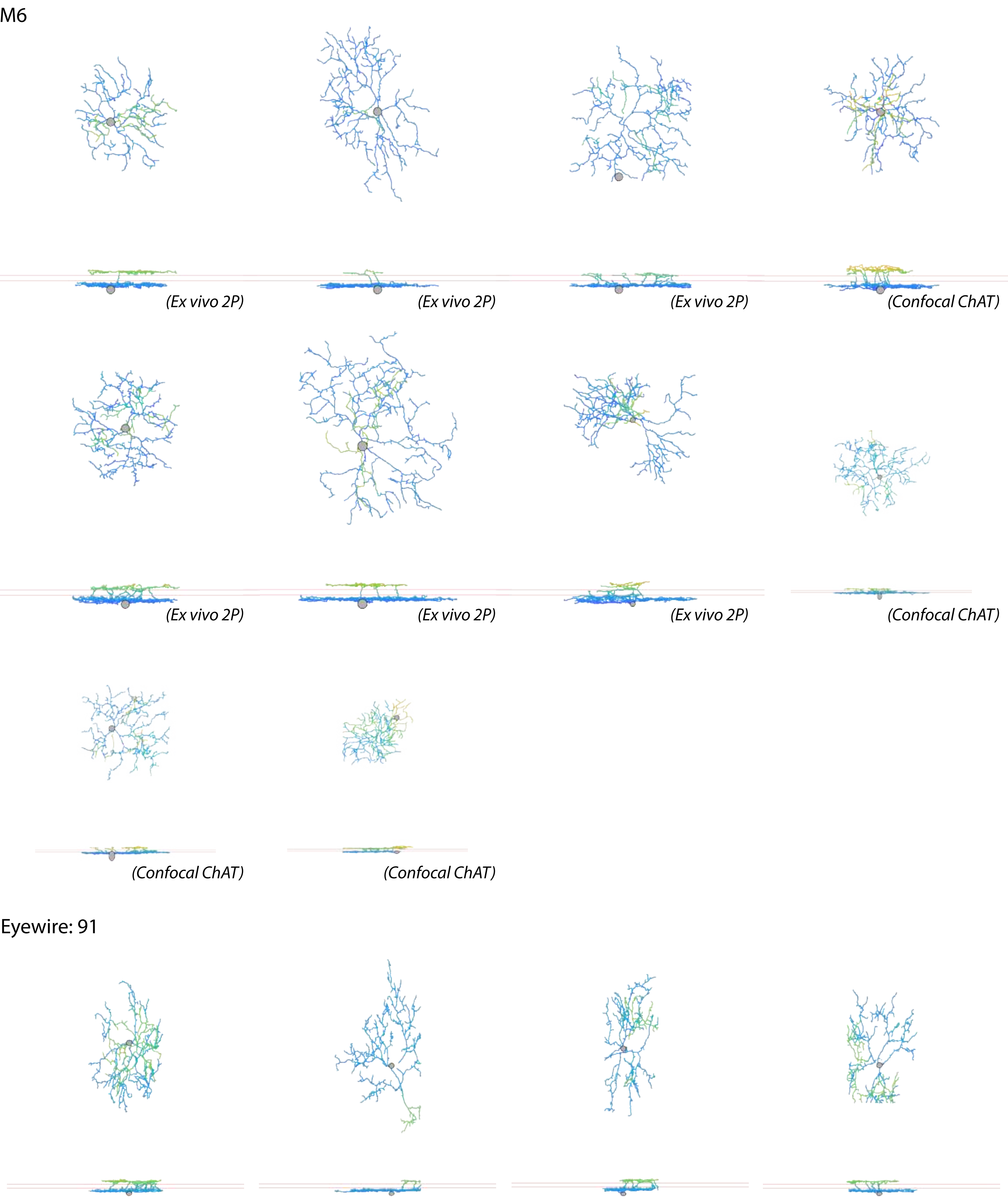
M6
(Ex vivo 2P)
(Ex vivo 2P)
(Ex vivo 2P)
(Confocal ChAT)
(Ex vivo 2P)
(Ex vivo 2P)
(Ex vivo 2P)
(Confocal ChAT)
(Confocal ChAT)
(Confocal ChAT)
Eyewire: 91

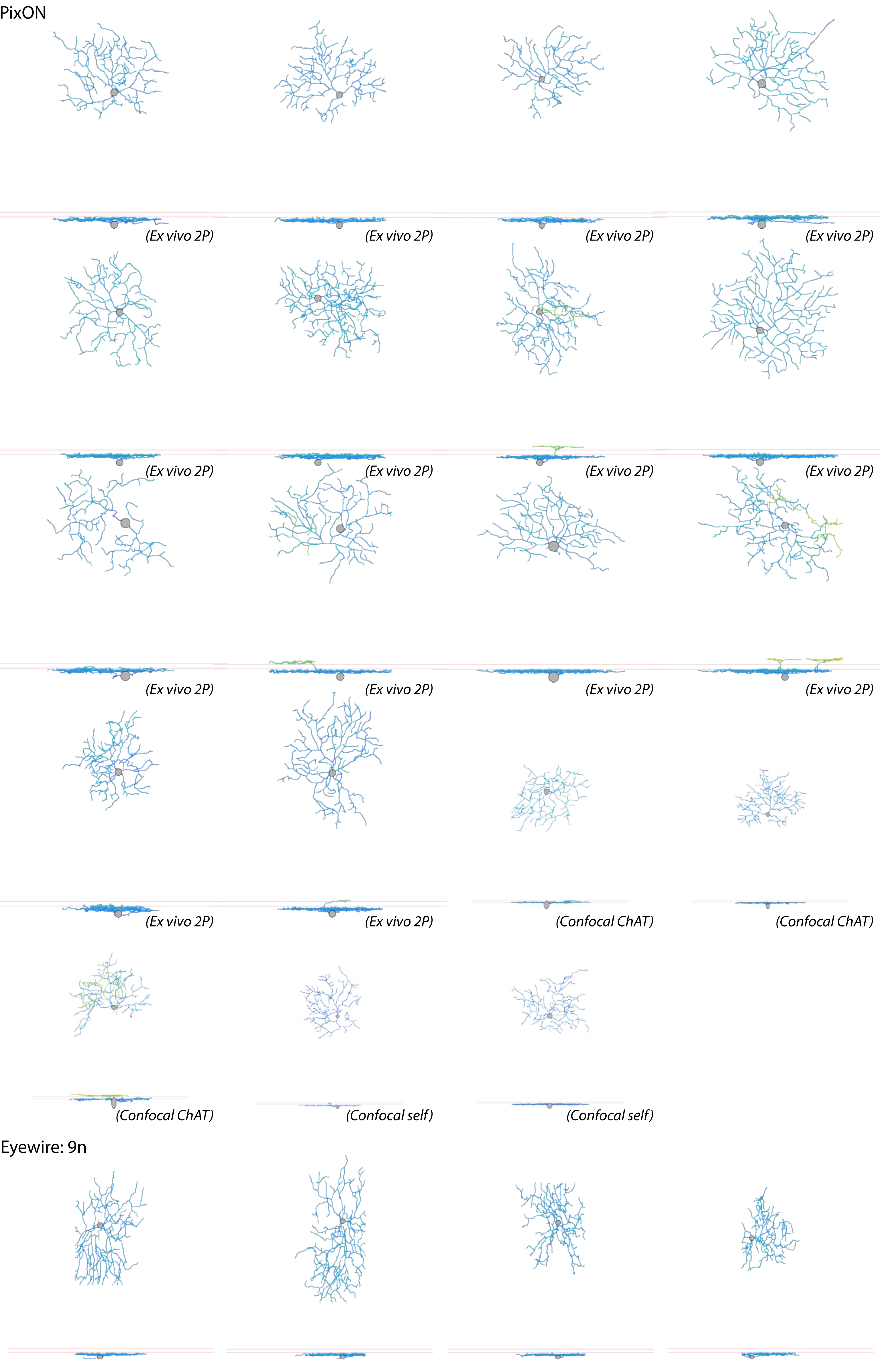

PixON
(Ex vivo 2P)
(Ex vivo 2P)
(Ex vivo 2P)
(Ex vivo 2P)
(Ex vivo 2P)
(Ex vivo 2P)
(Ex vivo 2P)
(Ex vivo 2P)
(Ex vivo 2P)
(Ex vivo 2P)
(Ex vivo 2P)
(Ex vivo 2P)
(Ex vivo 2P)
(Ex vivo 2P)
(Confocal ChAT)
(Confocal ChAT)
(Confocal ChAT)
(Confocal self)
(Confocal self)
Eyewire: 9n

M2

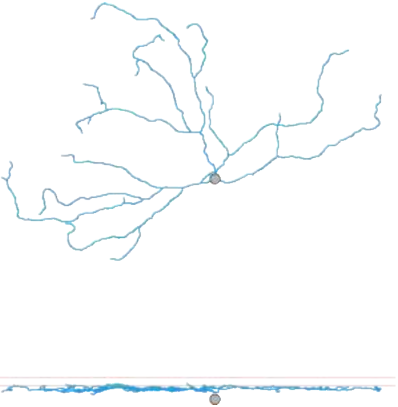

(Confocal ChAT)

Eyewire: 9w

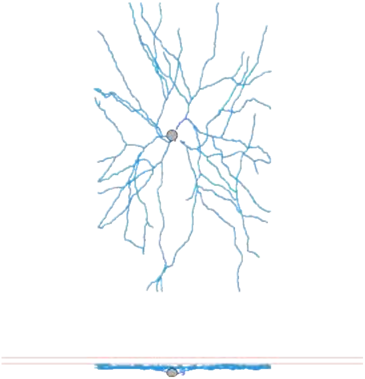
